# Serine metabolism changes link sphingolipid and one-carbon metabolism to cardiovascular risk factors in a cross-sectional study of asymptomatic adults

**DOI:** 10.64898/2026.09.20.753050

**Authors:** Mengjiang Huang, Debra K M Tacad, Tong Shen, Nancy L. Keim, Brian J. Bennett, John W. Newman

## Abstract

**BACKGROUND:** Metabolic perturbations associated with cardiovascular disease (CVD) risk factors prior to clinical disease manifestation may inform novel preventative measures. Here we characterize metabolome-wide associations with various cardiometabolic risk factors in an asymptomatic population to develop a mechanistic metabolic framework of preemergent metabolic shifts.

**METHODS:** Plasma was collected from a clinically healthy cross-sectional cohort (n = 361). Cohort mass spectrometry-based metabolomics (biocrates MxP^®^ Quant 500 kits) were complemented with expanded lipidomic coverage for a sex/age-matched subset with LDLc difference > 60 mg/dL (n = 38). LDLc-focused partial least square regressions and discriminate analyses were performed, after false discovery rate corrections.

**RESULTS:** As LDLc levels increased, plasma sphingolipids and cholesteryl esters, and transulfuration products were higher, while ether-linked phosphatidylcholines serine, glycine, alanine, methionine were lower. The subset analysis further highlighted elevations in triglycerides and deoxyceramides with elevated LDLc. When adjusted for alanine concentrations, deoxyceramide concentrations increased as serine declined (p = 0.0012), and the deoxyceramide:ceramide ratios were higher in the high LDLc group (p = 0.039). Lower serine levels were also accompanied by parallel changes in glycine (p= 0.0004), and anti-parallel changes in transsulfuration pathway metabolites particularly when adjusted for ceramide levels. Notably, the alanine:serine ratio was more strongly associated with fasting triglycerides (r = 0.38; p <0.0001) than LDLc.

**CONCLUSIONS:** Serine depletion by increased demands of sphingolipid, and possibly transsulfuration pathway, metabolism is associated with elevations in cytotoxic deoxyceramides and reductions in glycine and methionine availability for one-carbon metabolism. These changes were strongly associated with fasting triglycerides and insulin resistance, key markers of residual CVD risk. The parent study is registered at *<u>ClinicalTrials.gov</u>* <u>NCT02367287.</u>

**Significance Statement:** This study identifies serine as a central metabolic node linking cardiovascular risk factors, sphingolipid remodeling and one-carbon metabolism in clinically healthy adults. We report data consistent with an increased demand for ceramide synthesis leading to serine depletion and accumulation of cytotoxic deoxyceramides, providing a mechanistic metabolic framework explaining metabolomic shifts in asymptomatic adults. These findings support the targeting of serine-dependent pathways for managing residual cardiovascular risk.

## Introduction

Atherosclerotic cardiovascular disease is initiated and propagated by the retention, modification, and inflammatory remodeling of cholesterol-rich lipoproteins within the arterial wall, and low-density lipoprotein cholesterol (LDLc) remains a central causal risk factor and primary therapeutic target for atherosclerosis prevention.^1,2^ The seminal works of Keyes on dietary fats, and of Brown and Goldstein on genetics of familial hypercholesterolemia helped to establish LDL cholesterol (LDLc) as an independent risk factor for atherosclerosis leading to its widespread use as both a clinical biomarker and therapeutic target.^3,4^ Accordingly, plasma LDLc remains the routine biomarker and primary therapeutic target for atherosclerosis prevention.

Mounting evidence indicates that lipid alterations beyond cholesterol track cardiovascular disease progression ^5,6^. Modern lipidomics allows for the mapping of hundreds of glycolipids, glycerophospholipids, sphingolipids, cholesterols, and oxidized derivatives of lipid species that conventional lipid panels cannot.^7^ Moreover broad lipidomic changes that alter intracellular signaling and lipoprotein functions are associated with risks from cardiovascular death.^8^ Notably, sphingolipid changes track both atherosclerotic burden and cardiovascular disease risks, and ceramide-based risk scores (e.g., CERT1/2) have improved predictive performance of incidence over LDLc alone. ^9–11^

Pathways beyond lipid metabolism have also been implicated in atherosclerosis. For instance, one-carbon metabolism, which includes the folate and methionine cycles, provides methyl units that support many homeostatic processes including DNA/RNA biosynthesis, epigenetic regulation, and cellular oxidative stress responses ^12^. Disruptions in this network can promote oxidative stress and endothelial dysfunction, major steps in atherosclerosis progression.^13,14^ In particular, one-carbon metabolism and sphingolipid metabolism share the cellular serine pool. While these pathways have largely been studied in isolation, they may be influenced in a coordinated fashion during cardiometabolic disease progression.

In the current study, we integrated targeted plasma metabolomics with expanded lipidomic profiling of a cross-sectional cohort of men and women balanced for age and body mass index, who were clinically asymptomatic for cardiometabolic diseases including cardiovascular disease, hyperlipidemia or diabetes.^15^ This is in contrast to most metabolomic and lipidomic studies of cardiovascular disease which have focused on secondary prevention cohorts, comparing healthy controls with patients presenting clinical manifestation such as myocardial infarction or coronary artery disease.^16,17^ The goal of this effort was to identify system-level patterns associated with early established cardiovascular risk factors. We hypothesized that impacts on serine metabolism would be an early and unifying metabolic defect linking major pathways associated with cardiometabolic disease.

## Materials and Methods

### 2.1 Study participants

Participants enrolled in the USDA Western Human Nutrition Research Center (WHNRC) Cross-Sectional Nutritional Phenotyping Study (*ClinicalTrials.gov*: NCT02367287) conducted between 2015 and 2019 in Davis, CA provided samples after written informed consent as authorized under University of California Institutional Review Board identification number 691654.^15^ This parent study recruited 393 healthy male and female participants between age of 18 and 65 with BMIs between 18.5 to 40 kg/m^2^ into six sex x age x BMI bins to investigate responses to nutritional and psychosocial stresses (**Figure 1**). The population ethnic distribution were similar to the 2010 CA census: 61.1% (versus 60.6%), White; 15.4% (versus 13.7%), other race; 12.3% (versus 14.1%), Asian; 4.8% (versus 5.8%), Black; 4.8% (versus 4.7%), two or more races; 0.6% (versus 0.8%), native American; 1.1% (versus 0.4%), native Hawaiian or Pacific Islander. Detailed eligibility criteria and study procedures are published elsewhere.^15^ Participants were required to fast 12h after the consumption of a standardized meal before the clinical study visit. Participants were broadly phenotyped with assessments of clinical lipids, insulin sensitivity, and vascular function as previously reported. Here, fasting LDLc concentrations for all 393 study participants were screened as an indicator of atherosclerosis risk. Participants were also classified as low-risk (LDLc < 100 mg/dL) or high-risk (LDLc ≥ 160 mg/dL) according to current guidelines ^18^. The analysis selection criteria and fasting LDLc distribution are presented in **Table 1** and **Supplemental Table ST1**.

**Figure 1:**
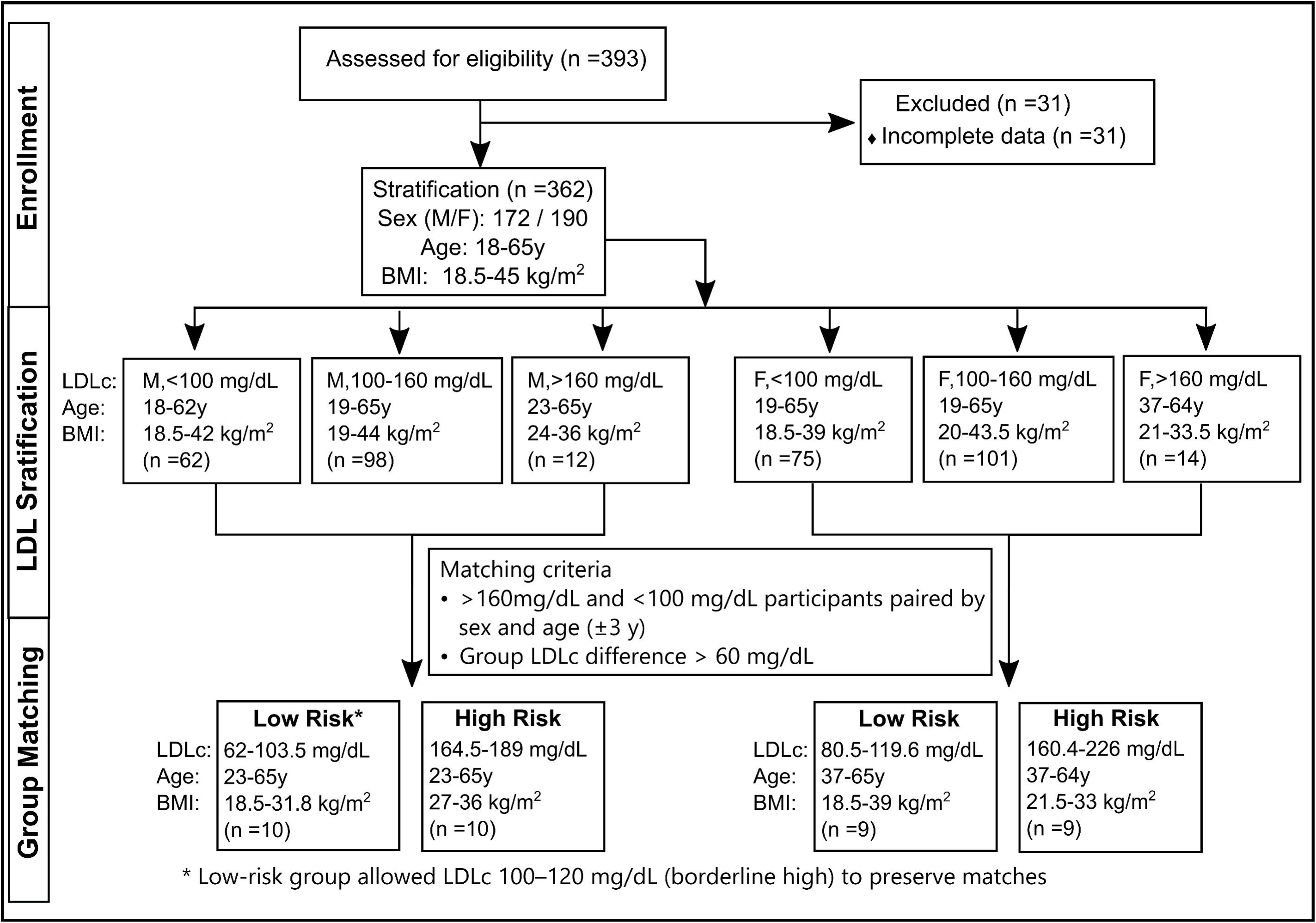
Participant data use flow diagram. The WHNRC nutritional phenotyping study enrolled 393 male (M) and (F) participants with attempts to balance into 3 age and BMI ranges. Of these, 361 were available for fasting metabolomic analyses. An age-x sex-matched subset of males (n = 10) and females (n = 9) were stratified by fasting LDLc level for targeted lipidomic analysis.

**Table 1:** Participant characteristics stratified by LDLc risk group and sex*^a^*.

|  |  | Sex |  | LDL Cholesterol Percentile |  |  |
| --- | --- | --- | --- | --- | --- | --- |
|  |  | Male<br>(n = 184) | Female<br>(n = 209) | 10<sup>th</sup><br>(n = 36) | 90<sup>th</sup><br>(n = 36) | P |
| Age | y | 39.7 ± 1.03 | 40.6 ± 0.95 | 30.9 ± 2.26 | 49.6 ± 2.2 | <.0001 |
| BMI | kg/m<sup>2</sup> | 27.1 ± 0.345 | 27.8 ± 0.33 | 26.8 ± 0.852 | 28.5 ± 0.84 | 0.13 |
| Cholesterol Homeostasis |  |  |  |  |  |  |
| Total-c | mg/dL | 172 ± 3.91 | 178 ± 4.2 | 124 ± 9.38 | 237 ± 9.3 | <.0001 |
| LDLc | mg/dL | 112 ± 2.99 | 109 ± 3 | 62 ± 6.26 | 171 ± 6.2 | <.0001 |
| HDLc | mg/dL | 49.5 ± 1.27 | 60.3 ± 1.5 | 55.2 ± 3.79 | 50.3 ± 3.7 | 0.10 |
| Rem-c | mg/dL | 10.7 ± 0.761 | 8.85 ± 0.73 | 7.31 ± 1.56 | 15.9 ± 1.5 | 0.0085 |
| Triglyceride Homeostasis |  |  |  |  |  |  |
| Fasting TGs | mg/dL | 101 ± 3.89 | 94.1 ± 3.7 | 81.2 ± 7.98 | 140 ± 7.9 | 0.0001 |
| High Fasting TG | % >1.5 mg/mL | 10% | 10% | 6% | 38% |  |
| High PPTG | % >200mg/dL | 41% | 26% | 19% | 70% |  |
| Glucose Homeostasis |  |  |  |  |  |  |
| Fasting Glucose | mg/dL | 96.1 ± 1.88 | 93.5 ± 1.9 | 91.3 ± 4.81 | 105 ± 4.7 | 0.039 |
| HOMA-IR |  | 2.26 ± 0.214 | 2.14 ± 0.11 | 1.89 ± 0.396 | 2.74 ± 0.39 | 0.048 |
| QUICKI |  | 3.35 ± 0.0824 | 3.25 ± 0.094 | 3.7 ± 0.187 | 3.13 ± 0.18 | 0.13 |
| McAuley-ISI |  | 6.87 ± 0.335 | 6.61 ± 0.3 | 9.18 ± 0.663 | 4.58 ± 0.65 | 0.0007 |
| Fasting NEFA | mEq/L | 0.29 ± 0.0101 | 0.346 ± 0.011 | 0.323 ± 0.0263 | 0.374 ± 0.026 | 0.15 |
| HOMA-IR status | % IR | 29% | 34% | 22% | 51% |  |
| Integrated Cardiometabolic Risk Factors |  |  |  |  |  |  |
| ASCVD Lifetime |  |  |  |  |  |  |
| Risk | Lifetime % | 31.3 ± 1.3 | 23.3 ± 1.1 | 18.6 ± 2.7 | 42.6 ± 2.6 | <.0001 |
| Heart Age | y | 43.2 ± 0.936 | 42.3 ± 0.93 | 34.5 ± 2.19 | 51.9 ± 2.2 | <.0001 |
| RHI |  | 0.668 ± 0.025 | 0.673 ± 0.027 | 0.672 ± 0.0614 | 0.742 ± 0.061 | <.0001 |
| TYG-BMI |  | 229 ± 5.36 | 227 ± 5.4 | 213 ± 13.5 | 252 ± 13 | 0.0007 |

<sup>a</sup> - Values are expressed as mean ± SD.
<sup>b</sup> - p-values were derived using unpaired two-tailed t-tests of the LDLc 10<sup>th</sup> and 90<sup>th</sup> percentile of the population.
**Abbreviations:** ASCVD - atherosclerotic cardiovascular disease; BMI - body mass index; HDLc - high-density lipoprotein cholesterol; HOMA-IR - Homeostasis Model Assessment of Insulin Resistance; IR - insulin resistant; LDLc - low-density lipoprotein cholesterol; McAuley-ISI - McAuley-insulin sensitivity index; NEFA - non-esterified fatty acids; PPTG - postprandial triglyceride; QUICKI - Quantitative Insulin Sensitivity Check Index; Rem-c - remnant cholesterol; RHI - reactive hyperemia index; TG - triglycerides; Total-c - total cholesterol; TYG-BMI - triglyceride-glucose-BMI score.

### 2.2 Chemicals and reagents

A suite of 33 deuterated lipids were purchased from Avanti Polar lipids, Matreya LLC, and Larodan Fine Lipids (**Supplemental Table ST2**) ^7^. All solvents were Optima grade from Fischer Scientific unless otherwise stated.

### 2.3 Metabolomic analysis

Plasma sample (50 µL) metabolomic/lipidomic profiling was performed by ultra performance liquid chromatography - tandem mass spectrometer (UPLC-MS/MS) with a combination of chromatographic and flow infusion analyses on a Nexera X2 (Shimadzu, Columbia, MD)/API 6500 QTRAP (SCIEX, Framingham, MA) using MxP^®^ Quant 500 kits (biocrates life sciences gmbh, Innsbruck, Austria) as per manufacturer’s instructions. Fasting plasma was analyzed for 361 participants randomized across five plates. The analysis of a single NIST Standard Reference Material 1950: Metabolites in Human Plasma (Sigma-Aldrich, St Louis, MO) was processed on each plate along with method blanks, manufacturer provided quality control plasma dilutions and 7pt calibration curves for a subset of metabolites. Results were normalized across plates by the median values for the manufacturers mid-range quality control plasma sample.

### 2.4 Chromatographic lipidomic profiling

Lipids were isolated and quantified using minor modifications of published procedures. ^7^ Briefly, 10 µL of fasting plasma were combined with 100 µL methanol: n-butanol (1:1, v/v) containing 10 mM ammonium formate and the internal standards surrogates listed in Supplemental Table ST2. Samples were vortexed, held at 4 °C and centrifuged at 14,000 g for 10 min. The supernatant was transferred to a 0.22 µm 96-well polyvinylidene fluoride filter plate (Agilent, USA) and centrifuged at 1,000 g for 5 min with filtrates collected in 300 µL 96-well polypropylene storage microplates (ThermoFisher, USA). Filtrates were held at 4 °C until analysis (≤24 h). Lipids were then separated and measured by UPLC-MS/MS after optimizing chromatographic resolution and detection on the employed system. Specifically, data were collected on a flow through needle Acquity I-Class UPLC (Waters, Milford, MA) equipped with a 2.1 × 100 mm 1.8 µm Zorbax Eclipse Plus C18 column (Agilent, USA) held at 60 °C. A 1 µL extract aliquot was injected at an initial flow rate of 0.40 mL min ¹ with 10 % mobile phase B. Lipids were eluted with the following solvent gradient: 10 to 45 % B (0.0–2.7 min), 45 to 53 % B (2.7–2.8 min), 53 to 65 % B (2.8–9.0 min), 65 to 85 % B (9.0–10.0 min), 85 to 92 % B (10.0– 14.0 min), and 92 to 100 % B (14.0–14.1 min), with a 100 % B hold to 17.0 min. Re-equilibration to 10 % B was performed from 17.0–17.1 min, followed by a 2.9-min hold, giving a total run time of 20 min. Eluting residues were detected with an API 4500 QQQ (SCIEX) operating in positive-ion mode with scheduled multiple-reaction monitoring (MRM; **Supplemental Table ST3**). The UPLC gradient between 9 and 11 minutes was extended relative to Hyunh et al. to minimize concurrent MRMs in this region to < 50. As shown in **Supplemental Figure 1**, retention times for lipids eluting before 9 minutes remained aligned with the reference method, while the modified gradient extended elution between 9–14 minutes. These adjustments substantially improved chromatographic resolution in this region, without altering the relative retention times, enhancing detection of triglyceride (TGs) and cholesteryl esters (CEs), which together comprised nearly one-third of the quantified analytes. (https://metabolomics.baker.edu.au/method/lipids)

Mass spectrometer source parameters: ion-source temperature, 200 °C; curtain gas, 17 L min ¹; nebulizer gas, 20 psi; ion-spray voltage, +5.5 kV; and auxiliary gas, 10 L min ¹. All MRM transitions, declustering potentials, collision voltages, and lipid-class assignments are listed in Supplementary Table ST3.

Peak areas were auto integrated in MultiQuant v3.0.2 (SCIEX) with Gaussian smoothing of ≤ 0.5 with minor manual adjustments as needed upon inspection. Analytes are reported as area ratios to the closest-eluting internal standard. While there is overlap in the MXP^®^ 500 shotgun lipidomics and the chromatographic lipidomic platform described here, the later method identifies more unique lipid species, including deoxyceramides.

### 2.5 Statistics

Statistical analyses were performed in JMP Pro v17.2 (SAS Institute, Cary, NC, USA). Group mean differences were calculated by 2-tailed Student’s t-tests after normality assessment using the Anderson-Darling test. Metabolites with < 66% completeness across samples were excluded to avoid unstable estimates. ^19^ In the metabolomic dataset (n = 361), 244 participants had 6% [1 – 33%] values missing for 21 of 401 analytes. In the paired LDLc-dependent subgroups (n = 38), 34 participants had 8% [3 – 34%] values missing for 107 of 689 analytes, <1.5% of the dataset. Missing data were imputed using singular value decomposition imputation.^20^ Raw peak areas were log-transformed, then mean-centered and unit-variance scaled prior to downstream analysis ^21^. Johnson normalization was applied when needed.

Partial least squares regression (PLSR) was used to assess lipidomic associations with continuous LDLc concentrations using the MXP^®^ 500 metabolomic data for the full cohort (n = 361). Partial least squares discriminant analysis (PLS-DA) of the sex- and age-matched pairs (n = 38) was used to identify the strongest LDLc-defined atherosclerotic risk groups discriminating lipids. Lipids with PLS-DA variable importance in projection (VIP) >1.2 were selected for bootstrap forest with variable clustering to identify a minimum LDLc-stratified risk group discriminating set with leave-one-out cross validation. Model construction and validation used a five-fold cross-validation strategy detailed in **Supplemental Figure 2**. Correlation analysis was applied on the alanine:serine ratio with various cardiometabolic risk factors.

## Results

### 3.1 Participant characteristics

Characteristics of the study population have been previously reported in detail.^15^ Baseline cardiometabolic health characteristics are summarized in **Table 1** for the male and female populations, the 10^th^ and 90^th^ percentile across LDLc level, and the sex and age pair-matched high-and low-LDLc groups (Supplemental Table ST1). The LDLc, total cholesterol, and both fasting and postprandial TG concentrations were significantly higher in the high LDLc group (p <0.01). The mean absolute LDLc difference between 10^th^ and 90^th^ percentile groups exceeded 80 mg/dL. The LDLc 90^th^ percentile were also ∼20 years older, more insulin resistance, less insulin sensitive, and had higher ASCVD risk and Heart Age scores, slightly impaired microvascular function as measured by reactive hyperemia index (RHI) using digital peripheral tonometry, and modestly higher Triglyceride Glucose-BMI indices (TYG-BMI).^22,23^ Therefore, this population provided a robust biochemical difference for downstream lipidomic comparisons in an unmedicated, clinically healthy population.

### 3.2 LDLc-associated metabolite patterns in the metabolomic data

The biocrates MxP Quant 500 provided 401 fasting metabolite concentrations for 361 individuals. The concentration of the metabolites for full cohort from metabolomic is in **Supplemental Table ST4**. PLSR (Q² = 0.84; X² = 0.52; Y² = 0.63) of LDLc concentration as a continuous variable and metabolites with VIP scores >1.2 were considered key contributors (**Figure 2**). Directionality of association was determined based on center-scaled regression coefficients. PLSR identified 32 metabolites positively correlated with LDLc, including ceramides (n = 9), cholesterol esters (n = 7) and sphingomyelins (n = 6;). Conversely, ether-linked phosphatidylcholines (i.e. plasmalogens and/or other alkyl ether lipids; n = 14), amino acids (n = 10) were negatively correlated LDLc. Considering amino acids, serine was strongly negatively associated with LDLc, whereas alanine showed weaker negative LDLc associations, along with an increased alanine:serine ratio respectively. These multivariate associations were used for pathway prioritization rather than as evidence of causal direction.

**Figure 2:**
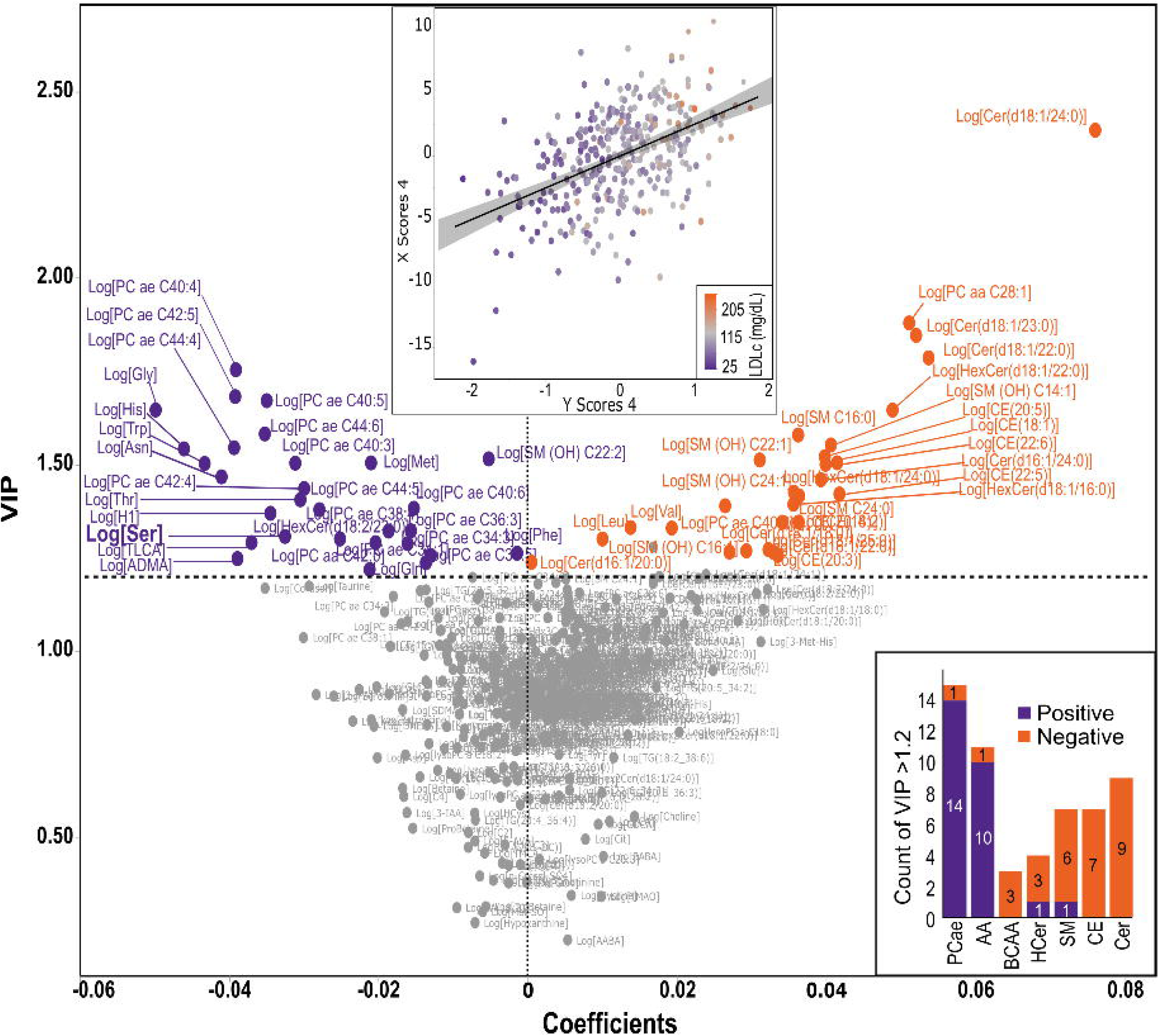
Partial-least squares-regression (PLSR) of metabolomic data against the population fasting LDLc concentrations. The PLSR-based coefficient x variable importance in projection (VIP) volcano plot is shown with variables with VIP > 1.2 labeled. ***Top middle inset -*** PLSR model (Q² = 0.84; X² = 0.52; Y² = 0.63) of LDLc concentrations using 401 metabolites from 361 participants. ***Lower right inset -*** the number of lipid species with VIP scores >1.2. Lipid classes with fewer than three contributing species were excluded. Color codes: purple, negatively correlated with LDLc; orange, positively correlated with LDLc.

### 3.3 LDLc-stratified group lipidomic differences

To expand the exploration of altered lipid species associated with LDLc, we next analyzed an LDLc-stratified, age -matched subset of males and females. A PLS-DA model (Q² = 0.41; X² = 0.34; Y² = 0.76) was built using the 639 unique lipid features. A total of 126 analytes (i.e. 18.3%) had a VIP > 1.2. The PLS-DA scores plot (**Figure 3**) shows modest separation between risk groups, with 90% confidence ellipses showing minimal overlap, consistent with the Q^2^Y of ≈ 0.4. The counts of lipids higher (n = 52) or lower (n = 49) in the high LDLc group are summarized by class in Figure 3. Glycerolipid (including TGs and diacylglycerides), sphingolipids (including sphingomyelins, deoxyceramides and ceramides), and cholesterol esters were elevated in the high LDLc group, while a variety of phospholipids were lower. Given the modest size of the expanded lipidomics subset, the PLS-DA model was interpreted as a lipid-feature discovery analysis, with emphasis on lipid-class patterns and pathway coherence rather than classification performance alone.

**Figure 3:**
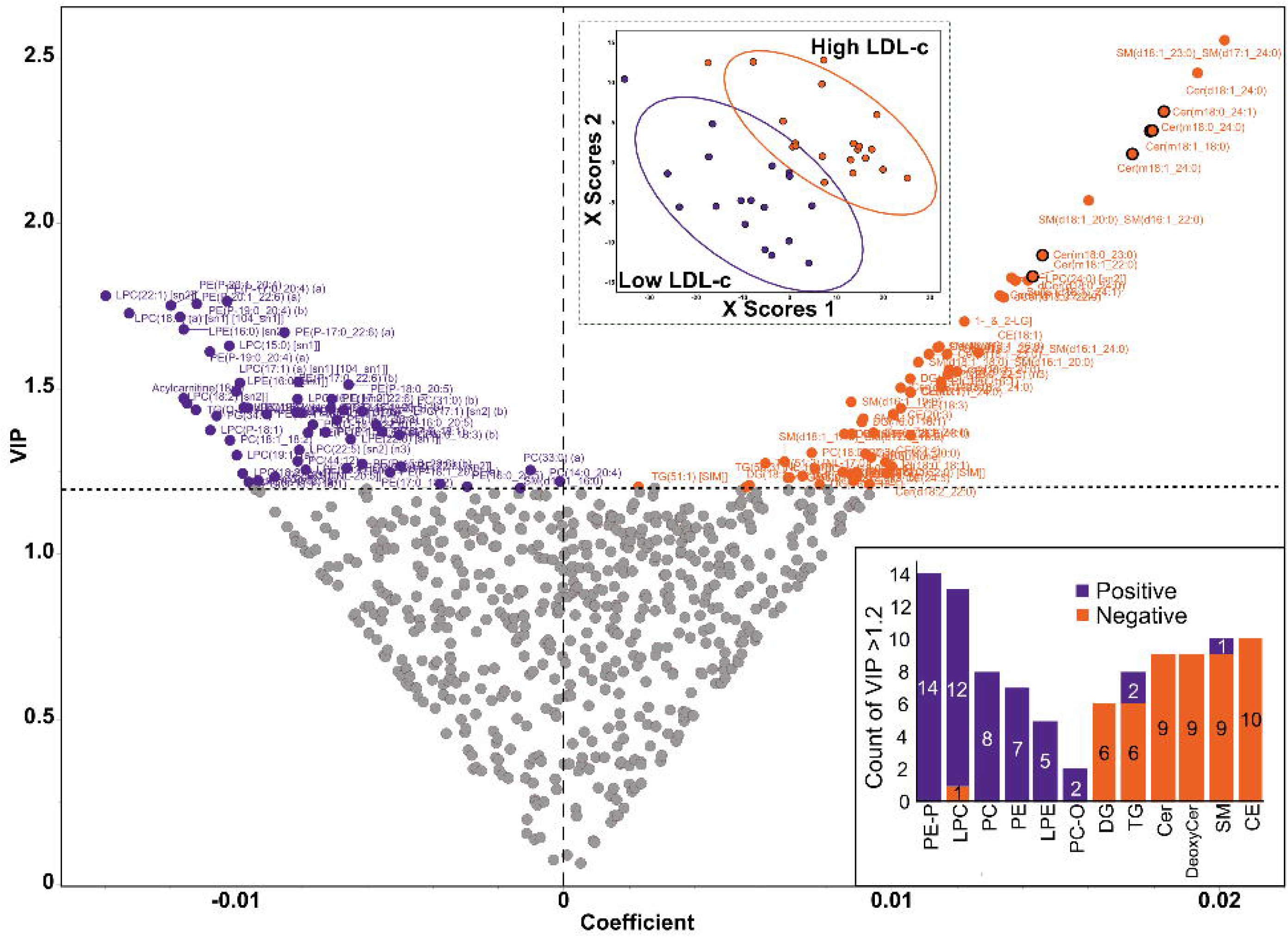
Lipidomic data partial least squares discriminate analysis (PLS-DA) of age- x sex-matched low- and high-risk LDLc subgroup pairs. The PLS-DA-based coefficient x variable importance in projection (VIP) volcano plot is shown with variables with VIP > 1.2 labeled. *Top middle inset* -PLSDA (Q² = 0.41; X² = 0.34; Y² = 0.76) Score plot with 90% confidence density ellipses. *Lower right inset -*the number of lipid species with VIP scores >1.2. Lipid classes with fewer than three contributing species were excluded. *Color codes*: purple, lower in High-risk; orange, higher in High-risk.

### 3.4 Serine-associated metabolic shifts link deoxysphingolipids, one-carbon metabolism-related metabolites and TG-associated traits

Canonically, serine palmitoyl-transferase condenses palmitoyl-CoA with serine to initiate ceramide biosynthesis. However, substitution with alanine when serine is insufficient yields deoxyceramides, which accumulate as lipotoxic “dead-end” metabolites.^24^ In the metabolomic dataset, alanine was positively correlated with LDL-C in pairwise analysis but had a small negative regression coefficient in the PLSR model. Association involving serine and deoxyceramides were further examined in the expanded lipidomic subset. Plasma serine concentrations were lower in the high-risk group than in low-risk group (p = 0.03). Lower serine concentrations were associated with higher total deoxyceramide abundance (p = 0.0012) and the deoxyceramide:ceramide ratio showed positive correlations with the alanine:serine ratio (p = 0.017). After adjustment for alanine concentrations, deoxyceramide is negative associated with the serine (p = 0.17).

Associations between the alanine ratio and other cardiometabolic traits were then examined across complete cohort, with no significant effect modification by sex. Among measured cardiometabolic traits, the alanine:serine ratio explained 13.7% of the variance in fasting TGs (r = 0.37; p < 0.0001), exceeding the 3.6% explanation of LDLc (r = 0.19; p = 0.004), supporting a link between serine-associated substrate balance and TG-rich lipoprotein biology (**Figure 4**).

**Figure 4:**
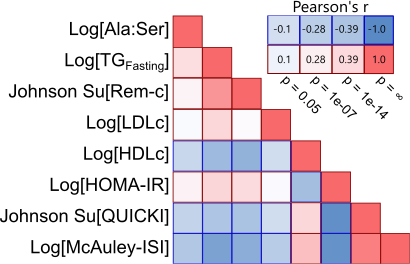
The plasma alanine:serine (Ala:Ser) ratio correlation with clinical cardiometabolic risk factors. The Ala:Ser was positively correlated with fasting triglycerides (TG_Fasting_; r = 0.37) > insulin resistance (HOMA-IR; r= 0.26) > remnant cholesterol (Rem-c; r = 0.25) > low density lipoprotein cholesterol (LDLc; r = 0.19). Conversely, this ratio was negatively correlated with the McAuley-insulin sensitivity index (McAuley-ISI; r = -0.35) > Quantitative Insulin Sensitivity Check Index (QUICKI; r = -0.22) > high density lipoprotein cholesterol (HDLc; r = -0.18). Transformation of metabolites were the most normal distribution.

## Discussion

While metabolic changes associated with cardiovascular disease risk are numerous, recognition of these changes in early stages of disease are poorly understood and may underlie factors associated with LDLc-independent (i.e. residual) cardiovascular disease risk. In the current study, we observed metabolic patterns consistent with coordinated alterations in ceramide, deoxysphingolipid, amino acid, and one-carbon-related metabolism in a clinically healthy cohort. The first step in sphingolipid metabolism is the condensation of serine and saturated fatty acids by serine palmitoyl transferase (SPT). Serine deprivation *in vitro* reduces cellular glycine levels, and results in mitochondrial dysfunction due to constrained sphingolipid metabolism.^25,26^ Elevated levels of ceramides, deoxyceramides, and sphingomyelins were a dominant phenotype associated with LDLc enrichment in our clinically healthy cohort. This is consistent with previously reported ceramide-based cardiovascular risk metrics.^27–29^ Ceramide accumulation has been widely associated with insulin resistance and lipid oversupply, particularly in the context of saturated fatty acid availability favoring *de novo* ceramide synthesis. ^30^ Beyond canonical ceramide metabolism, we also observed plasma concentrations of alanine and serine tightly associated with deoxyceramide and ceramide concentrations. Notably, negative correlations between deoxyceramides and serine concentrations were stronger than positive correlations with alanine concentrations in the LDLc-stratified subgroups, suggesting that deoxyceramide abundance was driven by reduced serine availability rather than alanine enrichment. Previous studies have shown that elevated alanine relative to serine can promote the formation of deoxy-sphingolipids through altered SPT substrate utilization ^31^, with the production of deoxy-sphingolipids implicated in metabolic disorders including diabetes ^32,33^, diabetic neuropathy ^34,35^, and metabolic dysfunction-associated steatotic liver disease ^36^, as well as increased epicardial fat volume.^37^ Notably a prospective study of diabetes development by Chen *et al.* identified baseline plasma 1-deoxyceramide concentrations as a prominent predictor of 10-yr diabetes risk ^33^. Therefore, the enrichment in deoxyceramides reflects reduced serine availability and a subsequent metabolic imbalance observable in the plasma alanine:serine ratio that may also serve as markers of metabolic dysregulation relevant to cardiometabolic risk.

Beyond its role as a key substrate in sphingolipid biosynthesis, serine is also a critical substrate driving one-carbon metabolism through its interconversion with glycine via serine hydroxymethyltransferase (i.e. SHMT) in the folate cycle.^14,38^ The differences between the participants in the 10th and 90th LDLc percentile in sphingolipid, transsulfuration and one-carbon metabolism have been mapped to highlight the integrated changes observed in these pathways (**Figure 5**). Along with lower serine levels, we observed lower glycine levels and somewhat lower methionine levels, changes previously associated with insulin resistance ^39–41^. While changes in homocysteine levels were not observed, the coordinated change in serine, glycine and methionine suggest constraints on one-carbon metabolism, theoretically stressing multiple metabolic systems. Serine also participates in the transsulfuration pathway condensing with homocysteine to form cysteine, leading to both taurine and glutathione production.^42^ As lower serine concentrations were observed, multiple distal transsulfuration products were found at higher levels, including the cysteine disulfide cystine, consistent with elevated oxidative stress.^43^ The lower ether-linked phosphatidylcholine (PC-ae) levels observed in individuals with higher LDLc further supports an elevated oxidative stress environment.^44,45^ Together, these findings are consistent with a model in which sphingolipid metabolism may place greater demand on the circulating serine pool, reducing serine availability for glycine and one-carbon-related metabolism. However, because pathway flux and enzymatic activity were not directly assessed, this interpretation should be considered hypothesis-generating. Prior intervention studies involving serine supplementation in neuropathy ^46^ and SPT inhibition in atherosclerosis ^47^ provide biological rationale for future testing, but do not establish these mechanisms in the present cohort.

**Figure 5:**
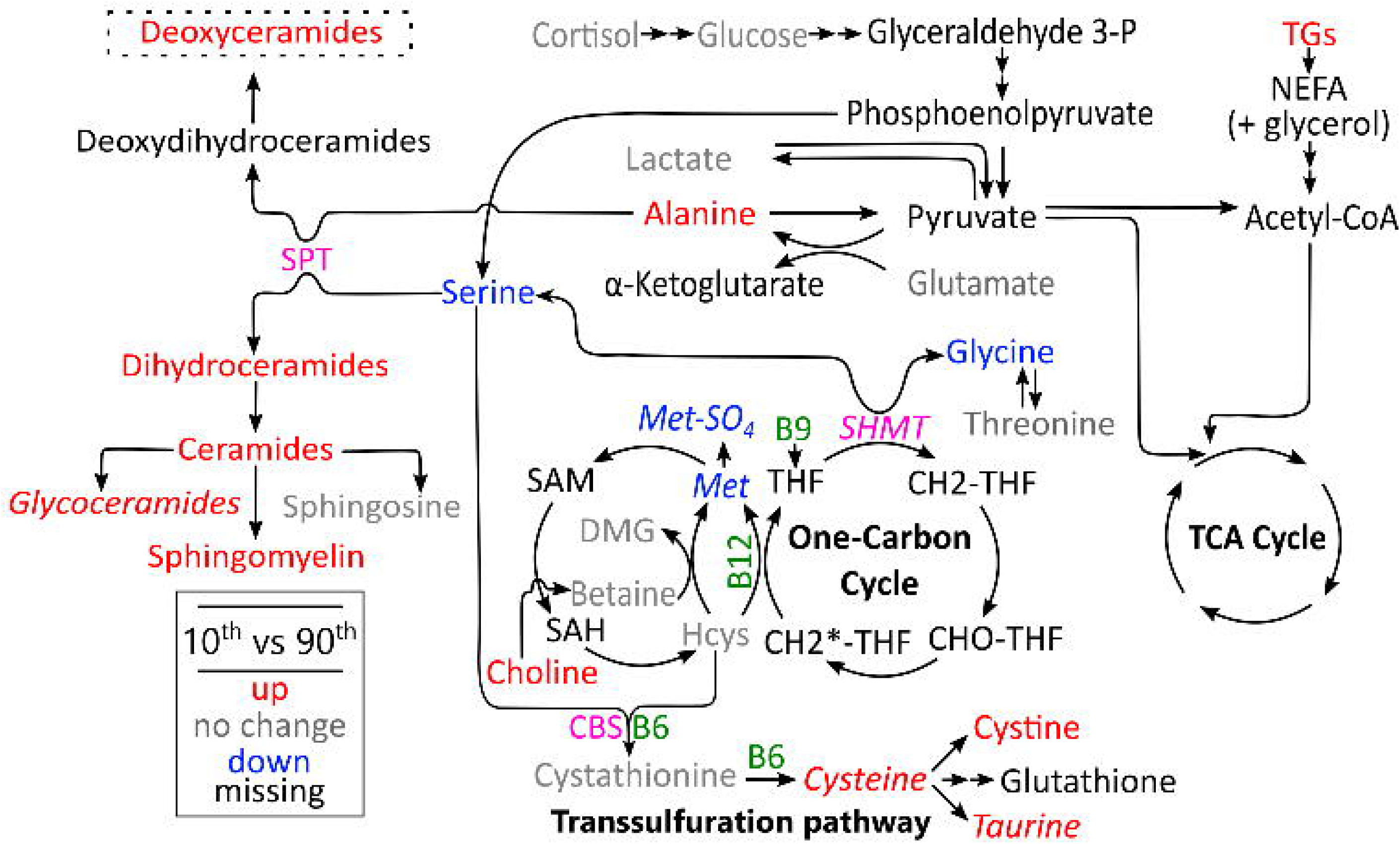
Changes in serine-associated metabolism link sphingolipid and one-carbon metabolism to cardiovascular risk factors. Mean differences in the 90^th^ vs 10^th^ percentile of the study population LDL cholesterol (LDLc) levels are used to annotate the pathways except for deoxyceramide changes established in the 19 pair matched subgroup with dash line. Changes in italicized metabolites are at p <0.1. ***Color code:*** red - higher with LDLc; blue – lower with LDLc; green - critical B vitamins; magenta – serine metabolizing enzymes. *Abbreviations:* CBS - Cystathionine β-synthase; DMG – dimethylglycine; Hcys – homocysteine; Met – methionine; NEFA – non-esterified fatty acids; SAM – S-adenosylmethionine; SAH – S-adenosylhomocysteine; SHMT - serine hydroxymethyltransferase; SPT - serine palmitoyl-transferase; TG – triglyceride; THF – tetrahydrofolate.

Because LDLc enrichment was accompanied by differences in age, insulin-resistance indices, and TG-related traits, subsequent pathway-level interpretations were evaluated in the context of these cardiometabolic covariates. Although therapeutic LDLc lowering reduces atherosclerosis risk, elevated fasting TGs and insulin resistance are independent critical factors associated with uncontrolled adverse cardiac events.^48–50^ With respect to cardiometabolic traits associated with residual cardiovascular risk, the alanine:serine ratio showed a stronger association with fasting TGs than with LDLc. Therefore, the alanine:serine ratio may capture TG-associated metabolic shifts not fully reflected by LDLc, and the HOMA-IR. Whether this ratio improves prediction of cardiovascular outcomes will require validation in longitudinal cohorts. More broadly, CVD risk is influenced not only by circulating LDLc concentrations, but also by the compositional features of lipoproteins, particularly sphingolipid subclasses such as ceramides and deoxyceramides.^51,52^

In parallel, fasting TG levels, reflecting the burden of TG-rich lipoproteins and their remnants, have been consistently associated with increased CVD risk, including in populations with otherwise controlled LDLc levels.^48,53^ It is consistent with prior evidence linking hypertriglyceridemia to altered sphingolipid metabolism, particularly ceramide accumulation, although fewer studies have addressed whether deoxysphingolipid metabolism is similarly altered. During the fasting stage, alanine will be prioritized to support the gluconeogenesis via pyruvate, and alanine source is elevated by breakdown of the muscle protein.^54^ However, when this liver-muscle metabolic regulation is impaired by insulin resistance, it will lead to the accumulation of the precursor like alanine in the circulation.^55,56^ At the meantime, liver will be favor the de novo lipogenesis which could explain the observation of the increase of TG. ^35^ Also, the observation of the inverse association between serine and TG level might align with the existing research that the lower serine level and high TG level as a sign of the metabolic dysregulation.^35,57^

Our findings extend this framework by suggesting that the metabolic demands associated with exaggerated sphingolipid metabolism have cascading effects resulting in amino acid metabolic remodeling, reflected in the alanine:serine balance. These findings support the hypothesis that the alanine:serine ratio may reflect substrate-handling capacity in early cardiometabolic risk, motivating future studies testing whether modulation of serine availability, one-carbon cofactors, or SPT activity can alter sphingolipid remodeling and cardiometabolic risk and its markers.

Therefore, the alanine:serine ratio may provide an integrated metabolic readout connecting LDLc-associated sphingolipid remodeling with triglyceride-associated cardiometabolic traits. Due to the broad and balanced cross-sectional nature of study across sexes, ages, body weights, and ethnic backgrounds, these findings may be broadly generalizable to other populations. Future studies should determine whether this ratio adds predictive value for cardiovascular risk beyond conventional lipid measures.

### Limitations and Future Directions

Plasma metabolomic-lipidomic measurements do not necessarily reflect metabolic processes within specific cells, limiting the ability to extend the reported observations without further study. While drawn from a modestly sized population, the subset investigating broader lipidomic coverage was constrained by a small sample size. Nonetheless, the reproducible identification of elevated sphingolipid metabolism across platforms in a clinically healthy population, and the identification of deoxyceramide changes in association with the alanine:serine ratio, supports the idea that these metabolic alterations may be detectable before overt clinical disease. Future studies should validate these associations in longitudinal cohorts, integrating amino acid and nutrient pathways to capture metabolic interactions and determine whether incorporating sphingolipids into risk models improves prediction of cardiovascular outcomes when LDLc is controlled. As this study was designed to capture population-level metabolic associations, enzymatic activity and pathway flux were not directly assessed. The association of the pathway observed in The Jackson Heart Study with diabetes occurrence at a 10-year follow-up should be repeated with a focus on cardiovascular outcomes.^33^

## Conclusion

Targeted metabolomic and lipidomic analyses identify serine-associated metabolism connecting shifts in LDLc-associated sphingolipid remodeling, one-carbon metabolism-related metabolites, and TG-associated cardiometabolic traits. Longitudinal and mechanistic studies are needed to determine whether interventions targeting serine availability, one-carbon cofactors, or SPT activity influence sphingolipid remodeling and cardiovascular risk markers. These findings support the hypothesis that reduced serine availability may be associated with sphingolipid and one-carbon metabolism changes linked to cardiometabolic risk.

## Acknowledgements

The authors would like to specifically acknowledge and thank Peter J. Meikle, Kevin Hyunh and Natalie Mellett of the Baker Heart and Diabetes Institute, Melbourne, VIC, 3004, Australia for graciously sharing detailed analytical protocols, reagent recipes, technical advice and their time, as we adapted and implemented their outstanding targeted lipidomics method. Drs. Miekle and Hyunh also provided exemplary pre-submission external reviews that substantially improved the final manuscript. Virginia Artegoitia for initial efforts on lipidomic method transfer from the Baker Institute to the WHNRC.

## Author Contribution

The authors’ responsibilities were as follows **-** MH, BJB, JN: conceived and developed the research plan; NLK, BJB JWN: conducted the clinical research; MH, JWN: generated data; MH, JWN: lipidomic method transfer, modification and implementation; MH, JN, ST, DKMT: developed the statistical analysis plan; MH, JWN: performed the statistical analysis; MH, JWN: wrote the primary manuscript; DKMT, ST, BJB: provided significant editorial input of manuscript; all authors: have primary responsibility for final content and read and approved the final manuscript.

## Data Sharing

Data described in the manuscript are available upon reasonable request. All statistical code will be made available upon request.

## Abbreviations

BMI: body mass index
CE: cholesteryl ester
Cer: ceramide
CERT: ceramide risk score
CVD: cardiovascular disease
DAG: diacylglycerol
dhCer: dihydroceramide
dhHexCer: dihydrohexosylceramide
Hcys: homocysteine
HDL-c: high-density lipoprotein cholesterol
HexCer: hexosylceramide
HOMA-IR: Homeostasis Model Assessment of Insulin Resistance
UPLC–MS/MS: ultra performance liquid chromatography–tandem mass spectrometry
LDLc: low-density lipoprotein cholesterol
LPC: lysophosphatidylcholine
LPE: lysophosphatidylethanolamine
McAuley-ISI: McAuley-insulin sensitivity index
NEFA: non-esterified fatty acids
PC: phosphatidylcholine
PE: phosphatidylethanolamine
PI: phosphatidylinositol
PLS-DA: PLS-discriminant analysis
PLSR: PLS-regression
QUICKI: Quantitative Insulin Sensitivity Check Index
Rem-c: remnant cholesterol
RHI: reactive hyperemia index
SAH: S-adenosylhomocysteine
SAM: S-adenosylmethionine
SHMT: serine hydroxymethyltransferase
SM: sphingomyelin
SPT: serine palmitoyl transferase
TGs: triglycerides
TYG-BMI: triglyceride/glucose/BMI.

## Reference

1. Virani SS, Alonso A, Benjamin EJ, Bittencourt MS, Callaway CW, Carson AP, Chamberlain AM, Chang AR, Cheng S, Delling FN, et al. Heart Disease and Stroke Statistics-2020 Update: A Report From the American Heart Association. Circulation. 2020;141:e139–e596. doi: 10.1161/CIR.0000000000000757

2. Boren J, Chapman MJ, Krauss RM, Packard CJ, Bentzon JF, Binder CJ, Daemen MJ, Demer LL, Hegele RA, Nicholls SJ, et al. Low-density lipoproteins cause atherosclerotic cardiovascular disease: pathophysiological, genetic, and therapeutic insights: a consensus statement from the European Atherosclerosis Society Consensus Panel. European heart journal. 2020;41:2313–2330. doi: 10.1093/eurheartj/ehz962

3. Keys A. The diet and the development of coronary heart disease. J Chronic Dis. 1956;4:364–380. doi: 10.1016/0021-9681(56)90040-6

4. Goldstein JL, Brown MS. A century of cholesterol and coronaries: from plaques to genes to statins. Cell. 2015;161:161–172. doi: 10.1016/j.cell.2015.01.036

5. Poss AM, Maschek JA, Cox JE, Hauner BJ, Hopkins PN, Hunt SC, Holland WL, Summers SA, Playdon MC. Machine learning reveals serum sphingolipids as cholesterol-independent biomarkers of coronary artery disease. J Clin Invest. 2020;130:1363–1376. doi: 10.1172/JCI131838

6. Alshehry ZH, Mundra PA, Barlow CK, Mellett NA, Wong G, McConville MJ, Simes J, Tonkin AM, Sullivan DR, Barnes EH, et al. Plasma Lipidomic Profiles Improve on Traditional Risk Factors for the Prediction of Cardiovascular Events in Type 2 Diabetes Mellitus. Circulation. 2016;134:1637–1650. doi: 10.1161/CIRCULATIONAHA.116.023233

7. Huynh K, Barlow CK, Jayawardana KS, Weir JM, Mellett NA, Cinel M, Magliano DJ, Shaw JE, Drew BG, Meikle PJ. High-Throughput Plasma Lipidomics: Detailed Mapping of the Associations with Cardiometabolic Risk Factors. Cell Chem Biol. 2019;26:71–84 e74. doi: 10.1016/j.chembiol.2018.10.008

8. Ruuth M, Nguyen SD, Vihervaara T, Hilvo M, Laajala TD, Kondadi PK, Gistera A, Lahteenmaki H, Kittila T, Huusko J, et al. Susceptibility of low-density lipoprotein particles to aggregate depends on particle lipidome, is modifiable, and associates with future cardiovascular deaths. European heart journal. 2018;39:2562–2573. doi: 10.1093/eurheartj/ehy319

9. Mantovani A, Lunardi G, Bonapace S, Molinero AE, Morandin R, Fiorio V, Molon G, Byrne CD, Targher G. Ceramide-based risk score CERT-1 improves risk prediction for overall mortality and adverse cardiovascular outcomes in patients with and without cardiovascular disease: A prospective cohort study. Diabetes Obes Metab. 2025;27:1488–1497. doi: 10.1111/dom.16156

10. Papazoglou AS, Stalikas N, Moysidis DV, Otountzidis N, Kartas A, Karagiannidis E, Giannakoulas G, Sianos G. CERT2 ceramide- and phospholipid-based risk score and major adverse cardiovascular events: A systematic review and meta-analysis. J Clin Lipidol. 2022;16:272–276. doi: 10.1016/j.jacl.2022.02.001

11. Leiherer A, Muendlein A, Saely CH, Geiger K, Brandtner EM, Heinzle C, Gaenger S, Mink S, Laaksonen R, Fraunberger P, et al. Coronary Event Risk Test (CERT) as a Risk Predictor for the 10-Year Clinical Outcome of Patients with Peripheral Artery Disease. J Clin Med. 2023;12. doi: 10.3390/jcm12196151

12. Ducker GS, Rabinowitz JD. One-Carbon Metabolism in Health and Disease. Cell Metab. 2017;25:27–42. doi: 10.1016/j.cmet.2016.08.009

13. Lind MV, Lauritzen L, Vestergaard H, Hansen T, Pedersen O, Kristensen M, Ross AB. One-carbon metabolism markers are associated with cardiometabolic risk factors. Nutr Metab Cardiovasc Dis. 2018;28:402–410. doi: 10.1016/j.numecd.2018.01.005

14. Al Qassab M, Bou Ghanem A, Hussayni Y, Kadbey R, Ratel Y, Yehya S, Zahr M, Ghadieh HE, Abi Khattar Z, Azar S, et al. One-carbon metabolism and cardiovascular disease: Molecular mechanisms, genetic influences, and epigenetic regulation. Biochem Biophys Rep. 2026;46:102544. doi: 10.1016/j.bbrep.2026.102544

15. Baldiviez LM, Keim NL, Laugero KD, Hwang DH, Huang L, Woodhouse LR, Burnett DJ, Zerofsky MS, Bonnel EL, Allen LH, et al. Design and implementation of a cross-sectional nutritional phenotyping study in healthy US adults. BMC Nutrition. 2017;3. doi: 10.1186/s40795-017-0197-4

16. Chen H, Wang Z, Qin M, Zhang B, Lin L, Ma Q, Liu C, Chen X, Li H, Lai W, et al. Comprehensive Metabolomics Identified the Prominent Role of Glycerophospholipid Metabolism in Coronary Artery Disease Progression. Front Mol Biosci. 2021;8:632950. doi: 10.3389/fmolb.2021.632950

17. Meikle PJ, Wong G, Tsorotes D, Barlow CK, Weir JM, Christopher MJ, MacIntosh GL, Goudey B, Stern L, Kowalczyk A, et al. Plasma lipidomic analysis of stable and unstable coronary artery disease. Arterioscler Thromb Vasc Biol. 2011;31:2723–2732. doi: 10.1161/ATVBAHA.111.234096

18. Expert Panel on Detection E, Treatment of High Blood Cholesterol in A. Executive Summary of The Third Report of The National Cholesterol Education Program (NCEP) Expert Panel on Detection, Evaluation, And Treatment of High Blood Cholesterol In Adults (Adult Treatment Panel III). JAMA. 2001;285:2486-2497. doi: 10.1001/jama.285.19.2486

19. Taylor S, Ponzini M, Wilson M, Kim K. Comparison of imputation and imputation-free methods for statistical analysis of mass spectrometry data with missing data. Brief Bioinform. 2022;23. doi: 10.1093/bib/bbab353

20. Dakic A, Wu J, Wang T, Huynh K, Mellett N, Duong T, Beyene HB, Magliano DJ, Shaw JE, Carrington MJ, et al. Imputation of plasma lipid species to facilitate integration of lipidomic datasets. Nat Commun. 2024;15:1540. doi: 10.1038/s41467-024-45838-3

21. Van den Berg RA, Hoefsloot HC, Westerhuis JA, Smilde AK, van der Werf MJ. Centering, scaling, and transformations: improving the biological information content of metabolomics data. BMC Genomics. 2006;7:142. doi: 10.1186/1471-2164-7-142

22. Axtell AL, Gomari FA, Cooke JP. Assessing endothelial vasodilator function with the Endo-PAT 2000. J Vis Exp. 2010. doi: 10.3791/2167

23. Zhou S, Guo R. Understanding triglyceride glucose-body mass index: implications for diabetes and cardiovascular disease management. Front Endocrinol (Lausanne). 2025;16:1675270. doi: 10.3389/fendo.2025.1675270

24. Lone MA, Santos T, Alecu I, Silva LC, Hornemann T. 1-Deoxysphingolipids. Biochim Biophys Acta Mol Cell Biol Lipids. 2019;1864:512–521. doi: 10.1016/j.bbalip.2018.12.013

25. Gao X, Lee K, Reid MA, Sanderson SM, Qiu C, Li S, Liu J, Locasale JW. Serine Availability Influences Mitochondrial Dynamics and Function through Lipid Metabolism. Cell Rep. 2018;22:3507–3520. doi: 10.1016/j.celrep.2018.03.017

26. Alecu I, Tedeschi A, Behler N, Wunderling K, Lamberz C, Lauterbach MA, Gaebler A, Ernst D, Van Veldhoven PP, Al-Amoudi A, et al. Localization of 1-deoxysphingolipids to mitochondria induces mitochondrial dysfunction. J Lipid Res. 2017;58:42–59. doi: 10.1194/jlr.M068676

27. Di Pietro P, Izzo C, Abate AC, Iesu P, Rusciano MR, Venturini E, Visco V, Sommella E, Ciccarelli M, Carrizzo A, et al. The Dark Side of Sphingolipids: Searching for Potential Cardiovascular Biomarkers. Biomolecules. 2023;13. doi: 10.3390/biom13010168

28. de Carvalho LP, Tan SH, Ow GS, Tang Z, Ching J, Kovalik JP, Poh SC, Chin CT, Richards AM, Martinez EC, et al. Plasma Ceramides as Prognostic Biomarkers and Their Arterial and Myocardial Tissue Correlates in Acute Myocardial Infarction. JACC Basic Transl Sci. 2018;3:163–175. doi: 10.1016/j.jacbts.2017.12.005

29. Augusto SN, Jr., Suresh A, Tang WHW. Ceramides as Biomarkers of Cardiovascular Diseases and Heart Failure. Curr Heart Fail Rep. 2024;22:2. doi: 10.1007/s11897-024-00689-3

30. Sokolowska E, Blachnio-Zabielska A. The Role of Ceramides in Insulin Resistance. Frontiers in Endocrinology. 2019;10. doi: 10.3389/fendo.2019.00577

31. Esaki K, Sayano T, Sonoda C, Akagi T, Suzuki T, Ogawa T, Okamoto M, Yoshikawa T, Hirabayashi Y, Furuya S. L-Serine Deficiency Elicits Intracellular Accumulation of Cytotoxic Deoxysphingolipids and Lipid Body Formation. J Biol Chem. 2015;290:14595–14609. doi: 10.1074/jbc.M114.603860

32. Mwinyi J, Bostrom A, Fehrer I, Othman A, Waeber G, Marti-Soler H, Vollenweider P, Marques-Vidal P, Schioth HB, von Eckardstein A, et al. Plasma 1-deoxysphingolipids are early predictors of incident type 2 diabetes mellitus. PLoS One. 2017;12:e0175776. doi: 10.1371/journal.pone.0175776

33. Chen ZZ, Pacheco JA, Gao Y, Deng S, Peterson B, Shi X, Zheng S, Tahir UA, Katz DH, Cruz DE, et al. Nontargeted and Targeted Metabolomic Profiling Reveals Novel Metabolite Biomarkers of Incident Diabetes in African Americans. Diabetes. 2022;71:2426–2437. doi: 10.2337/db22-0033

34. Hornemann T. Serine deficiency causes complications in diabetes. Nature. 2023;614:42–43. doi: 10.1038/d41586-023-00054-9

35. Handzlik MK, Gengatharan JM, Frizzi KE, McGregor GH, Martino C, Rahman G, Gonzalez A, Moreno AM, Green CR, Guernsey LS, et al. Insulin-regulated serine and lipid metabolism drive peripheral neuropathy. Nature. 2023;614:118–124. doi: 10.1038/s41586-022-05637-6

36. Wipfli F, M AL, von Eckardstein A, Verrijken A, Francque S, Weyler J, Van Gaal L, Staels B, Hornemann T. Increased 1-deoxysphingolipids caused by an altered plasma alanine to serine ratio are associated with metabolic dysfunction-associated steatotic liver disease (MASLD). Metabolomics. 2025;21:157. doi: 10.1007/s11306-025-02359-4

37. Leandro AC, Michael LF, Almeida M, Kuokkanen M, Huynh K, Giles C, Duong T, Diego VP, Duggirala R, Clarke GD, et al. Influence of the Human Lipidome on Epicardial Fat Volume in Mexican American Individuals. Front Cardiovasc Med. 2022;9:889985. doi: 10.3389/fcvm.2022.889985

38. Pan S, Fan M, Liu Z, Li X, Wang H. Serine, glycine and one carbon metabolism in cancer (Review). Int J Oncol. 2021;58:158–170. doi: 10.3892/ijo.2020.5158

39. Wittemans LBL, Lotta LA, Oliver-Williams C, Stewart ID, Surendran P, Karthikeyan S, Day FR, Koulman A, Imamura F, Zeng L, et al. Assessing the causal association of glycine with risk of cardio-metabolic diseases. Nat Commun. 2019;10:1060. doi: 10.1038/s41467-019-08936-1

40. Handzlik MK, Metallo CM. Sources and Sinks of Serine in Nutrition, Health, and Disease. Annu Rev Nutr. 2023;43:123–151. doi: 10.1146/annurev-nutr-061021-022648

41. Rom O, Liu Y, Finney AC, Ghrayeb A, Zhao Y, Shukha Y, Wang L, Rajanayake KK, Das S, Rashdan NA, et al. Induction of glutathione biosynthesis by glycine-based treatment mitigates atherosclerosis. Redox Biol. 2022;52:102313. doi: 10.1016/j.redox.2022.102313

42. Majtan T, Mijatovic E, Petrosino M. Understanding the Impact of Mutations in the Cystathionine Beta-Synthase Gene: Towards Novel Therapeutics for Homocystinuria. Mol Cell Biol. 2025;45:327–342. doi: 10.1080/10985549.2025.2511338

43. Droge W. Oxidative stress and ageing: is ageing a cysteine deficiency syndrome? Philos Trans R Soc Lond B Biol Sci. 2005;360:2355–2372. doi: 10.1098/rstb.2005.1770

44. Graessler J, Schwudke D, Schwarz PE, Herzog R, Shevchenko A, Bornstein SR. Top-down lipidomics reveals ether lipid deficiency in blood plasma of hypertensive patients. PLoS One. 2009;4:e6261. doi: 10.1371/journal.pone.0006261

45. Gonzalez-Covarrubias V, Beekman M, Uh HW, Dane A, Troost J, Paliukhovich I, van der Kloet FM, Houwing-Duistermaat J, Vreeken RJ, Hankemeier T, et al. Lipidomics of familial longevity. Aging Cell. 2013;12:426–434. doi: 10.1111/acel.12064

46. Garofalo K, Penno A, Schmidt BP, Lee HJ, Frosch MP, von Eckardstein A, Brown RH, Hornemann T, Eichler FS. Oral L-serine supplementation reduces production of neurotoxic deoxysphingolipids in mice and humans with hereditary sensory autonomic neuropathy type 1. J Clin Invest. 2011;121:4735–4745. doi: 10.1172/JCI57549

47. Park TS, Rosebury W, Kindt EK, Kowala MC, Panek RL. Serine palmitoyltransferase inhibitor myriocin induces the regression of atherosclerotic plaques in hyperlipidemic ApoE-deficient mice. Pharmacol Res. 2008;58:45–51. doi: 10.1016/j.phrs.2008.06.005

48. Schuitema PCE, Visseren FLJ, Nordestgaard BG, Teraa M, van der Meer MG, Ruigrok YM, Onland-Moret NC, Koopal C, Group U-SS. Elevated triglycerides are related to higher residual cardiovascular disease and mortality risk independent of lipid targets and intensity of lipid-lowering therapy in patients with established cardiovascular disease. Atherosclerosis. 2025;408:120411. doi: 10.1016/j.atherosclerosis.2025.120411

49. Gomez-Delgado F, Raya-Cruz M, Katsiki N, Delgado-Lista J, Perez-Martinez P. Residual cardiovascular risk: When should we treat it? Eur J Intern Med. 2024;120:17–24. doi: 10.1016/j.ejim.2023.10.013

50. Vargas-Vazquez A, Fermin-Martinez CA, Antonio-Villa NE, Fernandez-Chirino L, Ramirez-Garcia D, Davila-Lopez G, Diaz-Sanchez JP, Aguilar-Salinas CA, Seiglie JA, Bello-Chavolla OY. Insulin resistance potentiates the effect of remnant cholesterol on cardiovascular mortality in individuals without diabetes. Atherosclerosis. 2024;395:117508. doi: 10.1016/j.atherosclerosis.2024.117508

51. Chapman MJ, Orsoni A, Tan R, Mellett NA, Nguyen A, Robillard P, Giral P, Therond P, Meikle PJ. LDL subclass lipidomics in atherogenic dyslipidemia: effect of statin therapy on bioactive lipids and dense LDL. J Lipid Res. 2020;61:911–932. doi: 10.1194/jlr.P119000543

52. Yi C, Huynh K, Schooneveldt Y, Olshansky G, Liang A, Wang T, Beyene HB, Dakic A, Wu J, Cinel M, et al. Statin effects on the lipidome: Predicting statin usage and implications for cardiovascular risk prediction. J Lipid Res. 2025;66:100800. doi: 10.1016/j.jlr.2025.100800

53. Raposeiras-Roubin S, Rossello X, Oliva B, Fernandez-Friera L, Mendiguren JM, Andres V, Bueno H, Sanz J, Martinez de Vega V, Abu-Assi E, et al. Triglycerides and Residual Atherosclerotic Risk. J Am Coll Cardiol. 2021;77:3031-3041. doi: 10.1016/j.jacc.2021.04.059

54. Sookoian S, Pirola CJ. Alanine and aspartate aminotransferase and glutamine-cycling pathway: their roles in pathogenesis of metabolic syndrome. World J Gastroenterol. 2012;18:3775–3781. doi: 10.3748/wjg.v18.i29.3775

55. Chen X, Iqbal N, Boden G. The effects of free fatty acids on gluconeogenesis and glycogenolysis in normal subjects. J Clin Invest. 1999;103:365–372. doi: 10.1172/JCI5479

56. Alves-Bezerra M, Cohen DE. Triglyceride Metabolism in the Liver. Compr Physiol. 2017;8:1–8. doi: 10.1002/cphy.c170012

57. Truong V, Huang S, Dennis J, Lemire M, Zwingerman N, Aissi D, Kassam I, Perret C, Wells P, Morange PE, et al. Blood triglyceride levels are associated with DNA methylation at the serine metabolism gene PHGDH. Scientific reports. 2017;7:11207. doi: 10.1038/s41598-017-09552-z

