## Supplemental Materials (FigS1-2, Tables S1-4) for "Serine metabolism changes link sphingolipid and one-carbon metabolism to cardiovascular risk factors in a cross-sectional study of asymptomatic adults"

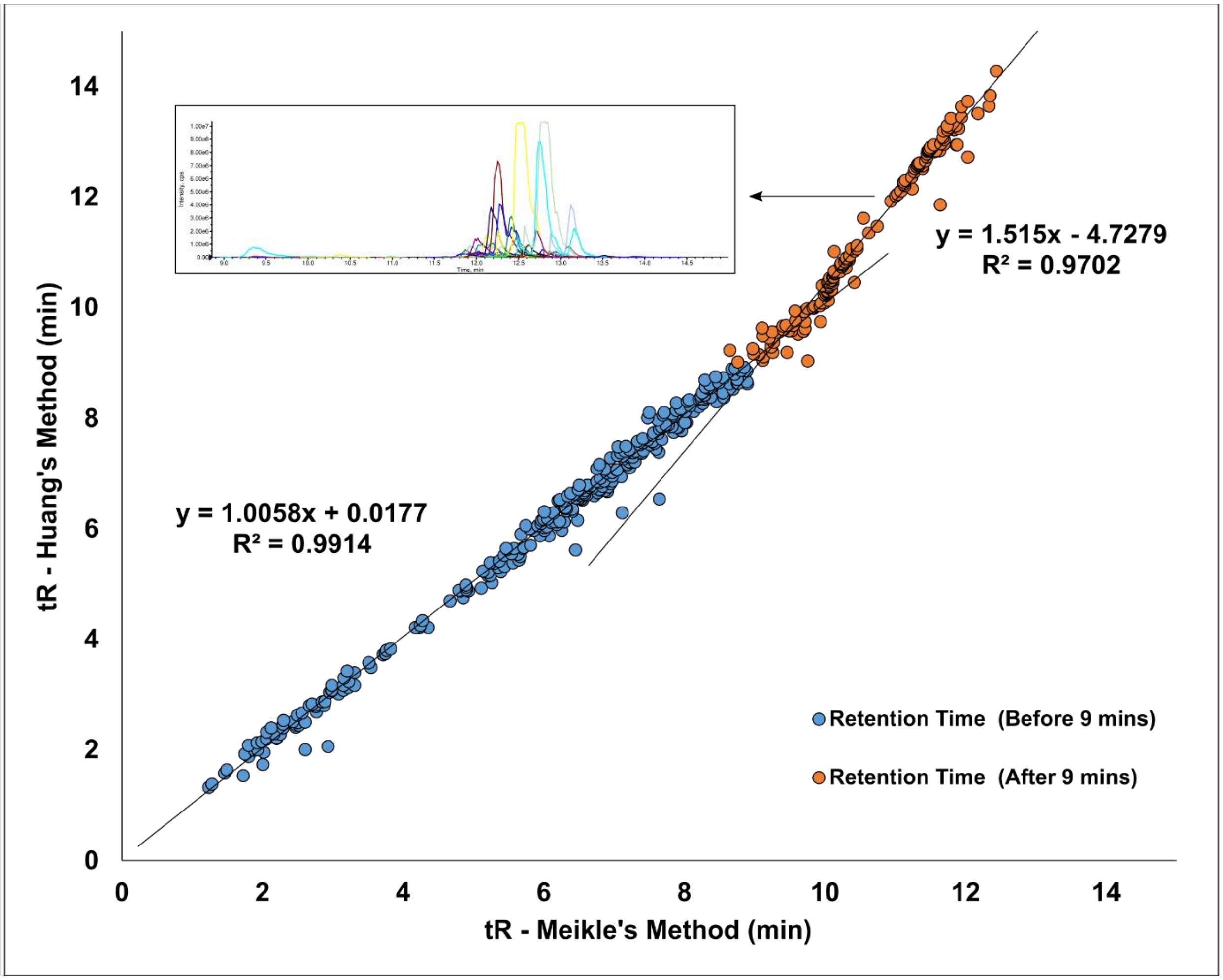

Supplemental Figure 1: Retention time comparison between original and modified method.

Lipids retention time were marked as dot with X axis was the Huynh's method and Y axis was modified method. Blue dots were lipids eluted before 9 minutes and orange dots were lipid eluted after 9 minutes. Solid line was the linearity before 9 minutes and dash line was after 9 minutes. Linear coefficient and goodness of fit ($R^2$) were indicated on the side.

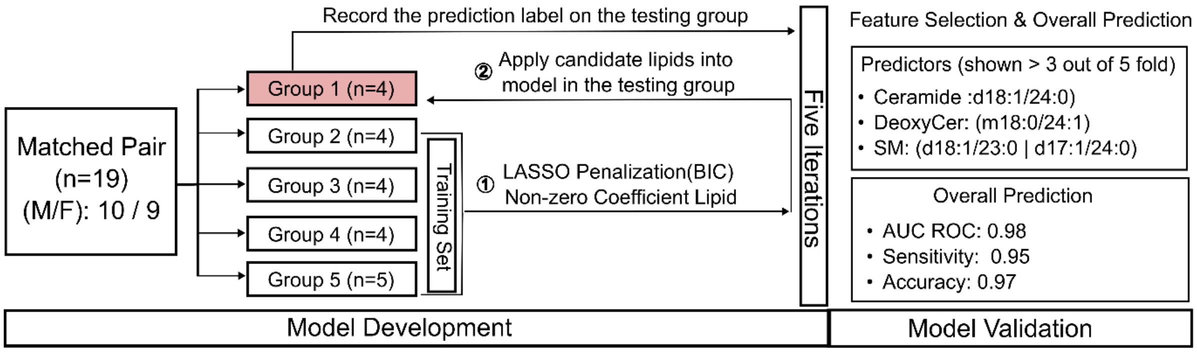

Supplemental Figure 2. Workflow of statistical analysis for lipid predictor discovery.

Pair-matched participants (n = 38; 19 pairs stratified by sex) were randomly divided into five groups. In each iteration, four groups were used as the training set, where generalized regression with LASSO penalization (BIC) identified lipids with non-zero coefficients. The selected lipids were then applied to the held-out group to generate predicted class labels. This procedure was repeated across all folds so that each pair was used once as a test set. Model validation was performed by aggregating predictions across folds to compute overall accuracy, and confusion matrix. Robust predictors were defined as lipids consistently selected across the five iterations (>3 times)

Supplemental Table 1: Participant characteristics stratified by LDLc risk group and sex

| | | Low Risk (n= 10) | High Risk (n= 10) | p-value | Low Risk (n= 9) | High Risk (n= 9) | p-value |
|---|---|---|---|---|---|---|---|
| | Units | Male (n= 10/grp) | | | Female (n= 9/grp) | | |
| Age | y | 45.5 ± 13.4 | 45.5 ± 13.6 | | 56.1 ± 8.7 | 55.8 ± 7.9 | |
| BMI | kg/m² | 25.2 ± 4.1 | 30.2 ± 3.1 | <0.01 | 26.6 ± 6.8 | 26.9 ± 4.0 | 0.9 |
| LDLc | mg/dL | 89.3 ± 11.2 | 174 ± 8 | <0.01 | 91.1 ± 12.0 | 179 ± 20 | <0.01 |
| HDL-C | mg/dL | 53.9 ± 10.9 | 44.8 ± 6.6 | <0.05 | 71.7 ± 15.7 | 59.8 ± 13.0 | 0.07 |
| Total-C | mg/dL | 150 ± 17 | 237 ± 15 | <0.01 | 169 ± 14 | 256 ± 27 | <0.01 |
| Glucose | mg/dL | 93.1 ± 12.9 | 99.9 ± 14.5 | 0.28 | 93.5 ± 10.6 | 97.2 ± 9.0 | 0.43 |
| NEFA | mg/dL | 0.20 ± 0.07 | 0.38 ± 0.15 | <0.01 | 0.30 ± 0.08 | 0.41 ± 0.15 | 0.07 |
| TG | mg/dL | 81.7 ± 28.8 | 147 ± 44 | <0.01 | 79.7 ± 38.3 | 148 ± 74 | <0.02 |

**Supplemental Table 2: Surrogate Standards**

| order | SSTD Stock Solution | Lipid name | cat # | Supplier | Purchased | Molecular weight (mg/mmol) |
|---|---|---|---|---|---|---|
| 1 | d7-PC_(A) | PC(15:0_18:1) d7 | 791637 | Avanti | solution | 753.09 |
| 2 | d7-PE_(A) | PE(15:0_18:1) d7 | 791638 | Avanti | solution | 711.01 |
| 3 | d7-PS_(A) | PS(15:0_18:1) d7 | 791639 | Avanti | solution | 777 |
| 4 | d7-PG_(A) | PG(15:0_18:1) d7 | 791640 | Avanti | solution | 764.01 |
| 5 | d7-PI_(A) | PI(15:0_18:1) d7 | 791641 | Avanti | solution | 847.12 |
| 6 | d7-PA_(A) | PA(15:0_18:1) d7 | 791642 | Avanti | solution | 689.93 |
| 7 | d9-PE_(A) | PE(P-18:0/18:1) d9 | 852474 | Avanti | solution | 739.11 |
| 8 | d9-PC_(A) | PC(P-18:0/18:1) d9 | 852475 | Avanti | solution | 781.19 |
| 9 | d7-LPC_(A) | LPC(18:1) d7 | 791643 | Avanti | solution | 528.71 |
| 10 | d7-LPE_(A) | LPE(18:1) d7 | 791644 | Avanti | solution | 486.63 |
| 11 | LPI(C13:0)_(A) | LPI 13:0 | 850101 | Avanti | neat | 547.57 |
| 12 | d5-GM3_(A) | GM3(d18:1/18:0) d5 | 860073 | Avanti | neat | 1203.55 |
| 13 | d7-HexCer_(A) | HexCer(d18:1/15:0) d7 | 330729 | Avanti | neat | 693.06 |
| 14 | d7-Hex2Cer_(A) | Hex2Cer(d18:1/15:0) d7 | 330727 | Avanti | neat | 855.2 |
| 15 | Hex3Cer(C17)_(A) | Trihexosylceramide 17:0 | 1523 | Matreya LLC | neat | 1038 |
| 16 | AcylCer_(A) | 1-O-Acyl-Ceramide | 860526P | Avanti | neat | 816.37 |
| 17 | d7-dhCer_(A) | dhCer(d18:0/13:0) d7 | 330726 | Avanti | neat | 504.88 |
| 18 | Sph(17:1)_(A) | Sphingosine (Sph) 17:1 base | 860640P | Avanti | neat | 285.47 |
| 19 | Sphn(17:0)_(A) | Sphinganine (Sphn) 17:0 base | 860654P | Avanti | neat | 287.48 |
| 20 | d7-S1P_(A) | S1P d7 | 860659 | Avanti | neat | 386.515 |
| 21 | d9-SM_(A) | SM(d18:1/15:0) d9 | 860686 | Avanti | neat | 698.06 |
| 22 | d6-DesmE_(A) | desmosterol ester d6 | 700184 | Avanti | solution | 655.12 |
| 23 | d7-CholE_(A) | CE 18:1 d7 | 700185 | Avanti | solution | 658.14 |
| 24 | d6-Chol_(A) | cholesterol d6 | 700172 | Avanti | neat | 392.69 |
| 25 | d7-MAG_(A) | MAG 18:1 d7 | 791646 | Avanti | solution | 363.58 |
| 26 | d7-DG_(A) | DG 15:0 18:1 d7 | 791647 | Avanti | solution | 587.97 |
| 27 | d7-TG_(A) | TG 15:0 18:1-d7 15:0 | 791648 | Avanti | solution | 812.35 |
| 28 | d9-OA_(A) | oleic acid d9 | 861809 | Avanti | neat | 291.52 |
| 29 | d11-AA_(A) | arachidonic acid d11 | 861810 | Avanti | neat | 315.53 |
| 30 | d5-DHA_(A) | FA C22:6 d5 | 733326P | Avanti | neat | 333.52 |
| 31 | BMP_(A) | BMP 14:0 14:0 | 857131P | Avanti | neat | 683.89 |
| 32 | d3-AcylCarn_(A) | acylcarnitine 16:0d3 | 71-1746-5 | Larodan | neat | 439.09 |
| 33 | Sulph(18:1/12:0)_(A) | sulphatide d18:1/12:0 | 860573P | Avanti | neat | 741.03 |

**Supplemental Table 3: Transitions, Conditions, Retention Time**

| Lipid # | Lipid Species | Parent Ion | Precursor ion (Q1) | Product ion (Q3) | Approximate Retention Time (min) | Window (sec) | Primary/Secondary | Threshold | Dwell Weight | Declustering Potential | Entrance Potential | Collision Energy | Collision Cell Exit Potential |
|---|---|---|---|---|---|---|---|---|---|---|---|---|---|
| 1 | 1-_&_2-AG | $[M+NH_4]^+$ | 379.302 | 287.2 | 3.3 | 24 | 1 | 0 | 2 | 95 | 10 | 21 | 7 |
| 2 | 1-_&_2-LG | $[M+NH_4]^+$ | 355.202 | 263.2 | 3.2 | 24 | 1 | 0 | 1 | 40 | 8 | 24 | 10 |
| 3 | 1-_&_2-OG | $[M+NH_4]^+$ | 357.202 | 265.2 | 3.6 | 30 | 1 | 0 | 1 | 55 | 4 | 34 | 25 |
| 4 | Acylcarnitine(12:0) | $[M+H]^+$ | 344.3 | 85.1 | 1.25 | 18 | 1 | 0 | 1 | 148 | 5 | 22 | 9 |
| 5 | Acylcarnitine(13:0) | $[M+H]^+$ | 358.3 | 85.1 | 1.4 | 18 | 1 | 0 | 5 | 148 | 10 | 20 | 12 |
| 6 | Acylcarnitine(14:0) | $[M+H]^+$ | 372.3 | 85.1 | 1.82 | 18 | 1 | 0 | 5 | 148 | 8 | 60 | 8 |
| 7 | Acylcarnitine(14:1) | $[M+H]^+$ | 370.3 | 85.1 | 1.5 | 18 | 1 | 0 | 5 | 148 | 8 | 60 | 8 |
| 8 | Acylcarnitine(14:2) | $[M+H]^+$ | 368.3 | 85.1 | 1.18 | 18 | 1 | 0 | 5 | 148 | 8 | 60 | 8 |
| 9 | Acylcarnitine(15:0) (a) | $[M+H]^+$ | 386.3 | 85.1 | 1.95 | 18 | 1 | 0 | 5 | 148 | 5 | 11 | 9 |
| 10 | Acylcarnitine(16:0) | $[M+H]^+$ | 400.4 | 85.1 | 2.5 | 18 | 1 | 0 | 1 | 148 | 8 | 60 | 8 |
| 11 | Acylcarnitine(16:1) | $[M+H]^+$ | 398.3 | 85.1 | 2.05 | 18 | 1 | 0 | 5 | 148 | 5 | 11 | 9 |
| 12 | Acylcarnitine(17:0) (a) | $[M+H]^+$ | 414.4 | 85.1 | 2.7 | 18 | 1 | 0 | 5 | 148 | 5 | 8 | 5 |
| 13 | Acylcarnitine(18:0) | $[M+H]^+$ | 428.4 | 85.1 | 3.12 | 24 | 1 | 0 | 5 | 148 | 8 | 34 | 15 |
| 14 | Acylcarnitine(18:1) | $[M+H]^+$ | 426.4 | 85.1 | 2.67 | 18 | 1 | 0 | 1 | 148 | 5 | 8 | 5 |
| 15 | Acylcarnitine(18:2) | $[M+H]^+$ | 424.3 | 85.1 | 2.22 | 18 | 1 | 0 | 5 | 148 | 8 | 34 | 15 |
| 16 | CE(14:0) | $[M+NH_4]^+$ | 614.6 | 369.3 | 12.65 | 24 | 1 | 0 | 5 | 9 | 5 | 11 | 9 |
| 17 | CE(15:0) | $[M+NH_4]^+$ | 628.6 | 369.3 | 12.9 | 18 | 1 | 0 | 5 | 9 | 8 | 60 | 8 |
| 18 | CE(16:0) | $[M+NH_4]^+$ | 642.6 | 369.3 | 12.5 | 30 | 1 | 0 | 5 | 9 | 8 | 34 | 15 |
| 19 | CE(16:1) | $[M+NH_4]^+$ | 640.6 | 369.3 | 12.2 | 24 | 1 | 0 | 5 | 9 | 8 | 60 | 8 |
| 20 | CE(16:2) | $[M+NH_4]^+$ | 638.6 | 369.3 | 12.3 | 24 | 1 | 0 | 5 | 9 | 8 | 34 | 15 |
| 21 | CE(17:0) | $[M+NH_4]^+$ | 656.6 | 369.3 | 13.2 | 24 | 1 | 0 | 5 | 9 | 8 | 34 | 15 |
| 22 | CE(17:1) | $[M+NH_4]^+$ | 654.6 | 369.3 | 12.8 | 24 | 1 | 0 | 5 | 9 | 6 | 17 | 5 |
| 23 | CE(18:0) | $[M+NH_4]^+$ | 670.7 | 369.3 | 13.55 | 24 | 1 | 0 | 5 | 9 | 8 | 60 | 8 |
| 24 | CE(18:1) | $[M+NH_4]^+$ | 668.6 | 369.3 | 12.65 | 24 | 1 | 0 | 5 | 9 | 8 | 34 | 15 |
| 25 | CE(18:2) | $[M+NH_4]^+$ | 666.6 | 369.3 | 12.3 | 24 | 1 | 0 | 5 | 9 | 8 | 34 | 15 |
| 26 | CE(18:3) | $[M+NH_4]^+$ | 664.6 | 369.3 | 11.95 | 24 | 1 | 0 | 5 | 9 | 8 | 34 | 15 |
| 27 | CE(20:1) | $[M+NH_4]^+$ | 696.7 | 369.3 | 12 | 30 | 1 | 0 | 5 | 9 | 8 | 34 | 15 |
| 28 | CE(20:2) | $[M+NH_4]^+$ | 694.7 | 369.3 | 13.1 | 24 | 1 | 0 | 5 | 9 | 8 | 34 | 15 |
| 29 | CE(20:3) | $[M+NH_4]^+$ | 692.6 | 369.3 | 12.3 | 24 | 1 | 0 | 5 | 9 | 9 | 27 | 15 |
| 30 | CE(20:4) | $[M+NH_4]^+$ | 690.6 | 369.3 | 12.05 | 24 | 1 | 0 | 5 | 9 | 5 | 11 | 9 |
| 31 | CE(20:5) | $[M+NH_4]^+$ | 688.6 | 369.3 | 11.7 | 30 | 1 | 0 | 5 | 9 | 8 | 60 | 8 |
| 32 | CE(22:0) | $[M+NH_4]^+$ | 726.7 | 369.3 | 13.8 | 24 | 1 | 0 | 5 | 9 | 9 | 27 | 15 |
| 33 | CE(22:1) | $[M+NH_4]^+$ | 724.7 | 369.3 | 13.4 | 24 | 1 | 0 | 5 | 9 | 9 | 27 | 15 |
| 34 | CE(22:4) | $[M+NH_4]^+$ | 718.7 | 369.3 | 12.85 | 24 | 1 | 0 | 5 | 9 | 8 | 34 | 15 |
| 35 | CE(22:5) (n3) | $[M+NH_4]^+$ | 716.6 | 369.3 | 11.8 | 36 | 1 | 0 | 5 | 9 | 8 | 34 | 15 |
| 36 | CE(22:6) | $[M+NH_4]^+$ | 714.6 | 369.3 | 11.85 | 30 | 1 | 0 | 5 | 9 | 4 | 60 | 8 |
| 37 | CE(24:0) | $[M+NH_4]^+$ | 754.7 | 369.3 | 14.2 | 24 | 1 | 0 | 5 | 9 | 8 | 34 | 15 |
| 38 | CE(24:1) | $[M+NH_4]^+$ | 752.7 | 369.3 | 13.8 | 24 | 1 | 0 | 5 | 9 | 5 | 8 | 5 |
| 39 | CE(24:4) | $[M+NH_4]^+$ | 746.7 | 369.3 | 13.2 | 24 | 1 | 0 | 5 | 9 | 8 | 34 | 15 |
| 40 | CE(24:5) | $[M+NH_4]^+$ | 744.7 | 369.3 | 12.6 | 24 | 1 | 0 | 5 | 9 | 8 | 60 | 8 |
| 41 | CE(24:6) | $[M+NH_4]^+$ | 742.7 | 369.3 | 12.62 | 24 | 1 | 0 | 5 | 9 | 8 | 34 | 15 |
| 42 | Cer(d16:1_16:0) | $[M+H]^+$ | 510.6 | 236.3 | 6.65 | 24 | 1 | 0 | 5 | 90 | 8 | 34 | 15 |
| 43 | Cer(d16:1_18:0) | $[M+H]^+$ | 538.6 | 236.3 | 7.85 | 24 | 1 | 0 | 5 | 90 | 9 | 27 | 15 |
| 44 | Cer(d16:1_20:0) | $[M+H]^+$ | 566.6 | 236.3 | 9.15 | 24 | 1 | 0 | 5 | 90 | 8 | 34 | 15 |
| 45 | Cer(d16:1_22:0) | $[M+H]^+$ | 594.6 | 236.3 | 10.2 | 24 | 1 | 0 | 5 | 90 | 8 | 34 | 15 |
| 46 | Cer(d16:1_23:0) | $[M+H]^+$ | 608.6 | 236.3 | 10.37 | 24 | 1 | 0 | 5 | 90 | 8 | 60 | 8 |
| 47 | Cer(d16:1_24:0) | $[M+H]^+$ | 622.6 | 236.3 | 10.6 | 24 | 1 | 0 | 5 | 90 | 8 | 60 | 8 |
| 48 | Cer(d16:1_24:1) | $[M+H]^+$ | 620.6 | 236.3 | 10.2 | 24 | 1 | 0 | 5 | 90 | 9 | 27 | 15 |
| 49 | Cer(d17:1_16:0) | $[M+H]^+$ | 524.6 | 250.3 | 7.25 | 18 | 1 | 0 | 5 | 90 | 9 | 27 | 15 |
| 50 | Cer(d17:1_18:0) | $[M+H]^+$ | 552.6 | 250.3 | 8.4 | 24 | 1 | 0 | 5 | 90 | 8 | 34 | 15 |
| 51 | Cer(d17:1_20:0) | $[M+H]^+$ | 580.6 | 250.3 | 9.75 | 24 | 1 | 0 | 5 | 90 | 8 | 34 | 15 |
| 52 | Cer(d17:1_22:0) | $[M+H]^+$ | 608.6 | 250.3 | 10.8 | 24 | 1 | 0 | 5 | 90 | 8 | 34 | 15 |
| 53 | Cer(d17:1_23:0) | $[M+H]^+$ | 622.6 | 250.3 | 10.55 | 24 | 1 | 0 | 5 | 90 | 8 | 34 | 15 |
| 54 | Cer(d17:1_24:0) | $[M+H]^+$ | 636.6 | 250.3 | 10.75 | 24 | 1 | 0 | 5 | 90 | 8 | 34 | 15 |
| 55 | Cer(d17:1_24:1) | $[M+H]^+$ | 634.6 | 250.3 | 10.4 | 24 | 1 | 0 | 5 | 90 | 8 | 34 | 15 |
| 56 | Cer(d18:1_14:0) | $[M+H]^+$ | 510.5 | 264.3 | 6.7 | 24 | 1 | 0 | 5 | 90 | 8 | 34 | 15 |
| 57 | Cer(d18:1_16:0) | $[M+H]^+$ | 538.5 | 264.3 | 7.8 | 24 | 1 | 0 | 5 | 90 | 8 | 34 | 15 |
| 58 | Cer(d18:1_17:0) | $[M+H]^+$ | 552.5 | 264.3 | 8.4 | 24 | 1 | 0 | 5 | 90 | 9 | 27 | 15 |
| 59 | Cer(d18:1_18:0) | $[M+H]^+$ | 566.6 | 264.3 | 9.1 | 24 | 1 | 0 | 5 | 90 | 9 | 27 | 15 |
| 60 | Cer(d18:1_19:0) | $[M+H]^+$ | 580.6 | 264.3 | 9.4 | 24 | 1 | 0 | 5 | 90 | 9 | 27 | 15 |
| 61 | Cer(d18:1_20:0) | $[M+H]^+$ | 594.6 | 264.3 | 10.15 | 24 | 1 | 0 | 5 | 90 | 8 | 60 | 8 |
| 62 | Cer(d18:1_21:0) | $[M+H]^+$ | 608.6 | 264.3 | 10.35 | 24 | 1 | 0 | 5 | 90 | 8 | 34 | 15 |

| | | | | | | | | | | | | | |
|---|---|---|---|---|---|---|---|---|---|---|---|---|---|
| 63 | Cer(d18:1_22:0) | $[M+H]^+$ | 622.6 | 264.3 | 10.55 | 24 | 1 | 0 | 5 | 90 | 8 | 34 | 15 |
| 64 | Cer(d18:1_23:0) | $[M+H]^+$ | 636.6 | 264.3 | 10.7 | 24 | 1 | 0 | 5 | 90 | 10 | 12 | 12 |
| 65 | Cer(d18:1_24:0) | $[M+H]^+$ | 650.6 | 264.3 | 10.85 | 24 | 1 | 0 | 5 | 90 | 8 | 34 | 15 |
| 66 | Cer(d18:1_24:1) | $[M+H]^+$ | 648.6 | 264.3 | 10.55 | 24 | 1 | 0 | 5 | 90 | 8 | 34 | 15 |
| 67 | Cer(d18:1_26:0) | $[M+H]^+$ | 678.6 | 264.3 | 10.5 | 36 | 1 | 0 | 1 | 90 | 9 | 27 | 15 |
| 68 | Cer(d18:2_14:0) | $[M+H]^+$ | 508.5 | 262.3 | 5.85 | 24 | 1 | 0 | 5 | 90 | 10 | 21 | 12 |
| 69 | Cer(d18:2_16:0) | $[M+H]^+$ | 536.5 | 262.3 | 6.85 | 24 | 1 | 0 | 5 | 90 | 8 | 34 | 15 |
| 70 | Cer(d18:2_17:0) | $[M+H]^+$ | 550.5 | 262.3 | 7.1 | 36 | 1 | 0 | 5 | 90 | 8 | 34 | 15 |
| 71 | Cer(d18:2_18:0) | $[M+H]^+$ | 564.6 | 262.3 | 8.1 | 24 | 1 | 0 | 5 | 90 | 9 | 27 | 15 |
| 72 | Cer(d18:2_20:0) | $[M+H]^+$ | 592.6 | 262.3 | 9.45 | 24 | 1 | 0 | 5 | 90 | 10 | 15 | 12 |
| 73 | Cer(d18:2_21:0) | $[M+H]^+$ | 606.6 | 262.3 | 9.45 | 36 | 1 | 0 | 5 | 90 | 14 | 21 | 14 |
| 74 | Cer(d18:2_22:0) | $[M+H]^+$ | 620.6 | 262.3 | 10.25 | 24 | 1 | 0 | 5 | 90 | 8 | 34 | 15 |
| 75 | Cer(d18:2_23:0) | $[M+H]^+$ | 634.6 | 262.3 | 10.5 | 24 | 1 | 0 | 5 | 90 | 8 | 34 | 15 |
| 76 | Cer(d18:2_24:0) | $[M+H]^+$ | 648.6 | 262.3 | 10.65 | 24 | 1 | 0 | 5 | 90 | 5 | 45 | 5 |
| 77 | Cer(d18:2_24:1) | $[M+H]^+$ | 646.6 | 262.3 | 10.3 | 24 | 1 | 0 | 5 | 90 | 8 | 34 | 15 |
| 78 | Cer(d18:2_26:0) | $[M+H]^+$ | 676.6 | 262.3 | 10.8 | 24 | 1 | 0 | 5 | 90 | 8 | 34 | 15 |
| 79 | Cer(d19:1_16:0) | $[M+H]^+$ | 552.6 | 278.3 | 8.2 | 24 | 1 | 0 | 5 | 90 | 9 | 27 | 15 |
| 80 | Cer(d19:1_18:0) | $[M+H]^+$ | 580.6 | 278.3 | 9.1 | 24 | 1 | 0 | 5 | 90 | 8 | 34 | 15 |
| 81 | Cer(d19:1_20:0) | $[M+H]^+$ | 608.6 | 278.3 | 10 | 24 | 1 | 0 | 5 | 90 | 8 | 34 | 15 |
| 82 | Cer(d19:1_22:0) | $[M+H]^+$ | 636.6 | 278.3 | 10.65 | 24 | 1 | 0 | 5 | 90 | 5 | 32 | 10 |
| 83 | Cer(d19:1_23:0) | $[M+H]^+$ | 650.6 | 278.3 | 10.8 | 24 | 1 | 0 | 5 | 90 | 4 | 57 | 25 |
| 84 | Cer(d19:1_24:0) | $[M+H]^+$ | 664.6 | 278.3 | 10.9 | 24 | 1 | 0 | 1 | 90 | 8 | 34 | 15 |
| 85 | Cer(d19:1_24:1) | $[M+H]^+$ | 662.6 | 278.3 | 10.7 | 24 | 1 | 0 | 5 | 90 | 8 | 34 | 15 |
| 86 | Cer(d19:1_26:0) | $[M+H]^+$ | 692.6 | 278.3 | 11 | 24 | 1 | 0 | 5 | 90 | 4 | 9 | 25 |
| 87 | Cer(d20:1_22:0) | $[M+H]^+$ | 650.6 | 292.3 | 10.2 | 48 | 1 | 0 | 5 | 90 | 5 | 32 | 10 |
| 88 | Cer(d20:1_23:0) | $[M+H]^+$ | 664.6 | 292.3 | 10.4 | 48 | 1 | 0 | 5 | 90 | 5 | 32 | 10 |
| 89 | Cer(d20:1_24:0) | $[M+H]^+$ | 678.6 | 292.3 | 10.6 | 48 | 1 | 0 | 5 | 90 | 5 | 45 | 5 |
| 90 | Cer(d20:1_24:1) | $[M+H]^+$ | 676.6 | 292.3 | 10.6 | 24 | 1 | 0 | 5 | 90 | 12 | 41 | 36 |
| 91 | Cer(d20:1_26:0) | $[M+H]^+$ | 706.6 | 292.3 | 11 | 24 | 1 | 0 | 5 | 90 | 8 | 34 | 15 |
| 92 | Cer(m18:0_20:0) | $[M+H]^+$ | 580.6 | 268.4 | 10.53 | 24 | 1 | 0 | 5 | 90 | 5 | 48 | 32 |
| 93 | Cer(m18:0_22:0) | $[M+H]^+$ | 608.6 | 268.4 | 10.82 | 24 | 1 | 0 | 5 | 90 | 5 | 45 | 5 |
| 94 | Cer(m18:0_23:0) | $[M+H]^+$ | 622.6 | 268.4 | 11 | 24 | 1 | 0 | 5 | 90 | 12 | 41 | 36 |
| 95 | Cer(m18:0_24:0) | $[M+H]^+$ | 636.6 | 268.4 | 11.05 | 36 | 1 | 0 | 5 | 90 | 5 | 48 | 32 |
| 96 | Cer(m18:0_24:1) | $[M+H]^+$ | 634.6 | 268.4 | 10.85 | 24 | 1 | 0 | 5 | 90 | 5 | 32 | 10 |
| 97 | Cer(m18:1_18:0) | $[M+H]^+$ | 550.6 | 266.4 | 9.5 | 36 | 1 | 0 | 5 | 90 | 5 | 48 | 32 |
| 98 | Cer(m18:1_20:0) | $[M+H]^+$ | 578.6 | 266.4 | 10.15 | 36 | 1 | 0 | 5 | 90 | 4 | 56 | 12 |
| 99 | Cer(m18:1_22:0) | $[M+H]^+$ | 606.6 | 266.4 | 10.5 | 36 | 1 | 0 | 5 | 90 | 5 | 48 | 32 |
| 100 | Cer(m18:1_23:0) | $[M+H]^+$ | 620.6 | 266.4 | 10.7 | 36 | 1 | 0 | 5 | 90 | 5 | 48 | 32 |
| 101 | Cer(m18:1_24:0) | $[M+H]^+$ | 634.6 | 266.4 | 10.85 | 24 | 1 | 0 | 5 | 90 | 12 | 41 | 36 |
| 102 | Cer(m18:1_24:1) | $[M+H]^+$ | 632.6 | 266.4 | 10.5 | 36 | 1 | 0 | 5 | 90 | 5 | 32 | 10 |
| 103 | Cer1P(d18:1_16:0) | $[M+H]^+$ | 618.4 | 264.3 | 10.6 | 36 | 1 | 0 | 1 | 25 | 4 | 56 | 12 |
| 104 | COH (cholesterol) | $[M+NH_4]^+$ | 369.4 | 161.2 | 6.35 | 30 | 1 | 0 | 1 | 9 | 12 | 41 | 36 |
| 105 | dCer(d18:0_16:0) | $[M+H]^+$ | 540.5 | 284.3 | 8.2 | 48 | 1 | 0 | 5 | 90 | 12 | 41 | 36 |
| 106 | dCer(d18:0_18:0) | $[M+H]^+$ | 568.6 | 284.3 | 9.6 | 36 | 1 | 0 | 5 | 90 | 10 | 29 | 16 |
| 107 | dCer(d18:0_20:0) | $[M+H]^+$ | 596.6 | 284.3 | 10.3 | 30 | 1 | 0 | 5 | 90 | 5 | 32 | 10 |
| 108 | dCer(d18:0_22:0) | $[M+H]^+$ | 624.6 | 284.3 | 10.7 | 18 | 1 | 0 | 5 | 90 | 4 | 57 | 25 |
| 109 | dCer(d18:0_24:0) | $[M+H]^+$ | 652.7 | 284.3 | 10.9 | 24 | 1 | 0 | 5 | 90 | 5 | 48 | 32 |
| 110 | dCer(d18:0_24:1) | $[M+H]^+$ | 650.6 | 284.3 | 10.7 | 24 | 1 | 0 | 5 | 90 | 5 | 48 | 32 |
| 111 | DE(16:0) | $[M+H]^+$ | 640.8 | 367.4 | 11.5 | 48 | 1 | 0 | 5 | 5 | 5 | 32 | 10 |
| 112 | DE(18:1) | $[M+H]^+$ | 666.8 | 367.4 | 12.6 | 36 | 1 | 0 | 5 | 5 | 5 | 32 | 10 |
| 113 | DE(18:2) | $[M+H]^+$ | 664.8 | 367.4 | 12.3 | 36 | 1 | 0 | 5 | 5 | 4 | 56 | 12 |
| 114 | DE(20:4) | $[M+H]^+$ | 688.8 | 367.4 | 12.2 | 24 | 1 | 0 | 1 | 5 | 10 | 29 | 16 |
| 115 | DE(20:5) | $[M+H]^+$ | 686.8 | 367.4 | 11.8 | 36 | 1 | 0 | 5 | 5 | 5 | 32 | 10 |
| 116 | DE(22:6) | $[M+H]^+$ | 712.8 | 367.4 | 11.05 | 30 | 1 | 0 | 1 | 5 | 8 | 34 | 15 |
| 117 | DG(14:0_16:0) | $[M+H]^+$ | 558.5 | 313.3 | 8.9 | 24 | 1 | 0 | 1 | 5 | 12 | 41 | 36 |
| 118 | DG(14:0_18:2) | $[M+H]^+$ | 582.5 | 285.2 | 8.25 | 24 | 1 | 0 | 5 | 5 | 12 | 41 | 36 |
| 119 | DG(16:0_16:0) | $[M+H]^+$ | 586.5 | 313.3 | 9.65 | 24 | 1 | 0 | 5 | 5 | 12 | 41 | 36 |
| 120 | DG(16:0_16:1) | $[M+H]^+$ | 584.5 | 313.3 | 8.9 | 36 | 1 | 0 | 5 | 5 | 5 | 48 | 32 |
| 121 | DG(16:0_18:1) | $[M+H]^+$ | 612.6 | 313.3 | 10.15 | 24 | 1 | 0 | 1 | 5 | 12 | 41 | 36 |
| 122 | DG(16:0_18:2) | $[M+H]^+$ | 610.5 | 313.3 | 9.25 | 42 | 1 | 0 | 5 | 5 | 12 | 41 | 36 |
| 123 | DG(16:0_20:4) | $[M+H]^+$ | 634.5 | 313.3 | 9.05 | 42 | 1 | 0 | 5 | 5 | 5 | 48 | 32 |
| 124 | DG(16:0_22:5) | $[M+H]^+$ | 660.6 | 313.3 | 9.2 | 24 | 1 | 0 | 5 | 5 | 4 | 56 | 12 |
| 125 | DG(16:0_22:6) | $[M+H]^+$ | 658.5 | 313.3 | 9.1 | 24 | 1 | 0 | 5 | 5 | 10 | 25 | 12 |
| 126 | DG(16:1_18:1) | $[M+H]^+$ | 610.5 | 339.3 | 9.2 | 30 | 1 | 0 | 1 | 5 | 5 | 48 | 32 |
| 127 | DG(18:0_18:1) | $[M+H]^+$ | 640.6 | 341.3 | 10.1 | 60 | 1 | 0 | 5 | 5 | 5 | 48 | 32 |
| 128 | DG(18:0_18:2) | $[M+H]^+$ | 638.6 | 341.3 | 10.3 | 24 | 1 | 0 | 5 | 5 | 5 | 32 | 10 |
| 129 | DG(18:0_20:4) | $[M+H]^+$ | 662.6 | 341.3 | 10.27 | 24 | 1 | 0 | 5 | 5 | 10 | 25 | 12 |
| 130 | DG(18:0_22:6) | $[M+H]^+$ | 686.6 | 341.3 | 9.35 | 24 | 1 | 0 | 5 | 5 | 5 | 48 | 32 |
| 131 | DG(18:1_18:1) | $[M+H]^+$ | 638.6 | 339.3 | 9.75 | 30 | 1 | 0 | 5 | 5 | 5 | 48 | 32 |
| 132 | DG(18:1_18:2) | $[M+H]^+$ | 636.6 | 339.3 | 9.5 | 36 | 1 | 0 | 1 | 5 | 12 | 41 | 36 |
| 133 | DG(18:1_18:3) | $[M+H]^+$ | 634.5 | 339.2 | 8.8 | 48 | 1 | 0 | 5 | 5 | 12 | 41 | 36 |
| 134 | DG(18:1_20:3) | $[M+H]^+$ | 662.6 | 339.3 | 9.5 | 36 | 1 | 0 | 5 | 5 | 12 | 41 | 36 |
| 135 | DG(18:1_20:4) | $[M+H]^+$ | 660.6 | 339.3 | 9 | 24 | 1 | 0 | 5 | 5 | 5 | 48 | 32 |

| | | | | | | | | | | | | | |
|---|---|---|---|---|---|---|---|---|---|---|---|---|---|
| 136 | DG(18:1_20:5) | $[M+H]^+$ | 658.6 | 339.3 | 8.3 | 24 | 1 | 0 | 5 | 5 | 12 | 41 | 36 |
| 137 | DG(18:1_22:5) | $[M+H]^+$ | 686.6 | 339.3 | 9.5 | 60 | 1 | 0 | 5 | 5 | 5 | 32 | 10 |
| 138 | DG(18:1_22:6) | $[M+H]^+$ | 684.6 | 339.3 | 8.8 | 24 | 1 | 0 | 5 | 5 | 7 | 43 | 14 |
| 139 | DG(18:2_18:2) | $[M+H]^+$ | 634.5 | 337.2 | 8.3 | 24 | 1 | 0 | 1 | 5 | 12 | 41 | 36 |
| 140 | DG(18:2_20:4) | $[M+H]^+$ | 658.5 | 337.2 | 8.65 | 24 | 1 | 0 | 5 | 5 | 12 | 41 | 36 |
| 141 | DG(18:2_22:6) | $[M+H]^+$ | 682.6 | 337.3 | 8.37 | 24 | 1 | 0 | 5 | 5 | 5 | 48 | 32 |
| 142 | GM1(d18:1_16:0) | $[M+2H]^{2+}$ | 760.1 | 366.3 | 4.65 | 18 | 1 | 0 | 5 | 12 | 8 | 23 | 9 |
| 143 | GM3(d18:1_16:0) | $[M+H]^+$ | 1153.7 | 264.3 | 4.6 | 24 | 1 | 0 | 5 | 12 | 4 | 56 | 12 |
| 144 | GM3(d18:1_18:0) | $[M+H]^+$ | 1181.8 | 264.3 | 5.3 | 18 | 1 | 0 | 5 | 12 | 10 | 44 | 7 |
| 145 | GM3(d18:1_20:0) | $[M+H]^+$ | 1209.8 | 264.3 | 6.05 | 24 | 1 | 0 | 5 | 12 | 8 | 34 | 15 |
| 146 | GM3(d18:1_22:0) | $[M+H]^+$ | 1237.8 | 264.3 | 7.05 | 24 | 1 | 0 | 5 | 12 | 12 | 41 | 36 |
| 147 | GM3(d18:1_24:0) | $[M+H]^+$ | 1265.8 | 264.3 | 8.15 | 24 | 1 | 0 | 5 | 12 | 13 | 32 | 14 |
| 148 | GM3(d18:1_24:1) | $[M+H]^+$ | 1263.8 | 264.3 | 7.15 | 24 | 1 | 0 | 5 | 12 | 12 | 41 | 36 |
| 149 | GM3(d18:2_24:1) | $[M+H]^+$ | 1261.8 | 262.3 | 6.35 | 24 | 1 | 0 | 5 | 12 | 12 | 41 | 36 |
| 150 | GM3(d20:1_18:0) | $[M+H]^+$ | 1209.8 | 292.3 | 6.05 | 18 | 1 | 0 | 5 | 12 | 10 | 29 | 16 |
| 151 | H2Cer(d16:1_16:0) | $[M+H]^+$ | 834.6 | 236.3 | 5.05 | 24 | 1 | 0 | 5 | 90 | 5 | 45 | 5 |
| 152 | H2Cer(d16:1_24:1) | $[M+H]^+$ | 944.7 | 236.3 | 8.1 | 24 | 1 | 0 | 5 | 90 | 13 | 32 | 14 |
| 153 | H2Cer(d18:1_16:0) | $[M+H]^+$ | 862.6 | 264.3 | 5.85 | 24 | 1 | 0 | 1 | 90 | 12 | 41 | 36 |
| 154 | H2Cer(d18:1_18:0) | $[M+H]^+$ | 890.7 | 264.3 | 6.7 | 36 | 1 | 0 | 5 | 90 | 5 | 48 | 32 |
| 155 | H2Cer(d18:1_20:0) | $[M+H]^+$ | 918.7 | 264.3 | 7.9 | 24 | 1 | 0 | 5 | 90 | 5 | 32 | 10 |
| 156 | H2Cer(d18:1_22:0) | $[M+H]^+$ | 946.7 | 264.3 | 9.05 | 30 | 1 | 0 | 5 | 90 | 10 | 25 | 12 |
| 157 | H2Cer(d18:1_24:0) | $[M+H]^+$ | 974.8 | 264.3 | 10.13 | 24 | 1 | 0 | 5 | 90 | 4 | 57 | 25 |
| 158 | H2Cer(d18:1_24:1) | $[M+H]^+$ | 972.7 | 264.3 | 9.25 | 24 | 1 | 0 | 5 | 90 | 4 | 57 | 25 |
| 159 | H2Cer(d18:2_16:0) | $[M+H]^+$ | 860.6 | 262.3 | 5.2 | 24 | 1 | 0 | 5 | 90 | 12 | 41 | 36 |
| 160 | H2Cer(d18:2_24:1) | $[M+H]^+$ | 970.7 | 262.3 | 8.25 | 30 | 1 | 0 | 5 | 90 | 10 | 29 | 16 |
| 161 | H3Cer(d18:1_16:0) | $[M+H]^+$ | 1024.7 | 264.3 | 5.5 | 24 | 1 | 0 | 1 | 90 | 12 | 41 | 36 |
| 162 | H3Cer(d18:1_18:0) | $[M+H]^+$ | 1052.7 | 264.3 | 6.4 | 24 | 1 | 0 | 5 | 90 | 5 | 48 | 32 |
| 163 | H3Cer(d18:1_20:0) | $[M+H]^+$ | 1080.7 | 264.3 | 7.45 | 24 | 1 | 0 | 5 | 90 | 5 | 45 | 5 |
| 164 | H3Cer(d18:1_22:0) | $[M+H]^+$ | 1108.8 | 264.3 | 8.6 | 24 | 1 | 0 | 5 | 90 | 5 | 45 | 5 |
| 165 | H3Cer(d18:1_24:0) | $[M+H]^+$ | 1136.8 | 264.3 | 9.8 | 24 | 1 | 0 | 5 | 90 | 12 | 41 | 36 |
| 166 | H3Cer(d18:1_24:1) | $[M+H]^+$ | 1134.8 | 264.3 | 8.6 | 24 | 1 | 0 | 5 | 90 | 12 | 41 | 36 |
| 167 | HCer(d16:1_18:0) | $[M+H]^+$ | 700.6 | 236.3 | 6.45 | 24 | 1 | 0 | 1 | 30 | 12 | 41 | 36 |
| 168 | HCer(d16:1_20:0) | $[M+H]^+$ | 728.6 | 236.3 | 7.55 | 24 | 1 | 0 | 5 | 30 | 5 | 48 | 32 |
| 169 | HCer(d16:1_22:0) | $[M+H]^+$ | 756.7 | 236.3 | 8.55 | 30 | 1 | 0 | 5 | 30 | 5 | 32 | 10 |
| 170 | HCer(d16:1_24:0) | $[M+H]^+$ | 784.7 | 236.3 | 10 | 24 | 1 | 0 | 5 | 30 | 5 | 32 | 10 |
| 171 | HCer(d18:1_16:0) | $[M+H]^+$ | 700.6 | 264.3 | 6.9 | 24 | 1 | 0 | 1 | 30 | 5 | 32 | 10 |
| 172 | HCer(d18:1_18:0) | $[M+H]^+$ | 728.6 | 264.3 | 7.25 | 30 | 1 | 0 | 5 | 30 | 12 | 41 | 36 |
| 173 | HCer(d18:1_20:0) | $[M+H]^+$ | 756.6 | 264.3 | 8.45 | 30 | 1 | 0 | 5 | 30 | 13 | 32 | 14 |
| 174 | HCer(d18:1_22:0) | $[M+H]^+$ | 784.7 | 264.3 | 9.91 | 24 | 1 | 0 | 1 | 30 | 10 | 25 | 12 |
| 175 | HCer(d18:1_24:0) | $[M+H]^+$ | 812.7 | 264.3 | 10.4 | 24 | 1 | 0 | 5 | 30 | 10 | 29 | 16 |
| 176 | HCer(d18:1_24:1) | $[M+H]^+$ | 810.7 | 264.3 | 9.95 | 24 | 1 | 0 | 1 | 30 | 5 | 32 | 10 |
| 177 | HCer(d18:2_18:0) | $[M+H]^+$ | 726.6 | 262.3 | 6.4 | 36 | 1 | 0 | 5 | 30 | 12 | 41 | 36 |
| 178 | HCer(d18:2_20:0) | $[M+H]^+$ | 754.6 | 262.3 | 7.8 | 24 | 1 | 0 | 5 | 30 | 10 | 29 | 16 |
| 179 | HCer(d18:2_22:0) | $[M+H]^+$ | 782.7 | 262.3 | 8.5 | 30 | 1 | 0 | 5 | 30 | 5 | 45 | 5 |
| 180 | HCer(d18:2_24:0) | $[M+H]^+$ | 810.7 | 262.303 | 10.1 | 24 | 1 | 0 | 1 | 30 | 5 | 23 | 19 |
| 181 | LPC(14:0) [sn2] | $[M+H]^+$ | 468.3 | 184.1 | 1.75 | 30 | 1 | 0 | 5 | 36 | 13 | 32 | 14 |
| 182 | LPC(15:0) [sn2] | $[M+H]^+$ | 482.3 | 184.102 | 2.15 | 24 | 1 | 0 | 1 | 36 | 13 | 32 | 14 |
| 183 | LPC(15-MHDA) [sn1] [104_sn1] | $[M+H]^+$ | 510.4 | 104.1 | 2.8 | 60 | 1 | 0 | 1 | 36 | 8 | 23 | 9 |
| 184 | LPC(15-MHDA) [sn1] _ LPC(17:0) [sn2] | $[M+H]^+$ | 510.4 | 184.1 | 2.8 | 24 | 1 | 0 | 1 | 36 | 13 | 32 | 14 |
| 185 | LPC(16:0) [sn2] | $[M+H]^+$ | 496.3 | 184.1 | 2.5 | 24 | 1 | 0 | 1 | 36 | 8 | 23 | 9 |
| 186 | LPC(16:1) [sn2] | $[M+H]^+$ | 494.3 | 184.1 | 1.95 | 36 | 1 | 0 | 1 | 36 | 4 | 56 | 12 |
| 187 | LPC(17:1 ) [sn2] (a) | $[M+H]^+$ | 508.4 | 184.1 | 2.35 | 24 | 1 | 0 | 1 | 36 | 12 | 41 | 36 |
| 188 | LPC(17:1) (a) [sn1] [104_sn1] | $[M+H]^+$ | 508.4 | 104.1 | 2.35 | 18 | 1 | 0 | 1 | 36 | 13 | 32 | 14 |
| 189 | LPC(18:0) [sn2] | $[M+H]^+$ | 524.4 | 184.1 | 3.15 | 24 | 1 | 0 | 1 | 36 | 12 | 41 | 36 |
| 190 | LPC(18:1) [sn2] | $[M+H]^+$ | 522.4 | 184.1 | 2.65 | 24 | 1 | 0 | 1 | 36 | 12 | 41 | 36 |
| 191 | LPC(18:2) [sn2] | $[M+H]^+$ | 520.3 | 184.1 | 2.2 | 24 | 1 | 0 | 1 | 36 | 10 | 29 | 16 |
| 192 | LPC(18:3) (a) [sn1] [104_sn1] | $[M+H]^+$ | 518.3 | 104.1 | 1.95 | 24 | 1 | 0 | 1 | 36 | 5 | 32 | 10 |
| 193 | LPC(18:3) [sn1] (a)_LPC(18:3) [sn2] (b) | $[M+H]^+$ | 518.3 | 184.1 | 1.9 | 24 | 1 | 0 | 1 | 36 | 12 | 41 | 36 |
| 194 | LPC(19:0) (a) [sn1] [104_sn1] | $[M+H]^+$ | 538.4 | 104.1 | 3.4 | 24 | 1 | 0 | 5 | 36 | 13 | 32 | 14 |
| 195 | LPC(19:0) [sn1] (a) _ LPC(19:0) [sn2] (b) | $[M+H]^+$ | 538.4 | 184.1 | 3.15 | 42 | 1 | 0 | 1 | 36 | 13 | 32 | 14 |
| 196 | LPC(19:1) (a) | $[M+H]^+$ | 536.4 | 184.1 | 2.9 | 30 | 1 | 0 | 1 | 36 | 13 | 32 | 14 |
| 197 | LPC(20:0) [sn2] | $[M+H]^+$ | 552.4 | 184.1 | 3.65 | 24 | 1 | 0 | 1 | 36 | 5 | 48 | 32 |
| 198 | LPC(20:1) [sn2] | $[M+H]^+$ | 550.4 | 184.1 | 3.25 | 24 | 1 | 0 | 1 | 36 | 8 | 23 | 9 |
| 199 | LPC(20:2) [sn2] | $[M+H]^+$ | 548.4 | 184.1 | 2.8 | 24 | 1 | 0 | 1 | 36 | 13 | 32 | 14 |
| 200 | LPC(20:3) [sn2] | $[M+H]^+$ | 546.4 | 184.1 | 2.45 | 24 | 1 | 0 | 5 | 36 | 13 | 32 | 14 |
| 201 | LPC(20:4) [sn2] | $[M+H]^+$ | 544.3 | 184.1 | 2.15 | 36 | 1 | 0 | 1 | 36 | 10 | 31 | 22 |
| 202 | LPC(20:5) [sn2] | $[M+H]^+$ | 542.3 | 184.1 | 1.8 | 24 | 1 | 0 | 5 | 36 | 12 | 41 | 36 |
| 203 | LPC(22:0) [sn2] | $[M+H]^+$ | 580.4 | 184.1 | 4.15 | 30 | 1 | 0 | 1 | 36 | 13 | 32 | 14 |
| 204 | LPC(22:1) [sn2] | $[M+H]^+$ | 578.4 | 184.1 | 3.7 | 24 | 1 | 0 | 5 | 36 | 13 | 32 | 14 |
| 205 | LPC(22:4) [sn2] | $[M+H]^+$ | 572.4 | 184.1 | 2.65 | 24 | 1 | 0 | 1 | 36 | 12 | 41 | 36 |
| 206 | LPC(22:5) (n3) [sn1] [104_sn1] | $[M+H]^+$ | 570.4 | 104.1 | 2.4 | 24 | 1 | 0 | 5 | 36 | 12 | 41 | 36 |
| 207 | LPC(22:5) [sn1] (n3)_LPC(22:5) [sn2] (n6) | $[M+H]^+$ | 570.4 | 184.1 | 2.2 | 48 | 1 | 0 | 1 | 36 | 10 | 25 | 12 |
| 208 | LPC(22:6) [sn2] | $[M+H]^+$ | 568.3 | 184.1 | 2.1 | 36 | 1 | 0 | 1 | 36 | 12 | 41 | 36 |

| | | | | | | | | | | | | | |
|---|---|---|---|---|---|---|---|---|---|---|---|---|---|
| 209 | LPC(24:0) [sn2] | $[M+H]^+$ | 608.5 | 184.1 | 4.85 | 24 | 1 | 0 | 1 | 36 | 12 | 41 | 36 |
| 210 | LPC(26:0) [sn2] | $[M+H]^+$ | 636.5 | 184.1 | 5.85 | 18 | 1 | 0 | 1 | 36 | 12 | 41 | 36 |
| 211 | LPC(O-16:0) | $[M+H]^+$ | 482.4 | 104.1 | 2.9 | 18 | 1 | 0 | 1 | 36 | 13 | 32 | 14 |
| 212 | LPC(O-18:1) | $[M+H]^+$ | 508.4 | 104.103 | 2.9 | 48 | 1 | 0 | 1 | 36 | 4 | 57 | 25 |
| 213 | LPC(O-20:0) | $[M+H]^+$ | 538.4 | 104.103 | 4 | 18 | 1 | 0 | 1 | 36 | 13 | 32 | 14 |
| 214 | LPC(O-20:1) | $[M+H]^+$ | 536.4 | 104.1 | 3.45 | 30 | 1 | 0 | 1 | 36 | 5 | 32 | 10 |
| 215 | LPC(O-22:0) | $[M+H]^+$ | 566.5 | 104.1 | 4.6 | 24 | 1 | 0 | 1 | 36 | 8 | 23 | 9 |
| 216 | LPC(O-22:1) | $[M+H]^+$ | 564.4 | 104.1 | 4 | 24 | 1 | 0 | 1 | 36 | 12 | 41 | 36 |
| 217 | LPC(O-24:0) | $[M+H]^+$ | 594.5 | 104.107 | 5.4 | 24 | 1 | 0 | 5 | 36 | 12 | 41 | 36 |
| 218 | LPC(O-24:1) | $[M+H]^+$ | 592.5 | 104.1 | 4.6 | 24 | 1 | 0 | 5 | 36 | 13 | 32 | 14 |
| 219 | LPC(O-24:2) | $[M+H]^+$ | 590.5 | 104.1 | 4.15 | 18 | 1 | 0 | 5 | 36 | 13 | 32 | 14 |
| 220 | LPC(P-16:0) | $[M+H]^+$ | 480.3 | 104.1 | 2.87 | 18 | 1 | 0 | 1 | 36 | 5 | 32 | 10 |
| 221 | LPC(P-17:0) (a) | $[M+H]^+$ | 494.3 | 104.1 | 3.2 | 18 | 1 | 0 | 1 | 36 | 4 | 57 | 25 |
| 222 | LPC(P-18:1) | $[M+H]^+$ | 506.3 | 104.1 | 2.95 | 18 | 1 | 0 | 1 | 36 | 12 | 41 | 36 |
| 223 | LPE(16:0) [sn2] | $[M+H]^+$ | 454.3 | 313.3 | 2.6 | 24 | 1 | 0 | 5 | 39 | 12 | 41 | 36 |
| 224 | LPE(17:0) [sn2] (no isomer) | $[M+H]^+$ | 468.3 | 327.3 | 2.95 | 36 | 1 | 0 | 5 | 39 | 13 | 32 | 14 |
| 225 | LPE(18:0) [sn2] | $[M+H]^+$ | 482.3 | 341.3 | 3.25 | 24 | 1 | 0 | 1 | 39 | 13 | 32 | 14 |
| 226 | LPE(18:1) [sn2] | $[M+H]^+$ | 480.3 | 339.3 | 2.75 | 24 | 1 | 0 | 5 | 39 | 13 | 32 | 14 |
| 227 | LPE(18:2) [sn2] | $[M+H]^+$ | 478.3 | 337.3 | 2.3 | 24 | 1 | 0 | 5 | 39 | 5 | 32 | 10 |
| 228 | LPE(20:4) [sn2] | $[M+H]^+$ | 502.3 | 361.3 | 2.3 | 24 | 1 | 0 | 5 | 39 | 12 | 41 | 36 |
| 229 | LPE(22:6) [sn2] | $[M+H]^+$ | 526.3 | 385.3 | 2.2 | 24 | 1 | 0 | 5 | 39 | 12 | 41 | 36 |
| 230 | LPE(P-16:0) | $[M+H]^+$ | 438.3 | 266.4 | 2.97 | 18 | 1 | 0 | 5 | 39 | 12 | 41 | 36 |
| 231 | LPE(P-18:0) | $[M+H]^+$ | 466.3 | 294.4 | 3.5 | 24 | 1 | 0 | 5 | 39 | 13 | 32 | 14 |
| 232 | LPE(P-18:1) | $[M+H]^+$ | 464.3 | 292.4 | 3 | 24 | 1 | 0 | 1 | 39 | 8 | 23 | 9 |
| 233 | LPE(P-20:0) | $[M+H]^+$ | 494.3 | 322.4 | 4.05 | 18 | 1 | 0 | 5 | 39 | 7 | 43 | 14 |
| 234 | PC(14:0_16:0) | $[M+H]^+$ | 706.5 | 184.1 | 5.8 | 36 | 1 | 0 | 1 | 47 | 13 | 32 | 14 |
| 235 | PC(14:0_20:4) | $[M+H]^+$ | 754.5 | 184.1 | 5.45 | 24 | 1 | 0 | 1 | 47 | 13 | 32 | 14 |
| 236 | PC(15:0_20:3) | $[M+H]^+$ | 770.6 | 184.1 | 6.15 | 24 | 1 | 0 | 1 | 47 | 5 | 45 | 5 |
| 237 | PC(15:0_20:4) | $[M+H]^+$ | 768.6 | 184.1 | 5.9 | 18 | 1 | 0 | 5 | 47 | 13 | 32 | 14 |
| 238 | PC(15:0_22:6) | $[M+H]^+$ | 792.6 | 184.1 | 5.75 | 18 | 1 | 0 | 1 | 47 | 13 | 32 | 14 |
| 239 | PC(15-MHDA_18:1) | $[M+H]^+$ | 774.6 | 184.1 | 7.45 | 24 | 1 | 0 | 1 | 47 | 13 | 32 | 14 |
| 240 | PC(15-MHDA_18:2) | $[M+H]^+$ | 772.6 | 184.1 | 6.6 | 48 | 1 | 0 | 1 | 47 | 13 | 32 | 14 |
| 241 | PC(15-MHDA_20:4) | $[M+H]^+$ | 796.6 | 184.1 | 6.75 | 30 | 1 | 0 | 5 | 47 | 12 | 41 | 36 |
| 242 | PC(15-MHDA_22:6) | $[M+H]^+$ | 820.6 | 184.1 | 6.55 | 30 | 1 | 0 | 5 | 47 | 12 | 41 | 36 |
| 243 | PC(16:0_16:0) | $[M+H]^+$ | 734.6 | 184.1 | 6.95 | 24 | 1 | 0 | 1 | 47 | 13 | 32 | 14 |
| 244 | PC(16:0_18:0) | $[M+H]^+$ | 762.6 | 184.1 | 8.1 | 24 | 1 | 0 | 1 | 47 | 4 | 56 | 12 |
| 245 | PC(16:0_18:1) | $[M+H]^+$ | 760.6 | 184.1 | 6.9 | 24 | 1 | 0 | 1 | 47 | 13 | 32 | 14 |
| 246 | PC(16:0_18:2) | $[M+H]^+$ | 758.6 | 184.1 | 6.4 | 24 | 1 | 0 | 1 | 47 | 13 | 32 | 14 |
| 247 | PC(16:0_20:4) | $[M+H]^+$ | 782.6 | 184.1 | 5.8 | 48 | 1 | 0 | 1 | 47 | 13 | 32 | 14 |
| 248 | PC(16:0_22:6) | $[M+H]^+$ | 806.6 | 184.1 | 6.15 | 24 | 1 | 0 | 5 | 47 | 13 | 32 | 14 |
| 249 | PC(16:1_18:2) | $[M+H]^+$ | 756.6 | 184.1 | 5.7 | 42 | 1 | 0 | 5 | 47 | 7 | 43 | 14 |
| 250 | PC(16:1_20:4) | $[M+H]^+$ | 780.6 | 184.1 | 5.4 | 48 | 1 | 0 | 1 | 47 | 12 | 41 | 36 |
| 251 | PC(18:0_18:1) | $[M+H]^+$ | 788.6 | 184.1 | 8.3 | 24 | 1 | 0 | 5 | 47 | 12 | 41 | 36 |
| 252 | PC(18:0_20:3) | $[M+H]^+$ | 812.6 | 184.1 | 7.9 | 24 | 1 | 0 | 1 | 47 | 13 | 32 | 14 |
| 253 | PC(18:0_22:4) | $[M+H]^+$ | 838.6 | 184.1 | 8.35 | 48 | 1 | 0 | 1 | 47 | 13 | 32 | 14 |
| 254 | PC(18:0_22:5) (n3)_PC(20:1_20:4) | $[M+H]^+$ | 836.6 | 184.1 | 7.55 | 42 | 1 | 0 | 5 | 47 | 12 | 41 | 36 |
| 255 | PC(18:0_22:6) | $[M+H]^+$ | 834.6 | 184.1 | 7.2 | 24 | 1 | 0 | 1 | 47 | 12 | 41 | 36 |
| 256 | PC(18:1_18:1) | $[M+H]^+$ | 786.6 | 184.1 | 7.35 | 36 | 1 | 0 | 1 | 47 | 12 | 41 | 36 |
| 257 | PC(18:1_18:2) | $[M+H]^+$ | 784.6 | 184.1 | 6.6 | 36 | 1 | 0 | 1 | 47 | 5 | 45 | 5 |
| 258 | PC(18:1_20:3) | $[M+H]^+$ | 810.6 | 184.1 | 6.7 | 72 | 1 | 0 | 5 | 47 | 13 | 32 | 14 |
| 259 | PC(18:2_20:5) | $[M+H]^+$ | 804.6 | 184.1 | 5.2 | 60 | 1 | 0 | 5 | 47 | 13 | 32 | 14 |
| 260 | PC(28:0) | $[M+H]^+$ | 678.5 | 184.1 | 5.1 | 24 | 1 | 0 | 5 | 47 | 12 | 41 | 36 |
| 261 | PC(31:0) (a) | $[M+H]^+$ | 720.6 | 184.1 | 6.3 | 30 | 1 | 0 | 1 | 47 | 8 | 23 | 9 |
| 262 | PC(31:1) | $[M+H]^+$ | 718.5 | 184.1 | 5.7 | 24 | 1 | 0 | 5 | 47 | 7 | 43 | 14 |
| 263 | PC(32:1) | $[M+H]^+$ | 732.6 | 184.1 | 6.12 | 27.6 | 1 | 0 | 1 | 47 | 12 | 41 | 36 |
| 264 | PC(32:2) | $[M+H]^+$ | 730.5 | 184.1 | 5.45 | 24 | 1 | 0 | 1 | 47 | 13 | 32 | 14 |
| 265 | PC(33:0) (a) | $[M+H]^+$ | 748.6 | 184.1 | 7.35 | 24 | 1 | 0 | 1 | 47 | 8 | 23 | 9 |
| 266 | PC(33:1) | $[M+H]^+$ | 746.6 | 184.1 | 7.1 | 24 | 1 | 0 | 5 | 47 | 4 | 56 | 12 |
| 267 | PC(33:2) | $[M+H]^+$ | 744.6 | 184.1 | 5.9 | 24 | 1 | 0 | 1 | 47 | 12 | 41 | 36 |
| 268 | PC(34:5) | $[M+H]^+$ | 752.5 | 184.1 | 4.95 | 24 | 1 | 0 | 5 | 47 | 5 | 32 | 10 |
| 269 | PC(35:5) | $[M+H]^+$ | 766.5 | 184.1 | 5.4 | 18 | 1 | 0 | 1 | 47 | 13 | 32 | 14 |
| 270 | PC(36:6) (a) | $[M+H]^+$ | 778.5 | 184.1 | 5.2 | 30 | 1 | 0 | 5 | 47 | 12 | 41 | 36 |
| 271 | PC(38:2) | $[M+H]^+$ | 814.6 | 184.1 | 8.5 | 36 | 1 | 0 | 1 | 47 | 12 | 41 | 36 |
| 272 | PC(38:5) (a) | $[M+H]^+$ | 808.6 | 184.1 | 6.5 | 36 | 1 | 0 | 1 | 47 | 13 | 32 | 14 |
| 273 | PC(39:5) (a) | $[M+H]^+$ | 822.6 | 184.1 | 6.75 | 36 | 1 | 0 | 1 | 47 | 12 | 41 | 36 |
| 274 | PC(40:10) | $[M+H]^+$ | 826.6 | 184.1 | 4.8 | 24 | 1 | 0 | 1 | 47 | 13 | 32 | 14 |
| 275 | PC(40:7) (a) | $[M+H]^+$ | 832.6 | 184.1 | 6.1 | 48 | 1 | 0 | 5 | 47 | 5 | 32 | 10 |
| 276 | PC(40:8) | $[M+H]^+$ | 830.6 | 184.1 | 5.7 | 24 | 1 | 0 | 5 | 47 | 12 | 41 | 36 |
| 277 | PC(44:12) | $[M+H]^+$ | 878.6 | 184.1 | 5.4 | 24 | 1 | 0 | 5 | 47 | 12 | 41 | 36 |
| 278 | PC(O-16:0_16:0) | $[M+H]^+$ | 720.6 | 184.103 | 7.6 | 36 | 1 | 0 | 5 | 47 | 8 | 23 | 9 |
| 279 | PC(O-16:0_20:4) | $[M+H]^+$ | 768.6 | 184.103 | 6.8 | 36 | 1 | 0 | 5 | 47 | 4 | 56 | 12 |
| 280 | PC(O-16:0_22:6) | $[M+H]^+$ | 792.6 | 184.103 | 6.9 | 48 | 1 | 0 | 1 | 47 | 12 | 41 | 36 |
| 281 | PC(O-18:0_18:1) | $[M+H]^+$ | 774.6 | 184.103 | 9.15 | 24 | 1 | 0 | 1 | 47 | 13 | 32 | 14 |

| | | | | | | | | | | | | | |
|---|---|---|---|---|---|---|---|---|---|---|---|---|---|
| 282 | PC(O-18:0_20:4) | $[M+H]^+$ | 796.6 | 184.103 | 7.75 | 42 | 1 | 0 | 5 | 47 | 13 | 32 | 14 |
| 283 | PC(O-18:0_22:6) | $[M+H]^+$ | 820.6 | 184.103 | 7.95 | 30 | 1 | 0 | 5 | 47 | 4 | 57 | 25 |
| 284 | PC(O-18:1_18:1) | $[M+H]^+$ | 772.6 | 184.103 | 8.1 | 36 | 1 | 0 | 1 | 47 | 13 | 32 | 14 |
| 285 | PC(O-18:1_18:2) | $[M+H]^+$ | 770.6 | 184.103 | 7.1 | 48 | 1 | 0 | 1 | 47 | 13 | 32 | 14 |
| 286 | PC(O-32:1) | $[M+H]^+$ | 718.5 | 184.103 | 7.2 | 48 | 1 | 0 | 5 | 47 | 8 | 23 | 9 |
| 287 | PC(O-32:2) | $[M+H]^+$ | 716.6 | 184.1 | 6 | 72 | 1 | 0 | 5 | 47 | 5 | 26 | 25 |
| 288 | PC(O-34:1) | $[M+H]^+$ | 746.6 | 184.103 | 7.8 | 36 | 1 | 0 | 5 | 47 | 13 | 32 | 14 |
| 289 | PC(O-34:2) | $[M+H]^+$ | 744.6 | 184.103 | 7 | 36 | 1 | 0 | 5 | 47 | 5 | 25 | 20 |
| 290 | PC(O-34:4) | $[M+H]^+$ | 740.6 | 184.1 | 6.05 | 36 | 1 | 0 | 5 | 47 | 4 | 56 | 12 |
| 291 | PC(O-35:4) | $[M+H]^+$ | 754.5 | 184.103 | 6.56 | 24 | 1 | 0 | 1 | 47 | 12 | 41 | 36 |
| 292 | PC(O-36:0) | $[M+H]^+$ | 776.6 | 184.1 | 10.05 | 24 | 1 | 0 | 1 | 47 | 13 | 32 | 14 |
| 293 | PC(O-36:5) | $[M+H]^+$ | 766.5 | 184.103 | 6.35 | 60 | 1 | 0 | 5 | 47 | 13 | 32 | 14 |
| 294 | PC(O-38:5) | $[M+H]^+$ | 794.6 | 184.1 | 7.15 | 24 | 1 | 0 | 1 | 47 | 13 | 32 | 14 |
| 295 | PC(O-40:5) | $[M+H]^+$ | 822.6 | 184.103 | 8.05 | 42 | 1 | 0 | 1 | 47 | 13 | 32 | 14 |
| 296 | PC(O-40:7) (a) | $[M+H]^+$ | 818.6 | 184.1 | 7 | 24 | 1 | 0 | 5 | 47 | 13 | 32 | 14 |
| 297 | PC(P-15:0_20:4) (a) | $[M+H]^+$ | 752.5 | 184.103 | 5.85 | 54 | 1 | 0 | 5 | 47 | 12 | 41 | 36 |
| 298 | PC(P-16:0_14:0) | $[M+H]^+$ | 690.4 | 184.1 | 6.5 | 18 | 1 | 0 | 1 | 47 | 13 | 32 | 14 |
| 299 | PC(P-16:0_18:0) | $[M+H]^+$ | 746.6 | 184.107 | 8.55 | 18 | 1 | 0 | 5 | 47 | 5 | 45 | 5 |
| 300 | PC(P-16:0_18:1) | $[M+H]^+$ | 744.6 | 184.107 | 7.6 | 36 | 1 | 0 | 5 | 47 | 5 | 25 | 20 |
| 301 | PC(P-16:0_18:2) | $[M+H]^+$ | 742.5 | 184.1 | 6.95 | 24 | 1 | 0 | 1 | 47 | 8 | 23 | 9 |
| 302 | PC(P-16:0_20:5) | $[M+H]^+$ | 764.6 | 184.102 | 6.2 | 24 | 1 | 0 | 1 | 47 | 8 | 23 | 9 |
| 303 | PC(P-16:0_22:6) | $[M+H]^+$ | 790.6 | 184.1 | 6.6 | 30 | 1 | 0 | 5 | 47 | 13 | 32 | 14 |
| 304 | PC(P-17:0_20:4) (a) | $[M+H]^+$ | 780.6 | 184.103 | 7.2 | 42 | 1 | 0 | 5 | 47 | 7 | 43 | 14 |
| 305 | PC(P-18:0_20:4) | $[M+H]^+$ | 794.6 | 184.103 | 8 | 24 | 1 | 0 | 5 | 47 | 13 | 32 | 14 |
| 306 | PC(P-18:0_22:6) | $[M+H]^+$ | 818.6 | 184.103 | 7.6 | 30 | 1 | 0 | 5 | 47 | 12 | 41 | 36 |
| 307 | PC(P-18:1_18:1) | $[M+H]^+$ | 770.6 | 184.107 | 7.95 | 36 | 1 | 0 | 1 | 47 | 13 | 32 | 14 |
| 308 | PC(P-18:1_22:6) | $[M+H]^+$ | 816.6 | 184.1 | 6.6 | 36 | 1 | 0 | 5 | 47 | 7 | 43 | 14 |
| 309 | PC(P-20:0_20:4) | $[M+H]^+$ | 822.6 | 184.107 | 9.3 | 24 | 1 | 0 | 1 | 47 | 12 | 41 | 36 |
| 310 | PC(P-35:2) (a) | $[M+H]^+$ | 756.6 | 184.103 | 7 | 60 | 1 | 0 | 5 | 47 | 4 | 56 | 12 |
| 311 | PE(15-MHDA_18:1) | $[M+H]^+$ | 732.6 | 591.5 | 7.95 | 36 | 1 | 0 | 5 | 52 | 4 | 56 | 12 |
| 312 | PE(15-MHDA_18:2) | $[M+H]^+$ | 730.5 | 589.5 | 6.8 | 60 | 1 | 0 | 5 | 52 | 5 | 32 | 10 |
| 313 | PE(15-MHDA_20:4) | $[M+H]^+$ | 754.6 | 613.5 | 7.1 | 30 | 1 | 0 | 5 | 52 | 5 | 45 | 5 |
| 314 | PE(15-MHDA_22:6) | $[M+H]^+$ | 778.5 | 637.6 | 6.85 | 30 | 1 | 0 | 5 | 52 | 5 | 25 | 20 |
| 315 | PE(16:0_16:0) | $[M+H]^+$ | 692.5 | 551.5 | 7.25 | 24 | 1 | 0 | 5 | 52 | 5 | 25 | 20 |
| 316 | PE(16:0_16:1) | $[M+H]^+$ | 690.5 | 549.5 | 6.2 | 36 | 1 | 0 | 1 | 52 | 13 | 32 | 14 |
| 317 | PE(16:0_18:1) | $[M+H]^+$ | 718.5 | 577.5 | 7.25 | 36 | 1 | 0 | 1 | 52 | 5 | 32 | 10 |
| 318 | PE(16:0_18:2) | $[M+H]^+$ | 716.5 | 575.5 | 6.65 | 30 | 1 | 0 | 5 | 52 | 13 | 32 | 14 |
| 319 | PE(16:0_20:4) | $[M+H]^+$ | 740.5 | 599.5 | 6.4 | 36 | 1 | 0 | 5 | 52 | 13 | 32 | 14 |
| 320 | PE(16:0_22:6) | $[M+H]^+$ | 764.5 | 623.5 | 6 | 24 | 1 | 0 | 1 | 52 | 5 | 25 | 20 |
| 321 | PE(16:1_18:2) | $[M+H]^+$ | 714.5 | 573.502 | 5.95 | 36 | 1 | 0 | 1 | 52 | 5 | 25 | 20 |
| 322 | PE(16:1_20:4) | $[M+H]^+$ | 738.5 | 597.5 | 5.85 | 36 | 1 | 0 | 5 | 52 | 5 | 25 | 20 |
| 323 | PE(18:0_18:1) | $[M+H]^+$ | 746.6 | 605.6 | 8.1 | 30 | 1 | 0 | 5 | 52 | 5 | 25 | 20 |
| 324 | PE(18:0_20:3) (a) | $[M+H]^+$ | 770.6 | 629.6 | 8.2 | 48 | 1 | 0 | 5 | 52 | 10 | 15 | 13 |
| 325 | PE(18:0_20:4) | $[M+H]^+$ | 768.6 | 627.5 | 7.45 | 42 | 1 | 0 | 5 | 52 | 7 | 43 | 14 |
| 326 | PE(18:0_22:4) | $[M+H]^+$ | 796.6 | 655.6 | 8.5 | 48 | 1 | 0 | 5 | 52 | 5 | 25 | 20 |
| 327 | PE(18:0_22:5) (n3) | $[M+H]^+$ | 794.6 | 653.6 | 7.6 | 24 | 1 | 0 | 5 | 52 | 4 | 56 | 12 |
| 328 | PE(18:0_22:6) | $[M+H]^+$ | 792.6 | 651.5 | 7.3 | 30 | 1 | 0 | 5 | 52 | 8 | 23 | 9 |
| 329 | PE(18:1_18:1) | $[M+H]^+$ | 744.6 | 603.5 | 7.55 | 48 | 1 | 0 | 5 | 52 | 8 | 23 | 9 |
| 330 | PE(18:1_18:2) | $[M+H]^+$ | 742.5 | 601.5 | 6.6 | 48 | 1 | 0 | 5 | 52 | 5 | 25 | 20 |
| 331 | PE(18:1_22:6) (a) | $[M+H]^+$ | 790.5 | 649.5 | 6.6 | 36 | 1 | 0 | 5 | 52 | 8 | 23 | 9 |
| 332 | PE(36:0) | $[M+H]^+$ | 748.6 | 607.6 | 9.7 | 24 | 1 | 0 | 5 | 52 | 12 | 41 | 36 |
| 333 | PE(38:5) (a) | $[M+H]^+$ | 766.5 | 625.5 | 6.8 | 48 | 1 | 0 | 5 | 52 | 5 | 32 | 10 |
| 334 | PE(O-16:0_18:2) | $[M+H]^+$ | 702.5 | 561.5 | 6.8 | 24 | 1 | 0 | 5 | 52 | 5 | 25 | 20 |
| 335 | PE(O-16:0_20:3) | $[M+H]^+$ | 728.6 | 587.5 | 7.65 | 36 | 1 | 0 | 5 | 52 | 5 | 25 | 20 |
| 336 | PE(O-16:0_20:4) | $[M+H]^+$ | 726.5 | 585.5 | 6.78 | 24 | 1 | 0 | 5 | 52 | 13 | 32 | 14 |
| 337 | PE(O-16:0_22:4) | $[M+H]^+$ | 754.6 | 613.503 | 8.3 | 48 | 1 | 0 | 5 | 52 | 5 | 32 | 10 |
| 338 | PE(O-16:0_22:6) | $[M+H]^+$ | 750.6 | 609.5 | 6.9 | 36 | 1 | 0 | 5 | 52 | 5 | 25 | 20 |
| 339 | PE(O-18:0_22:5) (a) | $[M+H]^+$ | 780.6 | 639.6 | 8.7 | 24 | 1 | 0 | 5 | 52 | 4 | 56 | 12 |
| 340 | PE(O-18:0_22:6) | $[M+H]^+$ | 778.5 | 637.603 | 7.9 | 24 | 1 | 0 | 5 | 52 | 13 | 32 | 14 |
| 341 | PE(O-18:1_22:6) | $[M+H]^+$ | 776.6 | 635.5 | 7.15 | 36 | 1 | 0 | 5 | 52 | 13 | 32 | 14 |
| 342 | PE(O-34:1) | $[M+H]^+$ | 704.6 | 563.5 | 8.15 | 30 | 1 | 0 | 5 | 52 | 12 | 41 | 36 |
| 343 | PE(O-36:5) | $[M+H]^+$ | 724.5 | 583.5 | 6.25 | 24 | 1 | 0 | 5 | 52 | 5 | 25 | 20 |
| 344 | PE(O-38:5) (a) | $[M+H]^+$ | 752.6 | 611.5 | 7.5 | 48 | 1 | 0 | 5 | 52 | 8 | 23 | 9 |
| 345 | PE(P-15:0_20:4) (a) | $[M+H]^+$ | 710.5 | 361.3 | 6.5 | 24 | 1 | 0 | 1 | 52 | 13 | 32 | 14 |
| 346 | PE(P-15:0_22:6) (a) | $[M+H]^+$ | 734.5 | 385.3 | 6.4 | 24 | 1 | 0 | 5 | 52 | 8 | 23 | 9 |
| 347 | PE(P-16:0_18:1) | $[M+H]^+$ | 702.5 | 339.3 | 8.1 | 24 | 1 | 0 | 1 | 52 | 8 | 23 | 9 |
| 348 | PE(P-16:0_18:2) | $[M+H]^+$ | 700.5 | 337.3 | 7.3 | 24 | 1 | 0 | 5 | 52 | 13 | 32 | 14 |
| 349 | PE(P-16:0_18:3) | $[M+H]^+$ | 698.5 | 335.3 | 6.4 | 36 | 1 | 0 | 5 | 52 | 8 | 23 | 9 |
| 350 | PE(P-16:0_20:3) (a) | $[M+H]^+$ | 726.5 | 363.3 | 7.55 | 30 | 1 | 0 | 5 | 52 | 5 | 25 | 20 |
| 351 | PE(P-16:0_20:4) | $[M+H]^+$ | 724.5 | 361.3 | 7.2 | 24 | 1 | 0 | 5 | 52 | 5 | 25 | 20 |
| 352 | PE(P-16:0_20:5) | $[M+H]^+$ | 722.5 | 359.3 | 6.15 | 24 | 1 | 0 | 1 | 52 | 5 | 25 | 20 |
| 353 | PE(P-16:0_22:4) | $[M+H]^+$ | 752.6 | 389.3 | 8.1 | 24 | 1 | 0 | 1 | 52 | 5 | 32 | 10 |
| 354 | PE(P-16:0_22:5) (n3) | $[M+H]^+$ | 750.5 | 387.3 | 7 | 60 | 1 | 0 | 5 | 52 | 5 | 26 | 25 |

| | | | | | | | | | | | | | |
|---|---|---|---|---|---|---|---|---|---|---|---|---|---|
| 355 | PE(P-16:0_22:6) | $[M+H]^+$ | 748.5 | 385.3 | 6.8 | 30 | 1 | 0 | 5 | 52 | 5 | 25 | 20 |
| 356 | PE(P-17:0_20:4) (a) | $[M+H]^+$ | 738.6 | 361.3 | 7.3 | 48 | 1 | 0 | 5 | 52 | 5 | 32 | 10 |
| 357 | PE(P-17:0_22:6) (a) | $[M+H]^+$ | 762.6 | 385.3 | 7.15 | 42 | 1 | 0 | 5 | 52 | 5 | 32 | 10 |
| 358 | PE(P-18:0_18:1) | $[M+H]^+$ | 730.6 | 339.3 | 9.4 | 24 | 1 | 0 | 1 | 52 | 13 | 32 | 14 |
| 359 | PE(P-18:0_18:2) | $[M+H]^+$ | 728.6 | 337.3 | 8.3 | 30 | 1 | 0 | 5 | 52 | 5 | 32 | 10 |
| 360 | PE(P-18:0_18:3) | $[M+H]^+$ | 726.5 | 335.3 | 7.5 | 36 | 1 | 0 | 1 | 52 | 8 | 23 | 9 |
| 361 | PE(P-18:0_20:3) (a) | $[M+H]^+$ | 754.5 | 363.3 | 7.9 | 60 | 1 | 0 | 5 | 52 | 4 | 56 | 12 |
| 362 | PE(P-18:0_20:4) | $[M+H]^+$ | 752.6 | 361.3 | 8.35 | 24 | 1 | 0 | 5 | 52 | 13 | 32 | 14 |
| 363 | PE(P-18:0_20:5) | $[M+H]^+$ | 750.5 | 359.3 | 7.4 | 30 | 1 | 0 | 5 | 52 | 5 | 25 | 20 |
| 364 | PE(P-18:0_22:4) | $[M+H]^+$ | 780.6 | 389.3 | 8.8 | 24 | 1 | 0 | 5 | 52 | 7 | 43 | 14 |
| 365 | PE(P-18:0_22:5) (n3) | $[M+H]^+$ | 778.5 | 387.3 | 8.45 | 42 | 1 | 0 | 5 | 52 | 7 | 43 | 14 |
| 366 | PE(P-18:0_22:6) | $[M+H]^+$ | 776.6 | 385.3 | 7.85 | 42 | 1 | 0 | 5 | 52 | 8 | 23 | 9 |
| 367 | PE(P-18:1_18:1) (a) | $[M+H]^+$ | 728.6 | 339.3 | 8.15 | 42 | 1 | 0 | 5 | 52 | 5 | 25 | 20 |
| 368 | PE(P-18:1_18:2) (a) | $[M+H]^+$ | 726.5 | 337.3 | 7.3 | 30 | 1 | 0 | 5 | 52 | 7 | 43 | 14 |
| 369 | PE(P-18:1_18:3) | $[M+H]^+$ | 724.5 | 335.3 | 6.85 | 18 | 1 | 0 | 1 | 52 | 5 | 25 | 20 |
| 370 | PE(P-18:1_20:3) (a) | $[M+H]^+$ | 752.5 | 363.3 | 7.65 | 30 | 1 | 0 | 5 | 52 | 12 | 41 | 36 |
| 371 | PE(P-18:1_20:4) (a) | $[M+H]^+$ | 750.5 | 361.3 | 7.15 | 36 | 1 | 0 | 5 | 52 | 5 | 25 | 20 |
| 372 | PE(P-18:1_20:5) (a) | $[M+H]^+$ | 748.5 | 359.3 | 6.4 | 42 | 1 | 0 | 5 | 52 | 7 | 43 | 14 |
| 373 | PE(P-18:1_22:4) | $[M+H]^+$ | 778.5 | 389.3 | 8 | 42 | 1 | 0 | 5 | 52 | 4 | 56 | 12 |
| 374 | PE(P-18:1_22:5) (a) | $[M+H]^+$ | 776.6 | 387.3 | 7.5 | 48 | 1 | 0 | 5 | 52 | 8 | 23 | 9 |
| 375 | PE(P-18:1_22:6) (a) | $[M+H]^+$ | 774.5 | 385.3 | 6.9 | 42 | 1 | 0 | 5 | 52 | 8 | 23 | 9 |
| 376 | PE(P-19:0_20:4) (a) | $[M+H]^+$ | 766.6 | 361.3 | 8.7 | 48 | 1 | 0 | 5 | 52 | 5 | 25 | 20 |
| 377 | PE(P-20:0_18:1) | $[M+H]^+$ | 758.6 | 339.3 | 10.25 | 24 | 1 | 0 | 5 | 52 | 5 | 32 | 10 |
| 378 | PE(P-20:0_18:2) | $[M+H]^+$ | 756.6 | 337.3 | 9.8 | 24 | 1 | 0 | 5 | 52 | 5 | 32 | 10 |
| 379 | PE(P-20:0_20:4) | $[M+H]^+$ | 780.6 | 361.3 | 9.45 | 36 | 1 | 0 | 5 | 52 | 5 | 32 | 10 |
| 380 | PE(P-20:0_22:6) | $[M+H]^+$ | 804.6 | 385.3 | 9.2 | 30 | 1 | 0 | 5 | 52 | 8 | 23 | 9 |
| 381 | PE(P-20:1_20:4) | $[M+H]^+$ | 778.5 | 361.3 | 8.3 | 36 | 1 | 0 | 5 | 52 | 8 | 23 | 9 |
| 382 | PE(P-20:1_22:6) (a) | $[M+H]^+$ | 802.6 | 385.3 | 8.3 | 30 | 1 | 0 | 5 | 52 | 8 | 23 | 9 |
| 383 | PG(34:1) | $[M+NH_4]^+$ | 766.6 | 577.5 | 6.01 | 18 | 1 | 0 | 5 | 60 | 8 | 23 | 9 |
| 384 | PG(34:2) | $[M+NH_4]^+$ | 764.6 | 575.5 | 5.5 | 36 | 1 | 0 | 5 | 60 | 5 | 25 | 20 |
| 385 | PG(36:1) | $[M+NH_4]^+$ | 794.6 | 605.6 | 6.96 | 24 | 1 | 0 | 5 | 60 | 13 | 32 | 14 |
| 386 | PG(36:2) | $[M+NH_4]^+$ | 792.6 | 603.5 | 6.2 | 30 | 1 | 0 | 5 | 60 | 5 | 32 | 10 |
| 387 | PI(16:0_16:0) | $[M+NH_4]^+$ | 828.6 | 551.6 | 5.55 | 24 | 1 | 0 | 5 | 52 | 5 | 25 | 20 |
| 388 | PI(16:0_16:1) | $[M+NH_4]^+$ | 826.5 | 549.5 | 5.05 | 18 | 1 | 0 | 5 | 52 | 8 | 23 | 9 |
| 389 | PI(16:0_20:3) (a) | $[M+NH_4]^+$ | 878.6 | 601.6 | 5.05 | 42 | 1 | 0 | 5 | 52 | 5 | 26 | 25 |
| 390 | PI(16:0_20:4) | $[M+NH_4]^+$ | 876.6 | 599.6 | 5 | 24 | 1 | 0 | 5 | 52 | 5 | 25 | 20 |
| 391 | PI(17:0_18:1) (a+b) | $[M+NH_4]^+$ | 868.6 | 591.6 | 6.15 | 18 | 1 | 0 | 5 | 52 | 5 | 32 | 10 |
| 392 | PI(17:0_18:2) (a+b) | $[M+NH_4]^+$ | 866.6 | 589.6 | 5.4 | 24 | 1 | 0 | 5 | 52 | 8 | 23 | 9 |
| 393 | PI(17:0_20:4) (a+b) | $[M+NH_4]^+$ | 890.6 | 613.6 | 5.5 | 24 | 1 | 0 | 5 | 52 | 8 | 23 | 9 |
| 394 | PI(18:0_18:1) | $[M+NH_4]^+$ | 882.6 | 605.6 | 6.35 | 36 | 1 | 0 | 5 | 52 | 8 | 23 | 9 |
| 395 | PI(18:0_20:2) | $[M+NH_4]^+$ | 908.6 | 631.6 | 6.82 | 24 | 1 | 0 | 5 | 52 | 8 | 23 | 9 |
| 396 | PI(18:0_20:3) (a) | $[M+NH_4]^+$ | 906.6 | 629.6 | 6.1 | 36 | 1 | 0 | 1 | 52 | 7 | 43 | 14 |
| 397 | PI(18:0_20:4) | $[M+NH_4]^+$ | 904.6 | 627.6 | 5.65 | 36 | 1 | 0 | 5 | 52 | 8 | 23 | 9 |
| 398 | PI(18:0_22:4) | $[M+NH_4]^+$ | 932.6 | 655.6 | 6.5 | 36 | 1 | 0 | 5 | 52 | 8 | 23 | 9 |
| 399 | PI(18:0_22:5) (n3) | $[M+NH_4]^+$ | 930.6 | 653.6 | 6 | 42 | 1 | 0 | 5 | 52 | 8 | 23 | 9 |
| 400 | PI(18:0_22:6) | $[M+NH_4]^+$ | 928.6 | 651.6 | 5.8 | 24 | 1 | 0 | 5 | 52 | 8 | 23 | 9 |
| 401 | PI(34:0) | $[M+NH_4]^+$ | 856.6 | 579.6 | 6.5 | 18 | 1 | 0 | 5 | 52 | 8 | 23 | 9 |
| 402 | PI(34:1) | $[M+NH_4]^+$ | 854.6 | 577.6 | 5.6 | 24 | 1 | 0 | 5 | 52 | 8 | 23 | 9 |
| 403 | PI(36:2) (a+b) | $[M+NH_4]^+$ | 880.6 | 603.6 | 5.7 | 36 | 1 | 0 | 5 | 52 | 8 | 23 | 9 |
| 404 | PI(37:6) | $[M+NH_4]^+$ | 886.6 | 609.6 | 5.45 | 18 | 1 | 0 | 5 | 52 | 8 | 23 | 9 |
| 405 | PI(38:5) (a) | $[M+NH_4]^+$ | 902.6 | 625.602 | 4.95 | 48 | 1 | 0 | 5 | 52 | 8 | 23 | 9 |
| 406 | PI(38:6) | $[M+NH_4]^+$ | 900.6 | 623.6 | 4.85 | 30 | 1 | 0 | 5 | 52 | 8 | 23 | 9 |
| 407 | PI(39:6) (a+b) | $[M+NH_4]^+$ | 914.6 | 637.6 | 5.35 | 18 | 1 | 0 | 5 | 52 | 8 | 23 | 9 |
| 408 | PS(36:1) | $[M+H]^+$ | 790.6 | 605.6 | 6.75 | 60 | 1 | 0 | 5 | 86 | 5 | 29 | 5 |
| 409 | PS(36:2) | $[M+H]^+$ | 788.5 | 603.5 | 6.2 | 36 | 1 | 0 | 5 | 86 | 8 | 23 | 9 |
| 410 | PS(38:3) | $[M+H]^+$ | 814.6 | 629.6 | 6.3 | 36 | 1 | 0 | 5 | 86 | 8 | 23 | 9 |
| 411 | PS(38:4) | $[M+H]^+$ | 812.5 | 627.5 | 6.05 | 48 | 1 | 0 | 5 | 86 | 8 | 23 | 9 |
| 412 | PS(38:5) | $[M+H]^+$ | 810.5 | 625.5 | 5.25 | 60 | 1 | 0 | 5 | 86 | 8 | 23 | 9 |
| 413 | PS(40:5) | $[M+H]^+$ | 838.6 | 653.6 | 6.35 | 24 | 1 | 0 | 5 | 86 | 8 | 23 | 9 |
| 414 | PS(40:6) | $[M+H]^+$ | 836.5 | 651.5 | 5.9 | 48 | 1 | 0 | 5 | 86 | 5 | 26 | 25 |
| 415 | S1P(d16:1) | $[M+H]^+$ | 352.2 | 236.3 | 1.4 | 24 | 1 | 0 | 5 | 25 | 5 | 26 | 25 |
| 416 | S1P(d18:0) | $[M+H]^+$ | 382.2 | 284.3 | 2.2 | 24 | 1 | 0 | 5 | 25 | 8 | 23 | 9 |
| 417 | S1P(d18:1) | $[M+H]^+$ | 380.2 | 264.3 | 1.8 | 24 | 1 | 0 | 5 | 25 | 8 | 23 | 9 |
| 418 | S1P(d18:2) | $[M+H]^+$ | 378.2 | 262.3 | 1.6 | 24 | 1 | 0 | 5 | 25 | 8 | 23 | 9 |
| 419 | SM(34:3) | $[M+H]^+$ | 699.5 | 184.1 | 4.65 | 30 | 1 | 0 | 5 | 215 | 8 | 23 | 9 |
| 420 | SM(35:2) (b) | $[M+H]^+$ | 715.6 | 184.1 | 5.55 | 30 | 1 | 0 | 1 | 215 | 8 | 23 | 9 |
| 421 | SM(37:1) | $[M+H]^+$ | 745.6 | 184.1 | 7.1 | 60 | 1 | 0 | 5 | 215 | 8 | 23 | 9 |
| 422 | SM(37:2) | $[M+H]^+$ | 743.5 | 184.1 | 6.25 | 48 | 1 | 0 | 5 | 215 | 8 | 23 | 9 |
| 423 | SM(38:3) (a) | $[M+H]^+$ | 755.6 | 184.1 | 6.15 | 48 | 1 | 0 | 1 | 215 | 8 | 23 | 9 |
| 424 | SM(40:3) (a) | $[M+H]^+$ | 783.6 | 184.1 | 7.15 | 42 | 1 | 0 | 5 | 215 | 8 | 23 | 9 |
| 425 | SM(41:0) | $[M+H]^+$ | 803.7 | 184.1 | 10.15 | 24 | 1 | 0 | 1 | 215 | 5 | 26 | 25 |
| 426 | SM(41:1) (a) | $[M+H]^+$ | 801.7 | 184.1 | 9.5 | 48 | 1 | 0 | 1 | 215 | 8 | 23 | 9 |
| 427 | SM(43:1) | $[M+H]^+$ | 829.7 | 184.1 | 10.33 | 30 | 1 | 0 | 1 | 215 | 8 | 23 | 9 |

| | | | | | | | | | | | | | |
|---|---|---|---|---|---|---|---|---|---|---|---|---|---|
| 428 | SM(43:2) (b) | $[M+H]^+$ | 827.7 | 184.1 | 9.7 | 48 | 1 | 0 | 1 | 215 | 8 | 23 | 9 |
| 429 | SM(44:1) | $[M+H]^+$ | 843.6 | 184.1 | 10.25 | 48 | 1 | 0 | 5 | 215 | 8 | 23 | 9 |
| 430 | SM(44:2) | $[M+H]^+$ | 841.6 | 184.1 | 10.16 | 24 | 1 | 0 | 5 | 215 | 5 | 25 | 20 |
| 431 | SM(44:3) | $[M+H]^+$ | 839.6 | 184.1 | 9.6 | 24 | 1 | 0 | 1 | 215 | 5 | 25 | 20 |
| 432 | SM(d16:1_19:0) | $[M+H]^+$ | 717.6 | 184.1 | 6.15 | 36 | 1 | 0 | 1 | 215 | 8 | 11 | 21 |
| 433 | SM(d16:1_23:0)_SM(d17:1_22:0) | $[M+H]^+$ | 773.7 | 184.1 | 8.6 | 30 | 1 | 0 | 1 | 215 | 8 | 11 | 21 |
| 434 | SM(d16:1_24:1) | $[M+H]^+$ | 785.7 | 184.1 | 8.03 | 42 | 1 | 0 | 5 | 215 | 8 | 43 | 16 |
| 435 | SM(d17:1_14:0) | $[M+H]^+$ | 661.5 | 184.1 | 4.65 | 24 | 1 | 0 | 5 | 215 | 8 | 11 | 21 |
| 436 | SM(d17:1_16:0) | $[M+H]^+$ | 689.6 | 184.1 | 5.35 | 24 | 1 | 0 | 5 | 215 | 8 | 11 | 21 |
| 437 | SM(d17:1_24:1) | $[M+H]^+$ | 799.7 | 184.1 | 8.65 | 42 | 1 | 0 | 1 | 215 | 10 | 21 | 12 |
| 438 | SM(d18:0_14:0) | $[M+H]^+$ | 677.6 | 184.1 | 5.28 | 24 | 1 | 0 | 1 | 215 | 8 | 11 | 21 |
| 439 | SM(d18:0_16:0) | $[M+H]^+$ | 705.6 | 184.1 | 6.15 | 24 | 1 | 0 | 1 | 215 | 8 | 11 | 21 |
| 440 | SM(d18:0_22:0) | $[M+H]^+$ | 789.7 | 184.1 | 9.8 | 30 | 1 | 0 | 1 | 215 | 8 | 11 | 21 |
| 441 | SM(d18:1_14:0)_SM(d16:1_16:0) | $[M+H]^+$ | 675.5 | 184.1 | 4.95 | 30 | 1 | 0 | 1 | 215 | 8 | 11 | 21 |
| 442 | SM(d18:1_16:0) | $[M+H]^+$ | 703.6 | 184.1 | 5.75 | 30 | 1 | 0 | 1 | 215 | 8 | 11 | 21 |
| 443 | SM(d18:1_18:0)_SM(d16:1_20:0) | $[M+H]^+$ | 731.6 | 184.1 | 6.75 | 30 | 1 | 0 | 1 | 215 | 5 | 25 | 20 |
| 444 | SM(d18:1_20:0)_SM(d16:1_22:0) | $[M+H]^+$ | 759.6 | 184.1 | 7.9 | 30 | 1 | 0 | 1 | 215 | 8 | 11 | 21 |
| 445 | SM(d18:1_22:0)_SM(d16:1_24:0) | $[M+H]^+$ | 787.7 | 184.1 | 9.2 | 30 | 1 | 0 | 1 | 215 | 8 | 11 | 21 |
| 446 | SM(d18:1_24:0) | $[M+H]^+$ | 815.7 | 184.1 | 10.15 | 30 | 1 | 0 | 1 | 215 | 8 | 11 | 21 |
| 447 | SM(d18:1_24:1) | $[M+H]^+$ | 813.7 | 184.1 | 9.15 | 60 | 1 | 0 | 1 | 215 | 8 | 11 | 21 |
| 448 | SM(d18:2_14:0) | $[M+H]^+$ | 673.5 | 184.1 | 4.47 | 24 | 1 | 0 | 1 | 215 | 8 | 11 | 21 |
| 449 | SM(d18:2_16:0) | $[M+H]^+$ | 701.6 | 184.1 | 5.18 | 24 | 1 | 0 | 1 | 215 | 8 | 11 | 21 |
| 450 | SM(d18:2_18:0) | $[M+H]^+$ | 729.6 | 184.1 | 6 | 36 | 1 | 0 | 1 | 215 | 8 | 11 | 21 |
| 451 | SM(d18:2_18:1) | $[M+H]^+$ | 727.6 | 184.1 | 5.35 | 30 | 1 | 0 | 1 | 215 | 8 | 11 | 21 |
| 452 | SM(d18:2_20:0) | $[M+H]^+$ | 757.6 | 184.1 | 7.05 | 30 | 1 | 0 | 1 | 215 | 8 | 11 | 21 |
| 453 | Sph(d16:1) | $[M+H]^+$ | 272.3 | 254.3 | 1.75 | 24 | 1 | 0 | 5 | 45 | 8 | 11 | 21 |
| 454 | Sph(d18:1) | $[M+H]^+$ | 300.3 | 282.3 | 2.4 | 24 | 1 | 0 | 1 | 45 | 8 | 11 | 21 |
| 455 | Sph(d18:2) | $[M+H]^+$ | 298.3 | 280.3 | 1.7 | 36 | 1 | 0 | 5 | 45 | 8 | 11 | 21 |
| 456 | Sulfa (d18:1:_16:0(OH)) | $[M+H]^+$ | 796.8 | 264.3 | 4.75 | 18 | 1 | 0 | 5 | 45 | 8 | 11 | 21 |
| 457 | Sulfa (d18:1:_16:0) | $[M+H]^+$ | 780.8 | 264.3 | 4.9 | 18 | 1 | 0 | 1 | 45 | 8 | 11 | 21 |
| 458 | Sulfa (d18:1:_18:0(OH)) | $[M+H]^+$ | 824.8 | 264.3 | 6.1 | 24 | 1 | 0 | 5 | 45 | 8 | 11 | 21 |
| 459 | Sulfa (d18:1:_18:0) | $[M+H]^+$ | 808.8 | 264.3 | 6.3 | 24 | 1 | 0 | 5 | 45 | 8 | 11 | 21 |
| 460 | Sulfa (d18:1:_24:0(OH)) | $[M+H]^+$ | 908.8 | 264.3 | 8.35 | 18 | 1 | 0 | 5 | 45 | 8 | 11 | 21 |
| 461 | Sulfa (d18:1:_24:0) | $[M+H]^+$ | 892.8 | 264.3 | 8.6 | 24 | 1 | 0 | 5 | 45 | 8 | 11 | 21 |
| 462 | Sulfa (d18:1:_24:1(OH)) | $[M+H]^+$ | 906.8 | 264.3 | 7.25 | 24 | 1 | 0 | 5 | 45 | 8 | 11 | 21 |
| 463 | Sulfa (d18:1:_24:1) | $[M+H]^+$ | 890.7 | 264.303 | 7.6 | 24 | 1 | 0 | 5 | 45 | 8 | 11 | 21 |
| 464 | Sulfa (d18:2:_18:0(OH)) | $[M+H]^+$ | 822.8 | 262.3 | 5.9 | 24 | 1 | 0 | 5 | 45 | 8 | 11 | 21 |
| 465 | Sulfa (d18:2:_18:0) | $[M+H]^+$ | 806.8 | 262.3 | 6.1 | 24 | 1 | 0 | 5 | 45 | 8 | 11 | 21 |
| 466 | TG(48:0) [NL-16:0] | $[M+NH_4]^+$ | 824.8 | 551.5 | 12.2 | 24 | 1 | 0 | 5 | 50 | 8 | 11 | 21 |
| 467 | TG(48:0) [NL-18:0] | $[M+NH_4]^+$ | 824.8 | 523.5 | 12.2 | 24 | 1 | 0 | 5 | 50 | 8 | 11 | 21 |
| 468 | TG(48:0) [SIM] | $[M+NH_4]^+$ | 824.8 | 824.8 | 12.2 | 24 | 1 | 0 | 5 | 50 | 8 | 11 | 21 |
| 469 | TG(48:1) [NL-16:1] | $[M+NH_4]^+$ | 822.8 | 551.5 | 12.32 | 24 | 1 | 0 | 1 | 50 | 8 | 11 | 21 |
| 470 | TG(48:1) [NL-18:1] | $[M+NH_4]^+$ | 822.8 | 523.5 | 12.32 | 24 | 1 | 0 | 1 | 50 | 8 | 11 | 21 |
| 471 | TG(48:1) [SIM] | $[M+NH_4]^+$ | 822.8 | 822.8 | 12.32 | 24 | 1 | 0 | 1 | 50 | 8 | 11 | 21 |
| 472 | TG(48:2) [NL-14:0] | $[M+NH_4]^+$ | 820.8 | 575.5 | 12.03 | 24 | 1 | 0 | 1 | 50 | 8 | 11 | 21 |
| 473 | TG(48:2) [NL-14:1] | $[M+NH_4]^+$ | 820.8 | 577.5 | 12.03 | 24 | 1 | 0 | 1 | 50 | 8 | 11 | 21 |
| 474 | TG(48:2) [NL-16:1] | $[M+NH_4]^+$ | 820.8 | 549.5 | 12.03 | 24 | 1 | 0 | 1 | 50 | 8 | 11 | 21 |
| 475 | TG(48:2) [NL-18:2] | $[M+NH_4]^+$ | 820.8 | 523.5 | 12.03 | 24 | 1 | 0 | 1 | 50 | 8 | 11 | 21 |
| 476 | TG(48:2) [SIM] | $[M+NH_4]^+$ | 820.8 | 820.8 | 12.03 | 24 | 1 | 0 | 1 | 50 | 8 | 11 | 21 |
| 477 | TG(48:3) [NL-14:0] | $[M+NH_4]^+$ | 818.8 | 573.5 | 11.78 | 24 | 1 | 0 | 5 | 50 | 8 | 11 | 21 |
| 478 | TG(48:3) [NL-16:1] | $[M+NH_4]^+$ | 818.8 | 547.5 | 11.78 | 24 | 1 | 0 | 5 | 50 | 8 | 11 | 21 |
| 479 | TG(48:3) [NL-18:3] | $[M+NH_4]^+$ | 818.8 | 523.5 | 11.78 | 24 | 1 | 0 | 5 | 50 | 8 | 11 | 21 |
| 480 | TG(48:3) [SIM] | $[M+NH_4]^+$ | 818.8 | 818.8 | 11.78 | 24 | 1 | 0 | 1 | 50 | 8 | 11 | 21 |
| 481 | TG(49:1) [NL-16:1] | $[M+NH_4]^+$ | 836.8 | 565.5 | 11.9 | 30 | 1 | 0 | 5 | 50 | 8 | 11 | 21 |
| 482 | TG(49:1) [NL-17:1] | $[M+NH_4]^+$ | 836.8 | 551.5 | 11.9 | 30 | 1 | 0 | 5 | 50 | 8 | 11 | 21 |
| 483 | TG(49:1) [SIM] | $[M+NH_4]^+$ | 836.8 | 836.8 | 11.9 | 30 | 1 | 0 | 1 | 50 | 5 | 42 | 19 |
| 484 | TG(50:0) [NL-18:0] | $[M+NH_4]^+$ | 852.8 | 551.5 | 12.6 | 24 | 1 | 0 | 1 | 50 | 5 | 25 | 20 |
| 485 | TG(50:0) [SIM] | $[M+NH_4]^+$ | 852.8 | 852.8 | 12.6 | 24 | 1 | 0 | 1 | 50 | 8 | 11 | 21 |
| 486 | TG(50:1) [NL-14:0] | $[M+NH_4]^+$ | 850.8 | 605.5 | 12.25 | 24 | 1 | 0 | 5 | 50 | 8 | 11 | 21 |
| 487 | TG(50:1) [NL-16:0] | $[M+NH_4]^+$ | 850.8 | 577.5 | 12.25 | 24 | 1 | 0 | 5 | 50 | 8 | 11 | 21 |
| 488 | TG(50:1) [NL-18:1] | $[M+NH_4]^+$ | 850.8 | 551.5 | 12.25 | 24 | 1 | 0 | 1 | 50 | 8 | 11 | 21 |
| 489 | TG(50:1) [SIM] | $[M+NH_4]^+$ | 850.8 | 850.8 | 12.68 | 24 | 1 | 0 | 5 | 50 | 8 | 11 | 21 |
| 490 | TG(50:2) [NL-14:0] | $[M+NH_4]^+$ | 848.8 | 603.5 | 11.85 | 30 | 1 | 0 | 1 | 50 | 5 | 42 | 19 |
| 491 | TG(50:2) [NL-16:1] | $[M+NH_4]^+$ | 848.8 | 577.5 | 11.85 | 30 | 1 | 0 | 1 | 50 | 5 | 25 | 20 |
| 492 | TG(50:2) [NL-18:1] | $[M+NH_4]^+$ | 848.8 | 549.5 | 11.85 | 30 | 1 | 0 | 1 | 50 | 8 | 10.5 | 20.5 |
| 493 | TG(50:2) [NL-18:2] | $[M+NH_4]^+$ | 848.8 | 551.5 | 11.85 | 30 | 1 | 0 | 1 | 50 | 8 | 11 | 21 |
| 494 | TG(50:2) [SIM] | $[M+NH_4]^+$ | 848.8 | 848.8 | 11.85 | 30 | 1 | 0 | 5 | 50 | 8 | 11 | 21 |
| 495 | TG(50:3) [NL-14:0] | $[M+NH_4]^+$ | 846.8 | 601.5 | 11.7 | 24 | 1 | 0 | 1 | 50 | 8 | 11 | 21 |
| 496 | TG(50:3) [NL-14:1] | $[M+NH_4]^+$ | 846.8 | 603.5 | 11.7 | 24 | 1 | 0 | 5 | 50 | 8 | 11 | 21 |
| 497 | TG(50:3) [NL-16:1] | $[M+NH_4]^+$ | 846.8 | 575.5 | 11.7 | 24 | 1 | 0 | 1 | 50 | 8 | 11 | 21 |
| 498 | TG(50:3) [NL-18:2] | $[M+NH_4]^+$ | 846.8 | 549.5 | 11.7 | 24 | 1 | 0 | 1 | 50 | 8 | 11 | 21 |
| 499 | TG(50:3) [NL-18:3] | $[M+NH_4]^+$ | 846.8 | 551.5 | 11.7 | 24 | 1 | 0 | 5 | 50 | 8 | 11 | 21 |
| 500 | TG(50:3) [SIM] | $[M+NH_4]^+$ | 846.8 | 846.8 | 11.7 | 24 | 1 | 0 | 5 | 50 | 8 | 11 | 21 |

| | | | | | | | | | | | | | |
|---|---|---|---|---|---|---|---|---|---|---|---|---|---|
| 501 | TG(50:4) [NL-14:0] | $[M+NH_4]^+$ | 844.8 | 599.5 | 11.85 | 24 | 1 | 0 | 5 | 50 | 8 | 11 | 21 |
| 502 | TG(50:4) [NL-18:3] | $[M+NH_4]^+$ | 844.8 | 549.5 | 11.85 | 24 | 1 | 0 | 5 | 50 | 5 | 42 | 19 |
| 503 | TG(50:4) [NL-20:4] | $[M+NH_4]^+$ | 844.8 | 523.5 | 11.98 | 24 | 1 | 0 | 5 | 50 | 8 | 11 | 21 |
| 504 | TG(50:4) [SIM] | $[M+NH_4]^+$ | 844.8 | 844.8 | 11.85 | 24 | 1 | 0 | 1 | 50 | 8 | 11 | 21 |
| 505 | TG(51:0) [NL-16:0] | $[M+NH_4]^+$ | 866.7 | 593.4 | 13.1 | 36 | 1 | 0 | 5 | 50 | 8 | 11 | 21 |
| 506 | TG(51:0) [SIM] | $[M+NH_4]^+$ | 866.7 | 866.7 | 13.1 | 36 | 1 | 0 | 1 | 50 | 8 | 11 | 21 |
| 507 | TG(51:1) [NL-17:0] | $[M+NH_4]^+$ | 864.8 | 577.5 | 12.4 | 24 | 1 | 0 | 5 | 50 | 8 | 11 | 21 |
| 508 | TG(51:1) [SIM] | $[M+NH_4]^+$ | 864.8 | 864.8 | 12.8 | 48 | 1 | 0 | 1 | 50 | 5 | 42 | 19 |
| 509 | TG(51:2) [NL-15:0] | $[M+NH_4]^+$ | 862.8 | 603.5 | 12 | 30 | 1 | 0 | 5 | 50 | 8 | 11 | 21 |
| 510 | TG(51:2) [NL-17:0] | $[M+NH_4]^+$ | 862.8 | 575.5 | 12 | 30 | 1 | 0 | 5 | 50 | 8 | 11 | 21 |
| 511 | TG(51:2) [NL-17:1] | $[M+NH_4]^+$ | 862.8 | 577.5 | 12 | 30 | 1 | 0 | 1 | 50 | 8 | 11 | 21 |
| 512 | TG(51:2) [SIM] | $[M+NH_4]^+$ | 862.8 | 862.8 | 12 | 30 | 1 | 0 | 1 | 50 | 8 | 11 | 21 |
| 513 | TG(52:1) [NL-18:0] | $[M+NH_4]^+$ | 878.8 | 577.5 | 13.07 | 24 | 1 | 0 | 5 | 50 | 8 | 11 | 21 |
| 514 | TG(52:1) [NL-18:1] | $[M+NH_4]^+$ | 878.8 | 579.5 | 13.07 | 24 | 1 | 0 | 1 | 50 | 8 | 11 | 21 |
| 515 | TG(52:1) [SIM] | $[M+NH_4]^+$ | 878.8 | 878.8 | 12.65 | 24 | 1 | 0 | 1 | 50 | 5 | 42 | 19 |
| 516 | TG(52:2) [NL-16:0] | $[M+NH_4]^+$ | 876.8 | 603.5 | 12.3 | 24 | 1 | 0 | 5 | 50 | 8 | 11 | 21 |
| 517 | TG(52:2) [NL-18:2] | $[M+NH_4]^+$ | 876.8 | 579.5 | 12.3 | 24 | 1 | 0 | 1 | 50 | 8 | 11 | 21 |
| 518 | TG(52:2) [SIM] | $[M+NH_4]^+$ | 876.8 | 876.8 | 12.3 | 24 | 1 | 0 | 1 | 50 | 8 | 11 | 21 |
| 519 | TG(52:3) [NL-16:1] | $[M+NH_4]^+$ | 874.8 | 603.5 | 12 | 24 | 1 | 0 | 1 | 50 | 8 | 11 | 21 |
| 520 | TG(52:3) [NL-18:2] | $[M+NH_4]^+$ | 874.8 | 577.5 | 12 | 24 | 1 | 0 | 1 | 50 | 8 | 11 | 21 |
| 521 | TG(52:3) [SIM] | $[M+NH_4]^+$ | 874.8 | 874.8 | 12 | 24 | 1 | 0 | 5 | 50 | 5 | 42 | 19 |
| 522 | TG(52:4) [NL-16:1] | $[M+NH_4]^+$ | 872.8 | 601.5 | 12.13 | 24 | 1 | 0 | 1 | 50 | 8 | 11 | 21 |
| 523 | TG(52:4) [NL-18:2] | $[M+NH_4]^+$ | 872.8 | 575.5 | 12.13 | 24 | 1 | 0 | 1 | 50 | 5 | 42 | 19 |
| 524 | TG(52:4) [NL-18:3] | $[M+NH_4]^+$ | 872.8 | 577.5 | 12.13 | 24 | 1 | 0 | 1 | 50 | 8 | 11 | 21 |
| 525 | TG(52:4) [SIM] | $[M+NH_4]^+$ | 872.8 | 872.8 | 12.1 | 30 | 1 | 0 | 1 | 50 | 8 | 11 | 21 |
| 526 | TG(52:5) [NL-18:3] | $[M+NH_4]^+$ | 870.8 | 575.5 | 11.91 | 24 | 1 | 0 | 5 | 50 | 8 | 11 | 21 |
| 527 | TG(52:5) [NL-20:4] | $[M+NH_4]^+$ | 870.8 | 549.5 | 12.02 | 24 | 1 | 0 | 5 | 50 | 8 | 11 | 21 |
| 528 | TG(52:5) [NL-20:5] | $[M+NH_4]^+$ | 870.8 | 551.5 | 12.08 | 24 | 1 | 0 | 5 | 50 | 5 | 42 | 19 |
| 529 | TG(52:5) [SIM] | $[M+NH_4]^+$ | 870.8 | 870.8 | 11.91 | 24 | 1 | 0 | 1 | 50 | 5 | 25 | 20 |
| 530 | TG(53:2) [NL-17:1] | $[M+NH_4]^+$ | 890.8 | 605.5 | 12.3 | 24 | 1 | 0 | 5 | 50 | 8 | 34 | 15 |
| 531 | TG(53:2) [NL-18:1] | $[M+NH_4]^+$ | 890.8 | 591.5 | 12.3 | 24 | 1 | 0 | 1 | 50 | 5 | 42 | 19 |
| 532 | TG(53:2) [SIM] | $[M+NH_4]^+$ | 890.8 | 890.8 | 12.3 | 36 | 1 | 0 | 1 | 50 | 5 | 42 | 19 |
| 533 | TG(54:0) [NL-18:0] | $[M+NH_4]^+$ | 908.8 | 607.5 | 13.2 | 36 | 1 | 0 | 5 | 50 | 8 | 11 | 21 |
| 534 | TG(54:0) [SIM] | $[M+NH_4]^+$ | 908.8 | 908.8 | 13.2 | 36 | 1 | 0 | 1 | 50 | 8 | 11 | 21 |
| 535 | TG(54:1) [NL-18:1] | $[M+NH_4]^+$ | 906.8 | 607.5 | 13 | 30 | 1 | 0 | 5 | 50 | 8 | 11 | 21 |
| 536 | TG(54:1) [SIM] | $[M+NH_4]^+$ | 906.8 | 906.8 | 13 | 30 | 1 | 0 | 1 | 50 | 8 | 11 | 21 |
| 537 | TG(54:2) [NL-18:0] | $[M+NH_4]^+$ | 904.8 | 603.5 | 12.65 | 24 | 1 | 0 | 5 | 50 | 8 | 11 | 21 |
| 538 | TG(54:2) [NL-20:1] | $[M+NH_4]^+$ | 904.8 | 577.5 | 12.65 | 24 | 1 | 0 | 5 | 50 | 8 | 11 | 21 |
| 539 | TG(54:2) [SIM] | $[M+NH_4]^+$ | 904.8 | 904.8 | 12.65 | 24 | 1 | 0 | 1 | 50 | 8 | 11 | 21 |
| 540 | TG(54:3) [NL-18:1] | $[M+NH_4]^+$ | 902.8 | 603.5 | 12.3 | 24 | 1 | 0 | 1 | 50 | 5 | 42 | 19 |
| 541 | TG(54:3) [NL-18:2] | $[M+NH_4]^+$ | 902.8 | 605.5 | 12.3 | 24 | 1 | 0 | 5 | 50 | 8 | 11 | 21 |
| 542 | TG(54:3) [SIM] | $[M+NH_4]^+$ | 902.8 | 902.8 | 12.3 | 24 | 1 | 0 | 1 | 50 | 8 | 11 | 21 |
| 543 | TG(54:4) [NL-18:2] | $[M+NH_4]^+$ | 900.8 | 603.5 | 11.9 | 24 | 1 | 0 | 1 | 50 | 8 | 11 | 21 |
| 544 | TG(54:4) [NL-20:3] | $[M+NH_4]^+$ | 900.8 | 577.5 | 11.9 | 24 | 1 | 0 | 1 | 50 | 8 | 11 | 21 |
| 545 | TG(54:4) [SIM] | $[M+NH_4]^+$ | 900.8 | 900.8 | 11.9 | 24 | 1 | 0 | 1 | 50 | 8 | 11 | 21 |
| 546 | TG(54:5) [NL-18:3] | $[M+NH_4]^+$ | 898.8 | 603.5 | 12.23 | 24 | 1 | 0 | 5 | 50 | 8 | 11 | 21 |
| 547 | TG(54:5) [NL-20:4] | $[M+NH_4]^+$ | 898.8 | 577.5 | 12.23 | 24 | 1 | 0 | 1 | 50 | 8 | 11 | 21 |
| 548 | TG(54:5) [SIM] | $[M+NH_4]^+$ | 898.8 | 898.8 | 12.2 | 24 | 1 | 0 | 1 | 50 | 8 | 11 | 21 |
| 549 | TG(54:6) [NL-18:3] | $[M+NH_4]^+$ | 896.8 | 601.5 | 11.96 | 24 | 1 | 0 | 5 | 50 | 8 | 11 | 21 |
| 550 | TG(54:6) [NL-20:4] | $[M+NH_4]^+$ | 896.8 | 575.5 | 12.08 | 24 | 1 | 0 | 5 | 50 | 8 | 11 | 21 |
| 551 | TG(54:6) [NL-20:5] | $[M+NH_4]^+$ | 896.8 | 577.5 | 12.1 | 24 | 1 | 0 | 5 | 50 | 8 | 11 | 21 |
| 552 | TG(54:6) [NL-22:6] | $[M+NH_4]^+$ | 896.8 | 551.5 | 11.8 | 24 | 1 | 0 | 5 | 50 | 5 | 42 | 19 |
| 553 | TG(54:6) [SIM] | $[M+NH_4]^+$ | 896.8 | 896.8 | 11.9 | 36 | 1 | 0 | 5 | 50 | 5 | 42 | 19 |
| 554 | TG(54:7) [NL-20:5] | $[M+NH_4]^+$ | 894.8 | 575.5 | 11.85 | 24 | 1 | 0 | 5 | 50 | 8 | 11 | 21 |
| 555 | TG(54:7) [NL-22:6] | $[M+NH_4]^+$ | 894.8 | 549.5 | 11.91 | 24 | 1 | 0 | 5 | 50 | 5 | 42 | 19 |
| 556 | TG(54:7) [SIM] | $[M+NH_4]^+$ | 894.8 | 894.8 | 11.7 | 36 | 1 | 0 | 1 | 50 | 8 | 11 | 21 |
| 557 | TG(56:6) [NL-20:4] | $[M+NH_4]^+$ | 924.8 | 603.5 | 12.4 | 24 | 1 | 0 | 5 | 50 | 8 | 11 | 21 |
| 558 | TG(56:6) [NL-22:5](a_b) | $[M+NH_4]^+$ | 924.8 | 577.5 | 12.33 | 36 | 1 | 0 | 5 | 50 | 8 | 11 | 21 |
| 559 | TG(56:6) [SIM] | $[M+NH_4]^+$ | 924.8 | 924.8 | 12.3 | 36 | 1 | 0 | 1 | 50 | 8 | 11 | 21 |
| 560 | TG(56:7) [NL-20:4] | $[M+NH_4]^+$ | 922.8 | 601.5 | 12.15 | 24 | 1 | 0 | 5 | 50 | 8 | 11 | 21 |
| 561 | TG(56:7) [NL-20:5] | $[M+NH_4]^+$ | 922.8 | 603.5 | 12.15 | 24 | 1 | 0 | 5 | 50 | 8 | 11 | 21 |
| 562 | TG(56:7) [NL-22:5](a_b) | $[M+NH_4]^+$ | 922.8 | 575.5 | 12.03 | 36 | 1 | 0 | 5 | 50 | 8 | 11 | 21 |
| 563 | TG(56:7) [NL-22:6] | $[M+NH_4]^+$ | 922.8 | 577.5 | 11.8 | 24 | 1 | 0 | 1 | 50 | 5 | 42 | 19 |
| 564 | TG(56:7) [SIM] | $[M+NH_4]^+$ | 922.8 | 922.8 | 12.05 | 36 | 1 | 0 | 1 | 50 | 8 | 11 | 21 |
| 565 | TG(56:8) [NL-20:4] | $[M+NH_4]^+$ | 920.8 | 599.5 | 11.9 | 24 | 1 | 0 | 5 | 50 | 8 | 11 | 21 |
| 566 | TG(56:8) [NL-20:5] | $[M+NH_4]^+$ | 920.8 | 601.5 | 11.5 | 18 | 1 | 0 | 1 | 50 | 5 | 42 | 19 |
| 567 | TG(56:8) [NL-22:6] | $[M+NH_4]^+$ | 920.8 | 575.5 | 11.6 | 18 | 1 | 0 | 1 | 50 | 8 | 11 | 21 |
| 568 | TG(56:8) [SIM] | $[M+NH_4]^+$ | 920.8 | 920.8 | 11.85 | 30 | 1 | 0 | 5 | 50 | 8 | 11 | 21 |
| 569 | TG(56:9) [NL-22:6] | $[M+NH_4]^+$ | 918.8 | 573.5 | 11.77 | 18 | 1 | 0 | 5 | 50 | 8 | 11 | 21 |
| 570 | TG(56:9) [SIM] | $[M+NH_4]^+$ | 918.8 | 918.8 | 11.63 | 30 | 1 | 0 | 1 | 50 | 5 | 42 | 19 |
| 571 | TG(58:10) [NL-22:6] | $[M+NH_4]^+$ | 944.9 | 599.5 | 11.55 | 30 | 1 | 0 | 5 | 50 | 5 | 42 | 19 |
| 572 | TG(58:10) [SIM] | $[M+NH_4]^+$ | 944.9 | 944.9 | 11.55 | 30 | 1 | 0 | 1 | 50 | 8 | 11 | 21 |
| 573 | TG(58:8) [NL-22:6] | $[M+NH_4]^+$ | 948.8 | 603.5 | 12.3 | 24 | 1 | 0 | 5 | 50 | 8 | 11 | 21 |

| | | | | | | | | | | | | | |
|---|---|---|---|---|---|---|---|---|---|---|---|---|---|
| 574 | TG(58:8) [SIM] | $[M+NH_4]^+$ | 948.8 | 948.8 | 12.3 | 24 | 1 | 0 | 1 | 50 | 8 | 11 | 21 |
| 575 | TG(58:9) [NL-22:6] | $[M+NH_4]^+$ | 946.9 | 601.5 | 12 | 36 | 1 | 0 | 5 | 50 | 8 | 11 | 21 |
| 576 | TG(58:9) [SIM] | $[M+NH_4]^+$ | 946.9 | 946.9 | 12 | 24 | 1 | 0 | 1 | 50 | 8 | 11 | 21 |
| 577 | TG(O-50:1) [NL-15:0] | $[M+NH_4]^+$ | 836.8 | 577.5 | 12.85 | 18 | 1 | 0 | 5 | 50 | 5 | 42 | 19 |
| 578 | TG(O-50:1) [NL-16:0] | $[M+NH_4]^+$ | 836.8 | 563.5 | 12.85 | 18 | 1 | 0 | 5 | 50 | 5 | 42 | 19 |
| 579 | TG(O-50:1) [NL-18:1] | $[M+NH_4]^+$ | 836.8 | 537.5 | 12.85 | 18 | 1 | 0 | 5 | 50 | 8 | 11 | 21 |
| 580 | TG(O-50:2) [NL-16:1] | $[M+NH_4]^+$ | 834.8 | 563.503 | 12.75 | 18 | 1 | 0 | 1 | 50 | 8 | 11 | 21 |
| 581 | TG(O-50:2) [NL-18:1] | $[M+NH_4]^+$ | 834.8 | 535.5 | 12.75 | 18 | 1 | 0 | 5 | 50 | 8 | 11 | 21 |
| 582 | TG(O-50:2) [NL-18:2] | $[M+NH_4]^+$ | 834.8 | 537.5 | 12.75 | 18 | 1 | 0 | 5 | 50 | 8 | 11 | 21 |
| 583 | TG(O-50:2) [SIM] | $[M+NH_4]^+$ | 834.8 | 834.8 | 12.75 | 18 | 1 | 0 | 1 | 50 | 8 | 11 | 21 |
| 584 | TG(O-50:3) [NL-18:2] | $[M+NH_4]^+$ | 832.8 | 535.5 | 12.4 | 18 | 1 | 0 | 5 | 50 | 5 | 42 | 19 |
| 585 | TG(O-50:3) [SIM] | $[M+NH_4]^+$ | 832.8 | 832.8 | 12.4 | 18 | 1 | 0 | 1 | 50 | 5 | 42 | 19 |
| 586 | TG(O-52:1) [NL-16:0] | $[M+NH_4]^+$ | 864.8 | 591.5 | 13.25 | 18 | 1 | 0 | 5 | 50 | 8 | 11 | 21 |
| 587 | TG(O-52:1) [NL-18:1] | $[M+NH_4]^+$ | 864.8 | 565.5 | 13.25 | 18 | 1 | 0 | 1 | 50 | 8 | 11 | 21 |
| 588 | TG(O-52:2) [NL-16:0] | $[M+NH_4]^+$ | 862.8 | 589.5 | 12.88 | 18 | 1 | 0 | 5 | 50 | 5 | 42 | 19 |
| 589 | TG(O-52:2) [NL-18:1] | $[M+NH_4]^+$ | 862.8 | 563.5 | 12.88 | 18 | 1 | 0 | 1 | 50 | 8 | 11 | 21 |
| 590 | TG(O-54:2) [NL-17:1] | $[M+NH_4]^+$ | 890.8 | 605.503 | 13.4 | 24 | 1 | 0 | 5 | 50 | 8 | 11 | 21 |
| 591 | TG(O-54:2) [NL-18:1] | $[M+NH_4]^+$ | 890.8 | 591.503 | 13.4 | 24 | 1 | 0 | 5 | 50 | 8 | 11 | 21 |
| 592 | TG(O-54:3) [NL-17:1] | $[M+NH_4]^+$ | 888.8 | 603.5 | 13 | 24 | 1 | 0 | 5 | 50 | 8 | 11 | 21 |
| 593 | TG(O-54:3) [NL-18:1] | $[M+NH_4]^+$ | 888.8 | 589.5 | 13 | 24 | 1 | 0 | 5 | 50 | 5 | 42 | 19 |
| 594 | TG(O-54:3) [SIM] | $[M+NH_4]^+$ | 888.8 | 888.8 | 13 | 24 | 1 | 0 | 1 | 50 | 5 | 42 | 19 |
| 595 | TG(O-54:4) [NL-17:1] | $[M+NH_4]^+$ | 886.8 | 601.5 | 11.8 | 30 | 1 | 0 | 1 | 50 | 5 | 42 | 19 |
| 596 | TG(O-54:4) [NL-18:2] | $[M+NH_4]^+$ | 886.8 | 589.5 | 12.3 | 24 | 1 | 0 | 1 | 50 | 8 | 11 | 21 |
| 597 | TG(O-54:4) [SIM] | $[M+NH_4]^+$ | 886.8 | 886.8 | 12.7 | 24 | 1 | 0 | 5 | 50 | 8 | 11 | 21 |
| 598 | Ubiquinone | $[M+NH_4]^+$ | 880.7 | 197 | 11.75 | 24 | 1 | 0 | 5 | 60 | 5 | 42 | 19 |

Supplemental Table 4: Plasma metabolite concentration measured by biocrates MxP Quant 500

| Metabolites | Full Cohort (n = 361) | LDLc 10th Percentile (n = 36) | LDLc 90th Percentile (n = 36) |
|---|---|---|---|
| | Geometric mean ± SD (µM) | Mean ± SD (µM) | Mean ± SD (µM) |
| C0 | 37.8 ± 1.56 | 37.94 ± 18.01 | 40.22 ± 16.77 |
| C2 | 4.79 ± 1.43 | 5.14 ± 2.24 | 5.65 ± 1.90 |
| C3 | 0.369 ± 1.4 | 0.39 ± 0.15 | 0.42 ± 0.12 |
| C4 | 0.157 ± 1.45 | 0.17 ± 0.07 | 0.18 ± 0.05 |
| C6 (C4:1-DC) | 0.0763 ± 1.59 | 0.08 ± 0.04 | 0.08 ± 0.03 |
| C18:1 | 0.0967 ± 1.31 | 0.10 ± 0.03 | 0.11 ± 0.03 |
| TMAO | 3.95 ± 1.9 | 4.34 ± 2.77 | 5.64 ± 3.35 |
| Ala | 402 ± 1.31 | 378.06 ± 97.80 | 454.33 ± 144.15 |
| Arg | 97.1 ± 1.25 | 93.56 ± 21.99 | 101.06 ± 22.41 |
| Asn | 58.2 ± 1.26 | 59.52 ± 13.78 | 55.47 ± 12.32 |
| Asp | 4.7 ± 2.11 | 5.48 ± 3.13 | 6.12 ± 3.55 |
| Cys | 85.9 ± 1.27 | 85.40 ± 18.27 | 92.12 ± 24.52 |
| Gln | 672 ± 1.2 | 670.72 ± 131.13 | 664.64 ± 119.00 |
| Glu | 47.8 ± 1.63 | 52.31 ± 29.60 | 60.85 ± 29.61 |
| Gly | 287 ± 1.33 | 301.22 ± 65.77 | 271.00 ± 95.04 |
| His | 93.7 ± 1.22 | 94.79 ± 15.34 | 91.45 ± 22.82 |
| Ile | 71.2 ± 1.29 | 73.09 ± 19.85 | 76.41 ± 19.88 |
| Leu | 122 ± 1.25 | 124.07 ± 27.84 | 130.51 ± 29.63 |
| Lys | 193 ± 1.21 | 187.28 ± 31.11 | 202.19 ± 41.27 |
| Met | 29.3 ± 1.23 | 29.96 ± 5.99 | 28.11 ± 5.90 |
| Phe | 60.1 ± 1.26 | 60.51 ± 12.70 | 61.59 ± 16.26 |
| Pro | 164 ± 1.37 | 167.18 ± 58.39 | 181.99 ± 65.41 |
| Ser | 103 ± 1.43 | 116.33 ± 30.85 | 94.93 ± 32.80 |
| Thr | 135 ± 1.26 | 139.34 ± 33.49 | 130.27 ± 36.74 |
| Trp | 62.2 ± 1.23 | 63.23 ± 11.40 | 62.98 ± 13.10 |
| Tyr | 70 ± 1.26 | 67.84 ± 14.00 | 75.67 ± 22.85 |
| Val | 212 ± 1.28 | 212.28 ± 48.31 | 225.97 ± 48.59 |
| 1-Met-His | 3.99 ± 1.35 | 4.02 ± 1.30 | 4.37 ± 0.97 |
| 3-Met-His | 2.32 ± 3.08 | 2.80 ± 3.20 | 4.84 ± 4.48 |
| 5-AVA | 0.032 ± 1.66 | 0.04 ± 0.02 | 0.03 ± 0.01 |
| AABA | 14.8 ± 1.5 | 15.02 ± 5.16 | 16.03 ± 6.17 |
| ADMA | 0.447 ± 1.37 | 0.47 ± 0.14 | 0.46 ± 0.19 |
| alpha-AAA | 0.726 ± 1.71 | 0.78 ± 0.37 | 0.92 ± 0.34 |
| BABA | 0.0328 ± 1.94 | 0.05 ± 0.03 | 0.04 ± 0.03 |
| Betaine | 11 ± 3.69 | 23.47 ± 21.70 | 17.39 ± 17.50 |
| Cit | 31.4 ± 1.31 | 30.36 ± 8.42 | 33.41 ± 8.28 |
| Creatinine | 69.8 ± 1.32 | 67.06 ± 17.63 | 75.08 ± 17.88 |
| Cystine | 63.4 ± 1.53 | 61.00 ± 23.04 | 74.24 ± 25.66 |
| HArg | 3.2 ± 1.52 | 3.43 ± 1.02 | 3.57 ± 1.27 |
| HCys | 5.75 ± 1.39 | 6.17 ± 1.96 | 6.22 ± 1.80 |
| Kynurenine | 1.73 ± 1.37 | 1.66 ± 0.60 | 2.02 ± 0.72 |
| Met-SO | 0.405 ± 2.24 | 0.48 ± 0.20 | 0.42 ± 0.14 |
| Orn | 59.1 ± 1.32 | 59.03 ± 17.27 | 60.46 ± 15.82 |
| ProBetaine | 2.55 ± 2.63 | 4.74 ± 5.92 | 3.84 ± 5.14 |
| Sarcosine | 3.77 ± 1.43 | 3.76 ± 1.34 | 4.10 ± 1.50 |
| SDMA | 0.554 ± 1.22 | 0.52 ± 0.10 | 0.55 ± 0.11 |
| t4-OH-Pro | 8.46 ± 1.52 | 9.49 ± 3.77 | 9.44 ± 5.59 |
| Taurine | 99.9 ± 1.43 | 95.40 ± 33.20 | 111.69 ± 40.50 |
| TrpBetaine | 0.593 ± 3.31 | 1.03 ± 1.01 | 1.05 ± 1.34 |
| CA | 0.0399 ± 2.64 | 0.06 ± 0.06 | 0.08 ± 0.10 |
| CDCA | 0.199 ± 1.27 | 0.18 ± 0.04 | 0.22 ± 0.04 |
| DCA | 0.261 ± 1.27 | 0.24 ± 0.05 | 0.28 ± 0.05 |

| | | | |
|---|---|---|---|
| GCA | 0.135 ± 2.5 | 1.08 ± 5.24 | 0.19 ± 0.22 |
| GCDCA | 0.242 ± 2.93 | 0.84 ± 2.38 | 0.30 ± 0.29 |
| GDCA | 0.09 ± 2.89 | 0.31 ± 0.85 | 0.11 ± 0.11 |
| GLCA | 0.00979 ± 2.94 | 0.03 ± 0.06 | 0.02 ± 0.04 |
| GLCAS | 0.24 ± 2.66 | 0.43 ± 0.34 | 0.35 ± 0.32 |
| GUDCA | 0.0532 ± 2.49 | 0.12 ± 0.25 | 0.07 ± 0.10 |
| TCA | 0.0481 ± 2.37 | 0.37 ± 1.87 | 0.06 ± 0.04 |
| TCDCA | 0.0445 ± 2.76 | 0.19 ± 0.73 | 0.05 ± 0.04 |
| TDCA | 0.0409 ± 2.87 | 0.18 ± 0.72 | 0.05 ± 0.04 |
| TLCA | 0.00394 ± 2.32 | 0.01 ± 0.01 | 0.00 ± 0.00 |
| TMCA | 0.00626 ± 3.19 | 0.02 ± 0.04 | 0.01 ± 0.01 |
| beta-Ala | 2.91 ± 1.38 | 2.82 ± 0.79 | 3.05 ± 1.12 |
| GABA | 0.166 ± 1.29 | 0.17 ± 0.04 | 0.18 ± 0.04 |
| Putrescine | 0.153 ± 1.42 | 0.16 ± 0.07 | 0.17 ± 0.05 |
| Serotonin | 0.494 ± 2.29 | 0.85 ± 1.54 | 0.66 ± 0.48 |
| Acocid | 7.27 ± 1.58 | 7.79 ± 2.70 | 8.63 ± 2.26 |
| Lac | 1760 ± 1.56 | 2190 ± 1720 | 2140 ± 1210 |
| Cer(d16:1/20:0) | 0.0736 ± 1.62 | 0.06 ± 0.03 | 0.11 ± 0.04 |
| Cer(d16:1/22:0) | 0.343 ± 1.6 | 0.28 ± 0.10 | 0.52 ± 0.20 |
| Cer(d16:1/23:0) | 0.144 ± 1.66 | 0.11 ± 0.04 | 0.21 ± 0.09 |
| Cer(d16:1/24:0) | 0.208 ± 1.6 | 0.15 ± 0.06 | 0.33 ± 0.13 |
| Cer(d18:1/16:0) | 0.46 ± 1.27 | 0.41 ± 0.09 | 0.55 ± 0.11 |
| Cer(d18:1/18:0) | 0.157 ± 1.34 | 0.14 ± 0.04 | 0.21 ± 0.06 |
| Cer(d18:1/20:0) | 0.152 ± 1.42 | 0.14 ± 0.06 | 0.21 ± 0.07 |
| Cer(d18:1/22:0) | 0.84 ± 1.32 | 0.71 ± 0.18 | 1.17 ± 0.26 |
| Cer(d18:1/23:0) | 0.853 ± 1.32 | 0.67 ± 0.15 | 1.15 ± 0.35 |
| Cer(d18:1/24:0) | 2.3 ± 1.33 | 1.84 ± 0.44 | 3.27 ± 0.73 |
| Cer(d18:1/24:1) | 1.02 ± 1.26 | 0.88 ± 0.21 | 1.21 ± 0.24 |
| Cer(d18:1/25:0) | 0.378 ± 1.36 | 0.32 ± 0.08 | 0.48 ± 0.17 |
| Cer(d18:2/16:0) | 0.109 ± 1.41 | 0.10 ± 0.03 | 0.14 ± 0.04 |
| Cer(d18:2/20:0) | 0.0764 ± 1.54 | 0.07 ± 0.03 | 0.09 ± 0.03 |
| Cer(d18:2/22:0) | 0.266 ± 1.4 | 0.23 ± 0.07 | 0.36 ± 0.12 |
| Cer(d18:2/23:0) | 0.132 ± 1.52 | 0.10 ± 0.04 | 0.17 ± 0.07 |
| Cer(d18:2/24:0) | 0.507 ± 1.37 | 0.44 ± 0.13 | 0.66 ± 0.21 |
| Cer(d18:2/24:1) | 0.228 ± 1.35 | 0.20 ± 0.06 | 0.27 ± 0.08 |
| CE(14:0) | 10.9 ± 1.66 | 8.40 ± 4.58 | 17.75 ± 10.10 |
| CE(15:0) | 3.35 ± 1.61 | 3.14 ± 1.63 | 4.39 ± 1.93 |
| CE(16:0) | 145 ± 1.47 | 111.24 ± 34.45 | 199.72 ± 96.97 |
| CE(16:1) | 32 ± 1.69 | 23.66 ± 9.82 | 55.22 ± 32.87 |
| CE(17:1) | 2.12 ± 1.9 | 1.75 ± 0.87 | 3.40 ± 1.56 |
| CE(18:0) | 8.86 ± 1.47 | 6.88 ± 1.97 | 12.84 ± 5.43 |
| CE(18:1) | 220 ± 1.41 | 169.42 ± 46.58 | 307.06 ± 88.52 |
| CE(18:2) | 1010 ± 1.4 | 766.44 ± 203.29 | 1363.86 ± 404.45 |
| CE(18:3) | 38.7 ± 1.54 | 29.25 ± 10.01 | 62.12 ± 36.32 |
| CE(20:3) | 14.5 ± 1.52 | 11.19 ± 3.75 | 23.27 ± 10.17 |
| CE(20:4) | 173 ± 1.49 | 132.86 ± 57.36 | 251.91 ± 98.86 |
| CE(20:5) | 32.5 ± 1.87 | 23.05 ± 11.49 | 63.87 ± 39.82 |
| CE(22:5) | 1.72 ± 1.46 | 1.31 ± 0.54 | 2.47 ± 1.06 |
| CE(22:6) | 24.6 ± 1.55 | 18.68 ± 7.93 | 37.58 ± 17.15 |
| p-Cresol-SO4 | 24.2 ± 2.65 | 32.81 ± 23.44 | 33.82 ± 24.64 |
| DG(16:0_18:1) | 2.27 ± 1.64 | 2.25 ± 1.26 | 3.44 ± 2.00 |
| DG(16:0_18:2) | 1.36 ± 1.57 | 1.21 ± 0.44 | 1.99 ± 1.02 |
| DG(16:1_18:2) | 0.496 ± 1.69 | 0.45 ± 0.20 | 0.80 ± 0.41 |
| DG(18:1_18:1) | 2.25 ± 1.56 | 2.06 ± 1.25 | 3.34 ± 1.33 |
| DG(18:2_18:2) | 3.25 ± 1.57 | 2.92 ± 1.41 | 4.25 ± 1.61 |
| FA(18:1) | 135 ± 1.5 | 146.55 ± 51.16 | 165.00 ± 69.17 |

| | | | |
|---|---|---|---|
| FA(18:2) | 33.2 ± 1.51 | 37.22 ± 13.99 | 38.34 ± 15.49 |
| FA(20:3) | 0.645 ± 1.53 | 0.67 ± 0.28 | 0.86 ± 0.33 |
| lysoPC a C16:0 | 86.3 ± 1.47 | 76.78 ± 23.53 | 103.76 ± 38.44 |
| lysoPC a C16:1 | 2.12 ± 1.5 | 1.98 ± 0.60 | 2.57 ± 0.98 |
| lysoPC a C17:0 | 1.6 ± 1.56 | 1.42 ± 0.54 | 1.96 ± 0.97 |
| lysoPC a C18:0 | 28.3 ± 1.48 | 23.92 ± 7.49 | 36.46 ± 14.83 |
| lysoPC a C18:1 | 20.6 ± 1.53 | 19.56 ± 6.54 | 23.46 ± 11.49 |
| lysoPC a C18:2 | 37.4 ± 1.58 | 37.52 ± 13.95 | 40.63 ± 23.06 |
| lysoPC a C20:3 | 2.22 ± 1.51 | 2.25 ± 0.97 | 2.85 ± 1.19 |
| lysoPC a C20:4 | 7.33 ± 1.51 | 7.26 ± 3.14 | 8.16 ± 3.08 |
| PC aa C28:1 | 3.18 ± 1.36 | 2.38 ± 0.50 | 4.22 ± 1.07 |
| PC aa C30:0 | 3.06 ± 1.45 | 2.53 ± 0.75 | 4.26 ± 1.63 |
| PC aa C32:0 | 11.1 ± 1.51 | 10.47 ± 4.53 | 15.01 ± 6.42 |
| PC aa C32:1 | 8.91 ± 1.83 | 8.39 ± 3.78 | 14.84 ± 9.06 |
| PC aa C32:2 | 2.59 ± 1.85 | 2.24 ± 0.84 | 3.99 ± 2.22 |
| PC aa C32:3 | 0.462 ± 1.56 | 0.40 ± 0.14 | 0.60 ± 0.27 |
| PC aa C34:1 | 141 ± 1.48 | 127.28 ± 42.57 | 190.37 ± 64.04 |
| PC aa C34:2 | 313 ± 1.45 | 282.53 ± 101.15 | 390.56 ± 121.34 |
| PC aa C34:3 | 11.9 ± 1.56 | 10.78 ± 3.36 | 16.31 ± 7.94 |
| PC aa C34:4 | 1.28 ± 1.67 | 1.05 ± 0.37 | 1.97 ± 1.15 |
| PC aa C36:0 | 0.98 ± 1.5 | 0.96 ± 0.38 | 1.30 ± 0.50 |
| PC aa C36:1 | 34.6 ± 1.4 | 29.42 ± 9.11 | 48.49 ± 15.32 |
| PC aa C36:2 | 189 ± 1.35 | 164.14 ± 49.01 | 233.67 ± 69.35 |
| PC aa C36:3 | 99.4 ± 1.37 | 85.40 ± 22.94 | 126.85 ± 39.48 |
| PC aa C36:4 | 144 ± 1.4 | 125.49 ± 42.96 | 180.38 ± 55.27 |
| PC aa C36:5 | 12.2 ± 1.66 | 10.51 ± 5.97 | 19.18 ± 9.83 |
| PC aa C36:6 | 0.487 ± 1.61 | 0.41 ± 0.20 | 0.71 ± 0.33 |
| PC aa C38:0 | 1.92 ± 1.4 | 1.75 ± 0.56 | 2.43 ± 0.82 |
| PC aa C38:3 | 34.7 ± 1.37 | 30.11 ± 8.25 | 49.12 ± 20.00 |
| PC aa C38:4 | 87.9 ± 1.36 | 74.94 ± 24.80 | 114.03 ± 39.68 |
| PC aa C38:5 | 38.1 ± 1.35 | 31.72 ± 9.80 | 49.28 ± 18.06 |
| PC aa C38:6 | 50.9 ± 1.46 | 43.10 ± 17.73 | 67.49 ± 23.72 |
| PC aa C40:2 | 0.236 ± 1.39 | 0.20 ± 0.05 | 0.29 ± 0.08 |
| PC aa C40:3 | 0.388 ± 1.3 | 0.35 ± 0.08 | 0.46 ± 0.11 |
| PC aa C40:4 | 2.58 ± 1.33 | 2.34 ± 0.82 | 3.43 ± 1.15 |
| PC aa C40:5 | 6.54 ± 1.36 | 5.54 ± 1.87 | 9.07 ± 4.03 |
| PC aa C40:6 | 16.9 ± 1.43 | 13.97 ± 4.54 | 24.07 ± 10.03 |
| PC aa C42:0 | 0.41 ± 1.35 | 0.40 ± 0.13 | 0.44 ± 0.12 |
| PC aa C42:1 | 0.211 ± 1.36 | 0.20 ± 0.07 | 0.24 ± 0.06 |
| PC aa C42:4 | 0.16 ± 1.34 | 0.15 ± 0.05 | 0.18 ± 0.05 |
| PC aa C42:5 | 0.232 ± 1.35 | 0.21 ± 0.06 | 0.29 ± 0.08 |
| PC aa C42:6 | 0.31 ± 1.3 | 0.28 ± 0.08 | 0.38 ± 0.10 |
| PC ae C30:2 | 0.0963 ± 1.4 | 0.08 ± 0.02 | 0.12 ± 0.04 |
| PC ae C32:1 | 2.27 ± 1.52 | 2.24 ± 0.94 | 2.72 ± 0.94 |
| PC ae C32:2 | 0.629 ± 1.52 | 0.60 ± 0.26 | 0.76 ± 0.27 |
| PC ae C34:0 | 1.03 ± 1.55 | 0.95 ± 0.39 | 1.43 ± 0.69 |
| PC ae C34:1 | 7.29 ± 1.49 | 7.00 ± 2.60 | 9.12 ± 3.35 |
| PC ae C34:2 | 9.08 ± 1.52 | 8.97 ± 3.64 | 10.88 ± 3.89 |
| PC ae C34:3 | 6.6 ± 1.58 | 6.93 ± 3.43 | 7.57 ± 2.71 |
| PC ae C36:1 | 5.53 ± 1.43 | 4.77 ± 1.61 | 7.27 ± 2.38 |
| PC ae C36:2 | 10.1 ± 1.42 | 9.42 ± 3.52 | 12.05 ± 3.93 |
| PC ae C36:3 | 5.53 ± 1.43 | 5.49 ± 1.89 | 6.64 ± 1.96 |
| PC ae C36:4 | 13.1 ± 1.48 | 12.86 ± 5.39 | 16.17 ± 4.59 |
| PC ae C36:5 | 8.62 ± 1.53 | 8.24 ± 3.89 | 10.63 ± 3.09 |
| PC ae C38:0 | 1.38 ± 1.41 | 1.15 ± 0.36 | 1.84 ± 0.60 |
| PC ae C38:1 | 0.188 ± 2.67 | 0.27 ± 0.17 | 0.22 ± 0.14 |

| | | | |
|---|---|---|---|
| PC ae C38:2 | 1.47 ± 1.39 | 1.34 ± 0.41 | 1.72 ± 0.54 |
| PC ae C38:3 | 2.83 ± 1.36 | 2.56 ± 0.78 | 3.41 ± 1.05 |
| PC ae C38:4 | 10.3 ± 1.35 | 9.70 ± 3.22 | 12.14 ± 3.16 |
| PC ae C38:5 | 14 ± 1.35 | 13.47 ± 4.65 | 16.15 ± 3.59 |
| PC ae C38:6 | 5.06 ± 1.41 | 4.58 ± 1.70 | 6.28 ± 1.74 |
| PC ae C40:1 | 1.06 ± 1.36 | 0.88 ± 0.22 | 1.38 ± 0.47 |
| PC ae C40:2 | 1.32 ± 1.34 | 1.11 ± 0.34 | 1.66 ± 0.41 |
| PC ae C40:3 | 0.808 ± 1.32 | 0.77 ± 0.22 | 0.89 ± 0.23 |
| PC ae C40:4 | 1.87 ± 1.29 | 1.79 ± 0.46 | 2.05 ± 0.46 |
| PC ae C40:5 | 2.74 ± 1.28 | 2.62 ± 0.73 | 2.99 ± 0.72 |
| PC ae C40:6 | 3.24 ± 1.33 | 2.94 ± 0.98 | 3.87 ± 1.24 |
| PC ae C42:1 | 0.352 ± 1.32 | 0.32 ± 0.09 | 0.43 ± 0.14 |
| PC ae C42:2 | 0.407 ± 1.31 | 0.36 ± 0.10 | 0.50 ± 0.14 |
| PC ae C42:3 | 0.645 ± 1.3 | 0.59 ± 0.15 | 0.73 ± 0.16 |
| PC ae C42:4 | 0.697 ± 1.34 | 0.70 ± 0.21 | 0.75 ± 0.19 |
| PC ae C42:5 | 1.59 ± 1.29 | 1.60 ± 0.46 | 1.67 ± 0.39 |
| PC ae C44:4 | 0.273 ± 1.33 | 0.29 ± 0.08 | 0.29 ± 0.09 |
| PC ae C44:5 | 1.21 ± 1.37 | 1.24 ± 0.44 | 1.26 ± 0.34 |
| PC ae C44:6 | 0.989 ± 1.36 | 1.02 ± 0.33 | 1.01 ± 0.27 |
| Hex2Cer(d18:1/16:0) | 2.84 ± 1.28 | 2.60 ± 0.58 | 3.18 ± 0.60 |
| Hex2Cer(d18:1/18:0) | 0.218 ± 1.4 | 0.20 ± 0.07 | 0.27 ± 0.09 |
| Hex2Cer(d18:1/20:0) | 0.117 ± 1.49 | 0.12 ± 0.09 | 0.16 ± 0.08 |
| Hex2Cer(d18:1/22:0) | 0.309 ± 1.48 | 0.30 ± 0.26 | 0.43 ± 0.20 |
| Hex2Cer(d18:1/24:0) | 0.257 ± 1.5 | 0.25 ± 0.15 | 0.34 ± 0.13 |
| Hex2Cer(d18:1/24:1) | 0.436 ± 1.36 | 0.41 ± 0.14 | 0.51 ± 0.13 |
| Hex3Cer(d18:1/16:0) | 1.39 ± 1.31 | 1.31 ± 0.35 | 1.57 ± 0.43 |
| Hex3Cer(d18:1/24:1) | 0.555 ± 1.44 | 0.52 ± 0.20 | 0.67 ± 0.23 |
| HexCer(d18:1/16:0) | 1.07 ± 1.3 | 0.92 ± 0.24 | 1.25 ± 0.28 |
| HexCer(d18:1/18:0) | 0.174 ± 1.43 | 0.15 ± 0.05 | 0.20 ± 0.06 |
| HexCer(d18:1/20:0) | 0.279 ± 1.37 | 0.25 ± 0.09 | 0.32 ± 0.09 |
| HexCer(d18:1/22:0) | 2.96 ± 1.34 | 2.64 ± 0.76 | 3.65 ± 0.94 |
| HexCer(d18:1/23:0) | 1.47 ± 1.34 | 1.26 ± 0.34 | 1.75 ± 0.50 |
| HexCer(d18:1/24:0) | 1.51 ± 1.36 | 1.35 ± 0.42 | 1.83 ± 0.48 |
| HexCer(d18:1/24:1) | 3.45 ± 1.36 | 3.10 ± 0.97 | 4.01 ± 1.06 |
| HexCer(d18:2/22:0) | 0.6 ± 1.39 | 0.56 ± 0.22 | 0.72 ± 0.31 |
| HexCer(d18:2/23:0) | 0.283 ± 1.4 | 0.25 ± 0.09 | 0.31 ± 0.16 |
| HexCer(d18:2/24:0) | 0.83 ± 1.42 | 0.75 ± 0.29 | 1.02 ± 0.35 |
| Cortisol | 0.244 ± 1.5 | 0.27 ± 0.14 | 0.28 ± 0.08 |
| DHEAS | 2.42 ± 1.93 | 3.50 ± 2.02 | 2.13 ± 1.17 |
| 3-IAA | 2.1 ± 2.22 | 2.99 ± 4.50 | 3.10 ± 3.98 |
| 3-IPA | 0.918 ± 3.11 | 1.76 ± 3.76 | 1.13 ± 1.26 |
| Ind-SO4 | 3.87 ± 1.73 | 4.15 ± 2.30 | 4.68 ± 2.42 |
| Hypoxanthine | 0.0473 ± 7.91 | 0.49 ± 0.86 | 0.27 ± 0.62 |
| SM (OH) C14:1 | 4.3 ± 1.44 | 3.33 ± 1.04 | 5.66 ± 1.95 |
| SM (OH) C16:1 | 2.24 ± 1.46 | 1.83 ± 0.67 | 2.91 ± 0.97 |
| SM (OH) C22:1 | 9.1 ± 1.44 | 6.90 ± 2.60 | 12.71 ± 4.14 |
| SM (OH) C22:2 | 6.82 ± 1.42 | 5.51 ± 1.72 | 8.48 ± 3.13 |
| SM (OH) C24:1 | 0.82 ± 1.42 | 0.67 ± 0.23 | 1.12 ± 0.37 |
| SM C16:0 | 79.9 ± 1.38 | 66.87 ± 22.16 | 98.79 ± 26.54 |
| SM C16:1 | 11.6 ± 1.43 | 9.93 ± 3.60 | 14.63 ± 5.24 |
| SM C18:0 | 13.8 ± 1.48 | 11.08 ± 4.31 | 18.28 ± 6.31 |
| SM C18:1 | 6.67 ± 1.5 | 5.71 ± 2.37 | 8.43 ± 3.25 |
| SM C20:2 | 0.238 ± 1.61 | 0.22 ± 0.10 | 0.28 ± 0.10 |
| SM C24:0 | 16 ± 1.37 | 13.07 ± 4.03 | 21.70 ± 4.76 |
| SM C24:1 | 31.4 ± 1.37 | 26.44 ± 8.25 | 37.29 ± 8.97 |
| SM C26:0 | 0.108 ± 1.55 | 0.10 ± 0.04 | 0.15 ± 0.06 |

| | | | |
|---|---|---|---|
| SM C26:1 | 0.217 ± 1.55 | 0.19 ± 0.09 | 0.30 ± 0.13 |
| H1 | 5440 ± 1.3 | 5630.83 ± 1520.16 | 5950.86 ± 3239.90 |
| TG(14:0_32:2) | 1.06 ± 2.21 | 1.04 ± 1.60 | 2.49 ± 2.33 |
| TG(14:0_34:0) | 1.25 ± 2.16 | 1.27 ± 1.59 | 3.29 ± 3.57 |
| TG(14:0_34:1) | 9.45 ± 2.37 | 10.39 ± 14.98 | 25.45 ± 23.16 |
| TG(14:0_34:2) | 7.98 ± 2.15 | 7.90 ± 9.76 | 18.45 ± 15.03 |
| TG(14:0_34:3) | 1.93 ± 2.07 | 1.86 ± 2.14 | 4.05 ± 3.08 |
| TG(14:0_36:1) | 2.56 ± 1.97 | 2.54 ± 3.32 | 5.29 ± 3.77 |
| TG(14:0_36:2) | 15.6 ± 1.89 | 15.25 ± 16.36 | 27.12 ± 13.78 |
| TG(14:0_36:3) | 15.6 ± 1.79 | 14.68 ± 13.02 | 24.85 ± 12.30 |
| TG(14:0_36:4) | 5.61 ± 1.83 | 5.35 ± 4.48 | 8.85 ± 4.84 |
| TG(16:0_28:1) | 0.648 ± 2.43 | 0.79 ± 1.52 | 2.11 ± 3.12 |
| TG(16:0_28:2) | 0.376 ± 2.02 | 0.38 ± 0.40 | 0.89 ± 1.12 |
| TG(16:0_30:2) | 1.07 ± 2.34 | 1.13 ± 1.80 | 2.59 ± 2.57 |
| TG(16:0_32:0) | 8.21 ± 2.95 | 12.94 ± 17.30 | 28.98 ± 43.48 |
| TG(16:0_32:1) | 15.8 ± 2.45 | 17.46 ± 25.30 | 49.97 ± 55.44 |
| TG(16:0_32:2) | 8.45 ± 2.25 | 8.61 ± 11.34 | 22.21 ± 21.47 |
| TG(16:0_32:3) | 1.53 ± 2.17 | 1.61 ± 2.08 | 3.97 ± 4.06 |
| TG(16:0_33:1) | 2.27 ± 2.07 | 2.34 ± 2.80 | 5.56 ± 5.74 |
| TG(16:0_33:2) | 1.36 ± 1.96 | 1.37 ± 1.38 | 2.82 ± 2.49 |
| TG(16:0_34:0) | 9.25 ± 2.25 | 10.61 ± 10.74 | 23.74 ± 28.56 |
| TG(16:0_34:1) | 73.6 ± 2.2 | 83.58 ± 82.10 | 165.47 ± 164.82 |
| TG(16:0_34:2) | 75.9 ± 1.93 | 73.34 ± 59.62 | 155.43 ± 134.37 |
| TG(16:0_34:3) | 21.8 ± 1.84 | 20.94 ± 16.99 | 42.99 ± 35.24 |
| TG(16:0_34:4) | 2.04 ± 1.92 | 2.00 ± 1.91 | 4.40 ± 4.13 |
| TG(16:0_35:1) | 1.96 ± 1.99 | 1.98 ± 2.24 | 4.26 ± 3.84 |
| TG(16:0_35:2) | 3.28 ± 1.91 | 3.41 ± 3.89 | 6.62 ± 5.14 |
| TG(16:0_35:3) | 1.65 ± 1.78 | 1.65 ± 1.49 | 2.85 ± 1.95 |
| TG(16:0_36:2) | 174 ± 1.69 | 164.71 ± 128.06 | 281.56 ± 141.03 |
| TG(16:0_36:3) | 204 ± 1.59 | 186.98 ± 114.47 | 301.51 ± 132.55 |
| TG(16:0_36:4) | 74.7 ± 1.64 | 69.56 ± 41.21 | 108.44 ± 51.98 |
| TG(16:0_36:5) | 9.02 ± 1.7 | 8.46 ± 5.96 | 14.89 ± 9.65 |
| TG(16:0_37:3) | 0.615 ± 1.75 | 0.60 ± 0.45 | 0.94 ± 0.53 |
| TG(16:0_38:1) | 0.53 ± 1.88 | 0.52 ± 0.44 | 0.88 ± 0.60 |
| TG(16:0_38:2) | 1.82 ± 1.73 | 1.83 ± 1.45 | 2.81 ± 1.44 |
| TG(16:0_38:3) | 3.57 ± 1.68 | 3.63 ± 2.63 | 5.32 ± 2.52 |
| TG(16:0_38:4) | 4.73 ± 1.69 | 4.66 ± 3.55 | 7.58 ± 4.20 |
| TG(16:0_38:5) | 4.89 ± 1.67 | 4.68 ± 3.61 | 8.52 ± 5.88 |
| TG(16:0_38:6) | 2.75 ± 1.7 | 2.45 ± 1.74 | 4.96 ± 3.60 |
| TG(16:0_40:6) | 2.86 ± 1.64 | 2.69 ± 2.07 | 4.48 ± 2.45 |
| TG(16:0_40:7) | 2.61 ± 1.66 | 2.28 ± 1.50 | 4.06 ± 1.92 |
| TG(16:1_30:1) | 0.828 ± 2.33 | 0.96 ± 1.78 | 2.20 ± 2.03 |
| TG(16:1_32:0) | 2.87 ± 2.53 | 3.16 ± 4.15 | 11.06 ± 15.20 |
| TG(16:1_32:1) | 5.03 ± 2.41 | 5.39 ± 7.12 | 14.08 ± 13.76 |
| TG(16:1_32:2) | 2.12 ± 2.13 | 2.06 ± 2.34 | 4.46 ± 3.44 |
| TG(16:1_33:1) | 0.788 ± 2 | 0.81 ± 0.88 | 1.61 ± 1.31 |
| TG(16:1_34:0) | 3.32 ± 2.05 | 3.36 ± 3.95 | 8.20 ± 8.37 |
| TG(16:1_34:1) | 28.2 ± 2 | 28.08 ± 30.74 | 62.33 ± 52.20 |
| TG(16:1_34:2) | 21.3 ± 1.88 | 20.28 ± 17.48 | 40.63 ± 29.53 |
| TG(16:1_34:3) | 4.54 ± 1.89 | 4.32 ± 3.39 | 8.08 ± 5.50 |
| TG(16:1_36:1) | 3.95 ± 1.73 | 3.65 ± 3.27 | 6.87 ± 3.95 |
| TG(16:1_36:2) | 24.3 ± 1.68 | 22.52 ± 17.60 | 36.86 ± 16.54 |
| TG(16:1_36:3) | 26.7 ± 1.64 | 24.34 ± 15.10 | 37.46 ± 17.04 |
| TG(16:1_36:4) | 9.55 ± 1.69 | 8.83 ± 5.32 | 13.19 ± 6.42 |
| TG(16:1_36:5) | 1.55 ± 1.72 | 1.51 ± 1.02 | 2.34 ± 1.41 |
| TG(16:1_38:4) | 0.789 ± 1.71 | 0.79 ± 0.66 | 1.22 ± 0.62 |

| | | | |
|---|---|---|---|
| TG(16:1_38:5) | 0.771 ± 1.71 | 0.74 ± 0.56 | 1.34 ± 0.82 |
| TG(17:0_34:1) | 1.77 ± 1.78 | 1.81 ± 1.76 | 3.34 ± 2.61 |
| TG(17:0_34:2) | 1.57 ± 1.64 | 1.46 ± 1.11 | 2.68 ± 1.75 |
| TG(17:0_36:3) | 3.32 ± 1.6 | 3.03 ± 1.96 | 4.75 ± 2.07 |
| TG(17:0_36:4) | 1.41 ± 1.57 | 1.29 ± 0.69 | 1.95 ± 0.82 |
| TG(17:1_32:1) | 0.657 ± 2.02 | 0.72 ± 0.82 | 1.39 ± 1.11 |
| TG(17:1_34:1) | 2.43 ± 1.89 | 2.57 ± 2.85 | 4.78 ± 3.22 |
| TG(17:1_34:2) | 1.7 ± 1.78 | 1.69 ± 1.60 | 3.02 ± 2.06 |
| TG(17:1_36:3) | 1.58 ± 1.61 | 1.47 ± 1.05 | 2.30 ± 0.95 |
| TG(17:1_36:4) | 0.704 ± 1.61 | 0.69 ± 0.43 | 0.98 ± 0.45 |
| TG(17:2_36:4) | 0.508 ± 1.7 | 0.47 ± 0.32 | 0.79 ± 0.47 |
| TG(17:2_38:6) | 0.287 ± 1.62 | 0.28 ± 0.13 | 0.38 ± 0.20 |
| TG(18:0_30:0) | 0.54 ± 2.52 | 0.63 ± 0.95 | 1.81 ± 2.53 |
| TG(18:0_30:1) | 0.399 ± 2.27 | 0.45 ± 0.89 | 1.02 ± 1.10 |
| TG(18:0_32:0) | 1.75 ± 2.4 | 2.14 ± 2.63 | 5.80 ± 8.92 |
| TG(18:0_32:1) | 1.79 ± 2.34 | 1.95 ± 2.84 | 5.72 ± 6.57 |
| TG(18:0_32:2) | 0.883 ± 2.03 | 0.90 ± 1.12 | 2.04 ± 1.86 |
| TG(18:0_34:2) | 6.79 ± 1.95 | 6.74 ± 6.33 | 14.67 ± 13.78 |
| TG(18:0_34:3) | 1.66 ± 1.75 | 1.55 ± 1.30 | 3.12 ± 2.29 |
| TG(18:0_36:2) | 10.7 ± 1.67 | 10.16 ± 7.24 | 16.55 ± 6.99 |
| TG(18:0_36:3) | 13 ± 1.59 | 11.73 ± 6.66 | 19.02 ± 7.94 |
| TG(18:0_36:4) | 6.25 ± 1.62 | 5.77 ± 3.39 | 8.97 ± 3.89 |
| TG(18:0_36:5) | 0.918 ± 1.56 | 0.80 ± 0.45 | 1.36 ± 0.66 |
| TG(18:1_26:0) | 0.593 ± 2.13 | 0.73 ± 1.49 | 1.49 ± 2.20 |
| TG(18:1_30:0) | 11.3 ± 2.34 | 12.70 ± 19.56 | 30.22 ± 27.05 |
| TG(18:1_30:1) | 7.65 ± 2.26 | 8.35 ± 13.62 | 16.98 ± 11.55 |
| TG(18:1_30:2) | 2.15 ± 2.02 | 2.12 ± 2.65 | 3.83 ± 2.08 |
| TG(18:1_32:0) | 44.8 ± 2.11 | 48.19 ± 43.69 | 96.66 ± 86.81 |
| TG(18:1_32:1) | 60.2 ± 1.9 | 58.47 ± 62.89 | 118.25 ± 78.32 |
| TG(18:1_32:2) | 19.8 ± 1.79 | 18.42 ± 16.98 | 32.44 ± 16.28 |
| TG(18:1_32:3) | 2.2 ± 1.8 | 2.13 ± 2.21 | 3.72 ± 2.12 |
| TG(18:1_33:0) | 2.17 ± 1.91 | 2.24 ± 2.42 | 4.46 ± 3.73 |
| TG(18:1_33:1) | 7.31 ± 1.77 | 7.11 ± 6.71 | 12.35 ± 6.57 |
| TG(18:1_33:2) | 3.4 ± 1.65 | 3.20 ± 2.42 | 4.98 ± 2.29 |
| TG(18:1_34:1) | 304 ± 1.7 | 287.47 ± 224.52 | 492.11 ± 247.87 |
| TG(18:1_34:2) | 221 ± 1.6 | 199.96 ± 127.23 | 331.73 ± 145.49 |
| TG(18:1_34:3) | 37.4 ± 1.62 | 34.06 ± 22.14 | 55.44 ± 24.12 |
| TG(18:1_34:4) | 3.75 ± 1.66 | 3.54 ± 2.74 | 5.68 ± 2.92 |
| TG(18:1_35:2) | 5.1 ± 1.63 | 4.72 ± 3.73 | 7.47 ± 3.14 |
| TG(18:1_35:3) | 1.73 ± 1.62 | 1.55 ± 1.09 | 2.47 ± 1.03 |
| TG(18:1_36:0) | 2.39 ± 1.73 | 2.42 ± 1.86 | 3.90 ± 1.99 |
| TG(18:1_36:1) | 25.2 ± 1.67 | 24.17 ± 16.87 | 38.61 ± 15.60 |
| TG(18:1_36:2) | 42.9 ± 3.78 | 52.36 ± 50.12 | 91.31 ± 65.98 |
| TG(18:1_36:3) | 103 ± 1.6 | 97.26 ± 51.10 | 142.15 ± 63.45 |
| TG(18:1_36:4) | 38.3 ± 1.63 | 35.56 ± 19.10 | 52.87 ± 23.46 |
| TG(18:1_36:5) | 8.31 ± 1.68 | 7.85 ± 4.91 | 11.71 ± 5.22 |
| TG(18:1_38:5) | 5.76 ± 1.5 | 5.28 ± 3.32 | 8.32 ± 3.63 |
| TG(18:1_38:6) | 3 ± 1.51 | 2.59 ± 1.42 | 4.46 ± 1.93 |
| TG(18:1_38:7) | 0.707 ± 1.54 | 0.64 ± 0.30 | 1.02 ± 0.59 |
| TG(18:2_28:0) | 1.13 ± 2.29 | 1.21 ± 1.91 | 2.55 ± 2.50 |
| TG(18:2_30:0) | 6.42 ± 2.1 | 6.38 ± 7.26 | 14.71 ± 12.88 |
| TG(18:2_30:1) | 4.08 ± 2.05 | 3.96 ± 4.87 | 8.00 ± 5.34 |
| TG(18:2_32:0) | 27.6 ± 1.88 | 26.53 ± 19.13 | 53.87 ± 47.32 |
| TG(18:2_32:1) | 33.7 ± 1.76 | 30.94 ± 26.11 | 58.39 ± 35.93 |
| TG(18:2_32:2) | 10.5 ± 1.78 | 9.64 ± 7.38 | 15.98 ± 8.12 |
| TG(18:2_33:0) | 1.35 ± 1.76 | 1.31 ± 1.04 | 2.46 ± 2.00 |

| | | | |
|---|---|---|---|
| TG(18:2_33:1) | 4.18 ± 1.66 | 3.82 ± 2.87 | 6.33 ± 3.40 |
| TG(18:2_33:2) | 1.96 ± 1.65 | 1.80 ± 1.17 | 2.64 ± 1.17 |
| TG(18:2_34:0) | 22.2 ± 1.64 | 20.46 ± 14.02 | 35.50 ± 20.92 |
| TG(18:2_34:1) | 186 ± 1.59 | 169.71 ± 102.67 | 274.36 ± 122.50 |
| TG(18:2_34:2) | 123 ± 1.63 | 114.34 ± 64.99 | 173.67 ± 82.07 |
| TG(18:2_34:3) | 19.9 ± 1.69 | 18.42 ± 11.57 | 28.03 ± 13.62 |
| TG(18:2_34:4) | 1.95 ± 1.75 | 1.89 ± 1.24 | 2.87 ± 1.56 |
| TG(18:2_35:1) | 3.43 ± 1.64 | 3.13 ± 2.30 | 5.05 ± 2.30 |
| TG(18:2_35:2) | 2.99 ± 1.61 | 2.69 ± 1.70 | 4.17 ± 1.80 |
| TG(18:2_35:3) | 0.919 ± 1.66 | 0.87 ± 0.52 | 1.33 ± 0.68 |
| TG(18:2_36:0) | 1.76 ± 1.6 | 1.62 ± 1.03 | 2.75 ± 1.36 |
| TG(18:2_36:1) | 16.7 ± 1.56 | 15.07 ± 8.25 | 23.71 ± 9.15 |
| TG(18:2_36:2) | 58.3 ± 1.59 | 53.84 ± 27.57 | 79.39 ± 34.08 |
| TG(18:2_36:3) | 48.5 ± 1.68 | 45.72 ± 24.14 | 65.52 ± 33.33 |
| TG(18:2_36:4) | 19.4 ± 1.77 | 18.84 ± 12.42 | 26.72 ± 13.62 |
| TG(18:2_36:5) | 3.98 ± 1.9 | 3.97 ± 3.33 | 5.78 ± 3.22 |
| TG(18:2_38:4) | 2.73 ± 1.56 | 2.58 ± 1.42 | 3.60 ± 1.41 |
| TG(18:2_38:5) | 3.27 ± 1.54 | 2.93 ± 1.65 | 4.54 ± 2.01 |
| TG(18:2_38:6) | 1.83 ± 1.6 | 1.55 ± 0.79 | 2.66 ± 1.19 |
| TG(18:3_30:0) | 0.777 ± 2.1 | 0.84 ± 1.08 | 1.95 ± 1.95 |
| TG(18:3_32:0) | 2.84 ± 1.87 | 2.74 ± 2.44 | 6.39 ± 6.58 |
| TG(18:3_32:1) | 3.78 ± 1.89 | 3.58 ± 3.62 | 7.41 ± 6.16 |
| TG(18:3_34:0) | 2.56 ± 1.69 | 2.56 ± 1.88 | 4.26 ± 3.02 |
| TG(18:3_34:1) | 20.1 ± 1.64 | 18.51 ± 14.41 | 33.29 ± 20.88 |
| TG(18:3_34:2) | 13.1 ± 1.68 | 11.88 ± 8.36 | 21.17 ± 13.35 |
| TG(18:3_34:3) | 2.31 ± 1.77 | 2.18 ± 1.56 | 3.77 ± 2.57 |
| TG(18:3_36:1) | 2.05 ± 1.61 | 1.88 ± 1.36 | 3.15 ± 1.39 |
| TG(18:3_36:2) | 9.22 ± 1.65 | 8.56 ± 5.52 | 13.26 ± 5.76 |
| TG(18:3_36:3) | 8.81 ± 1.75 | 8.18 ± 5.33 | 12.38 ± 6.00 |
| TG(18:3_36:4) | 2.91 ± 1.86 | 2.78 ± 2.13 | 4.24 ± 2.32 |
| TG(20:0_32:3) | 0.544 ± 1.59 | 0.51 ± 0.30 | 0.81 ± 0.41 |
| TG(20:0_32:4) | 0.582 ± 1.58 | 0.56 ± 0.30 | 0.80 ± 0.41 |
| TG(20:1_32:1) | 0.474 ± 1.85 | 0.50 ± 0.49 | 0.80 ± 0.48 |
| TG(20:1_34:1) | 1.57 ± 1.75 | 1.62 ± 1.44 | 2.39 ± 1.17 |
| TG(20:1_34:2) | 1.24 ± 1.68 | 1.26 ± 0.87 | 1.86 ± 0.80 |
| TG(20:2_32:1) | 0.783 ± 1.76 | 0.82 ± 0.75 | 1.34 ± 0.82 |
| TG(20:2_34:1) | 2.34 ± 1.71 | 2.39 ± 1.90 | 3.47 ± 1.63 |
| TG(20:2_34:2) | 1.81 ± 1.69 | 1.82 ± 1.17 | 2.55 ± 1.13 |
| TG(20:2_34:3) | 0.431 ± 1.66 | 0.40 ± 0.25 | 0.57 ± 0.26 |
| TG(20:3_32:1) | 0.971 ± 1.93 | 1.01 ± 1.03 | 1.89 ± 1.39 |
| TG(20:3_34:1) | 4.17 ± 1.72 | 4.24 ± 3.75 | 6.75 ± 3.77 |
| TG(20:3_34:2) | 3.05 ± 1.68 | 2.91 ± 2.21 | 4.77 ± 2.67 |
| TG(20:3_34:3) | 0.625 ± 1.74 | 0.60 ± 0.46 | 0.94 ± 0.47 |
| TG(20:3_36:3) | 1.66 ± 1.59 | 1.63 ± 0.89 | 2.17 ± 0.97 |
| TG(20:3_36:4) | 0.705 ± 1.58 | 0.66 ± 0.33 | 0.95 ± 0.45 |
| TG(20:4_30:0) | 0.5 ± 2.18 | 0.60 ± 0.97 | 1.48 ± 1.82 |
| TG(20:4_32:0) | 1.72 ± 2.04 | 1.78 ± 1.84 | 4.40 ± 5.55 |
| TG(20:4_32:1) | 1.83 ± 2.01 | 1.84 ± 2.31 | 4.50 ± 4.71 |
| TG(20:4_32:2) | 0.66 ± 1.82 | 0.61 ± 0.58 | 1.19 ± 0.93 |
| TG(20:4_34:0) | 1.38 ± 1.72 | 1.30 ± 1.15 | 2.63 ± 2.13 |
| TG(20:4_34:1) | 10.1 ± 1.67 | 9.33 ± 7.79 | 18.06 ± 12.94 |
| TG(20:4_34:2) | 7.55 ± 1.64 | 6.93 ± 4.66 | 12.69 ± 9.51 |
| TG(20:4_34:3) | 1.34 ± 1.66 | 1.27 ± 0.82 | 2.28 ± 1.87 |
| TG(20:4_36:2) | 7.08 ± 1.48 | 6.26 ± 3.25 | 10.17 ± 4.57 |
| TG(20:4_36:3) | 5.12 ± 1.49 | 4.73 ± 2.32 | 7.18 ± 3.45 |
| TG(20:4_36:4) | 2.15 ± 1.57 | 2.05 ± 1.08 | 3.06 ± 1.90 |

| | | | |
|---|---|---|---|
| TG(20:5_34:1) | 1.86 ± 1.92 | 1.75 ± 1.73 | 3.93 ± 3.56 |
| TG(20:5_34:2) | 1.43 ± 1.86 | 1.30 ± 1.09 | 2.85 ± 2.40 |
| TG(20:5_36:3) | 1.12 ± 1.72 | 1.04 ± 0.64 | 1.77 ± 1.06 |
| TG(22:4_34:2) | 1.03 ± 1.72 | 1.03 ± 0.88 | 1.58 ± 0.88 |
| TG(22:5_32:1) | 0.759 ± 1.82 | 0.80 ± 0.91 | 1.40 ± 1.08 |
| TG(22:5_34:1) | 4.42 ± 1.62 | 4.11 ± 3.35 | 6.86 ± 3.67 |
| TG(22:5_34:2) | 3.45 ± 1.61 | 3.05 ± 2.01 | 5.09 ± 2.61 |
| TG(22:6_32:0) | 0.898 ± 2.13 | 0.97 ± 1.14 | 2.09 ± 1.99 |
| TG(22:6_32:1) | 1.05 ± 2.18 | 1.09 ± 1.27 | 2.40 ± 1.99 |
| TG(22:6_34:1) | 4.79 ± 1.96 | 4.62 ± 4.48 | 8.98 ± 5.69 |
| TG(22:6_34:2) | 3.68 ± 1.92 | 3.44 ± 2.86 | 6.50 ± 3.73 |
| TG(22:6_34:3) | 0.736 ± 2 | 0.70 ± 0.54 | 1.25 ± 0.83 |
| Choline | 4.69 ± 1.38 | 4.26 ± 1.44 | 6.16 ± 2.61 |
